# Origin and rapid evolution of minicircular and highly heteroplasmic mitogenome in the holoparasitic plant genus *Rhopalocnemis*

**DOI:** 10.1101/2025.10.16.682905

**Authors:** Yan Zhong, Runxian Yu, Chenyu Sun, Jeffrey P. Mower, M. Virginia Sanchez-Puerta, Ying Liu, Renchao Zhou

## Abstract

The holoparasitic plant *Rhopalocnemis phalloides* displays unique features in mitogenome organization, sequence heteroplasmy, DNA replication and gene transcription. To understand the origin and evolution of these unique features, we compared the mitogenomes of three *R. phalloides* individuals and one individual of a newly discovered congeneric species. These mitogenomes comprise dozens of minicircular chromosomes (∼2-8 kb), with fairly small mitogenome sizes between 121.1 and 147.5 kb. Each *R. phalloides* individual contains extremely conserved regions (CRs) on all chromosomes, yet these CRs vary significantly among individuals, suggesting rapid divergence in a concerted manner. In contrast, the congeneric species lacks such CRs. Although nearly identical gene and intron content, there is significant sequence divergence between the two species. Extremely high mitogenome heteroplasmy was observed in all three individuals of *R. phalloides* and all the variants of protein coding genes are transcribed, but few heteroplasmic variants are shared among the three individuals. No sequence heteroplasmy was detected in the congeneric species. PacBio sequencing revealed concatenated mitochondrial chromosomes in the two species, suggesting rolling circle replication of mitochondrial DNA. We infer that the origins of the CR and sequence heteroplasmy are later than the origin of all-minicircular chromosomes in *Rhopalocnemis*, and propose a plasmid incorporation model to explain the origin of the CR. The rapid intraspecific variation in mitogenome structure, sequence and heteroplasmy in *R. phalloides* may result from relaxed selective constraint. The striking heteroplasmy in the *R. phalloides* mitogenome can not be explained by mitochondrion-targeted DNA-RRR gene loss characterized in this study.

## Introduction

The mitogenomes of most flowering plants are typically assembled as a single circular molecule of approximately 200–400 kb bearing about 30-40 protein-coding genes (Mower, 2020). In stark contrast to this generally prevailing genomic organization, the available mitogenomes of some parasitic plants (Sanchez-Puerta *et al*., 2017; Sanchez-Puerta *et al*., 2019; Roulet *et al*., 2020; Yu *et al*., 2022; Zhou *et al*., 2023) and some autotrophic plants (Sloan *et al*., 2010; Alverson *et al*., 2011; Yang *et al*., 2023) are instead composed of many smaller circular-mapping chromosomes. In these cases, recombination between chromosomes is minimal or absent, maintaining their multi-chromosomal structure. The most extreme example is the monotypic genus *Rhopalocnemis* from the holoparasitic family Balanophoraceae (Yu *et al*., 2022). The published mitogenome consists of 21 minicircular chromosomes of 4-8 kb in length, with a shared region containing the replication origin, and replicates via a rolling circle mechanism. Each minicircular chromosome includes a highly conserved region (CR) of 896 bp and 0-4 genes in the unique region (UR). The CR is flanked on each side by hypervariable (HV) and semi-conserved (SC) regions.

The mitochondrial gene sequences of angiosperms are generally highly conserved, with only a few rapidly evolving groups such as *Silene* (Wu & Sloan, 2019), *Viscum* (Skippington et al., 2015), *Plantago* (Cho *et al*., 2004), *Acorus* (Guo *et al*., 2023) and *Pelargonium* (Choi *et al*., 2021). Although angiosperm mitogenomes can form alternative conformations by repeat-mediated recombination, there are often no or very low levels of sequence heteroplasmy except for plants with biparental mtDNA inheritance (Ramsey & Mandel, 2019; Zardoya, 2020). In striking contrast, the mitogenome of *Rhopalocnemis* exhibits unprecedentedly high sequence heteroplasmy in both intergenic and genic regions, and transcribes all heteroplasmic variants (Yu *et al*., 2022).

In addition to sequence heteroplasmy, the origins of multiple minicircular chromosomes and the CR in the mitogenome of *Rhopalocnemis* are some of the most radical evolutionary changes in mitogenome architecture thus far established in plants. This kind of genome structure has been observed in mitochondria of some parasitic metazoa and protists (Armstrong *et al*., 2000; Gibson *et al*., 2007; Jiang *et al*., 2009; Shao *et al*., 2009; Dong *et al*., 2014; Yahalomi *et al*., 2017; Song *et al*., 2019) and plastids of free-living dinoflagellates (Zhang *et al*., 1999; Barbrook & Howe, 2000), providing an exceptional example of convergent evolution of organellar genome structure across diverse lineages of eukaryotes (Yu *et al*., 2022).

*Rhopalocnemis*, as traditionally defined, comprises a single species, *R. phalloides* Jungh., which is widely distributed across the East Himalayas, China, Vietnam, Cambodia, Malaysia and Indonesia (Junghuhn, 1841; Fagerl, 1938; Hansen, 1972). To better understand the origin and evolution of the exceptional genomic architecture of *Rhopalocnemis*, we sampled four individuals from different geographic locations and compared their mitogenomes. Our investigation indicated that one individual was genetically and morphologically distinct, leading us to suggest that it may be a distinct congeneric species. In this study, we aimed to address the following questions: 1) What is the pattern of intraspecific variation in *R. phalloides* with respect to mitogenomic organization and sequences? 2) Is the exceptional mitogenomic organization, sequence heteroplasmy and replication observed in *R. phalloides* shared with its congener? and 3) When and how did the exceptional genomic organization observed in *R. phalloides* arise?

## Results

### Phylogenetic analysis strongly supports the separation of two *Rhopalocnemis* species

The individual sequenced by Yu *et al*. (2022), collected from Daweishan (Pingbian County, Yunnan, China), was identified as *R. phalloides*. However, during a recent field survey in Malipo (Yunnan, China), we discovered a population of *Rhopalocnemis* that differs markedly from the Daweishan individual in both scale morphology and flowering season. Specifically, the Malipo population forms a wart-like structure in the central part of the peltate scale, as opposed to a recurved scale-like appendage (Figure S1), and flowers from August to September rather than May to June. Additionally, there is substantial divergence in mitogenome structure and gene sequences between them (see below). The morphological and genetic differences between *Rhopalocnemis* plants from Daweishan and Malipo suggest that they may represent congeneric but distinct species. However, determining which population is conspecific with *R. phalloides* from its type locality remains unresolved due to the lack of in-depth species delimitation studies. For clarity in this study, we refer to the Daweishan plants as *R. phalloides* following Yu *et al*. (2022) and label the Malipo plants as a congeneric species.

The three individuals of *Rhopalocnemis phalloides* and one individual of the congeneric new species are coded as DWS, WS, VIET and MLP, respectively (Table S1). The maximum likelihood trees are constructed for the four individuals of *Rhopalocnemis* and eight other selected genera of Balanophoraceae based on the concatenated sequences of two nuclear genes (18S and 28S rRNA genes) and the concatenated sequences of three available mitochondrial genes (*cox3*, *matR* and *rrnS*), respectively, with *Malania* from Olacaceae as an outgroup. As shown in Figure 1, the three individuals of *R. phalloides* form a monophyletic clade sister to the congeneric species (MLP) in both nuclear and mitochondrial trees with high bootstrap support. Within *R. phalloides*, the two geographically proximal individuals, DWS and WS, have the closest relationship. Only the position of *Sarcophyte* was discordant between nuclear and mitochondrial gene trees: while it is sister to *Balanophora* + *Langsdorffia* + *Thonningia* in the mitochondrial gene tree, it is sister to other Balanophoraceae genera except *Balanophora* + *Langsdorffia* + *Thonningia* in the nuclear gene tree. The mito-nuclear discordance may be caused by incomplete lineage sorting, ancient hybridization, or may be a phylogenetic artifact due to limited taxon sampling. The results are largely consistent with two recent phylogenetic studies, one involving nine Balanophoraceae genera based on concatenated nuclear sequences (18S, 28S and ITS1-5.8S-ITS2 rRNA operon) and a mitochondrial gene (*matR*) (Sanchez-Puerta *et al*., 2023), and the other involving six Balanophoraceae genera based on the concatenated 22-plastid gene matrix (Kim *et al*., 2023). The only topological conflict between our result and the two studies is also observed for *Sarcophyte*, which is sister to all other Balanophoraceae genera in the two studies. However, the nodes containing *Sarcophyte* are not well supported in all studies. Therefore, more nuclear and mitochondrial genes may be needed to determine the phylogenetic position of *Sarcophyte*.

**Figure 1.**
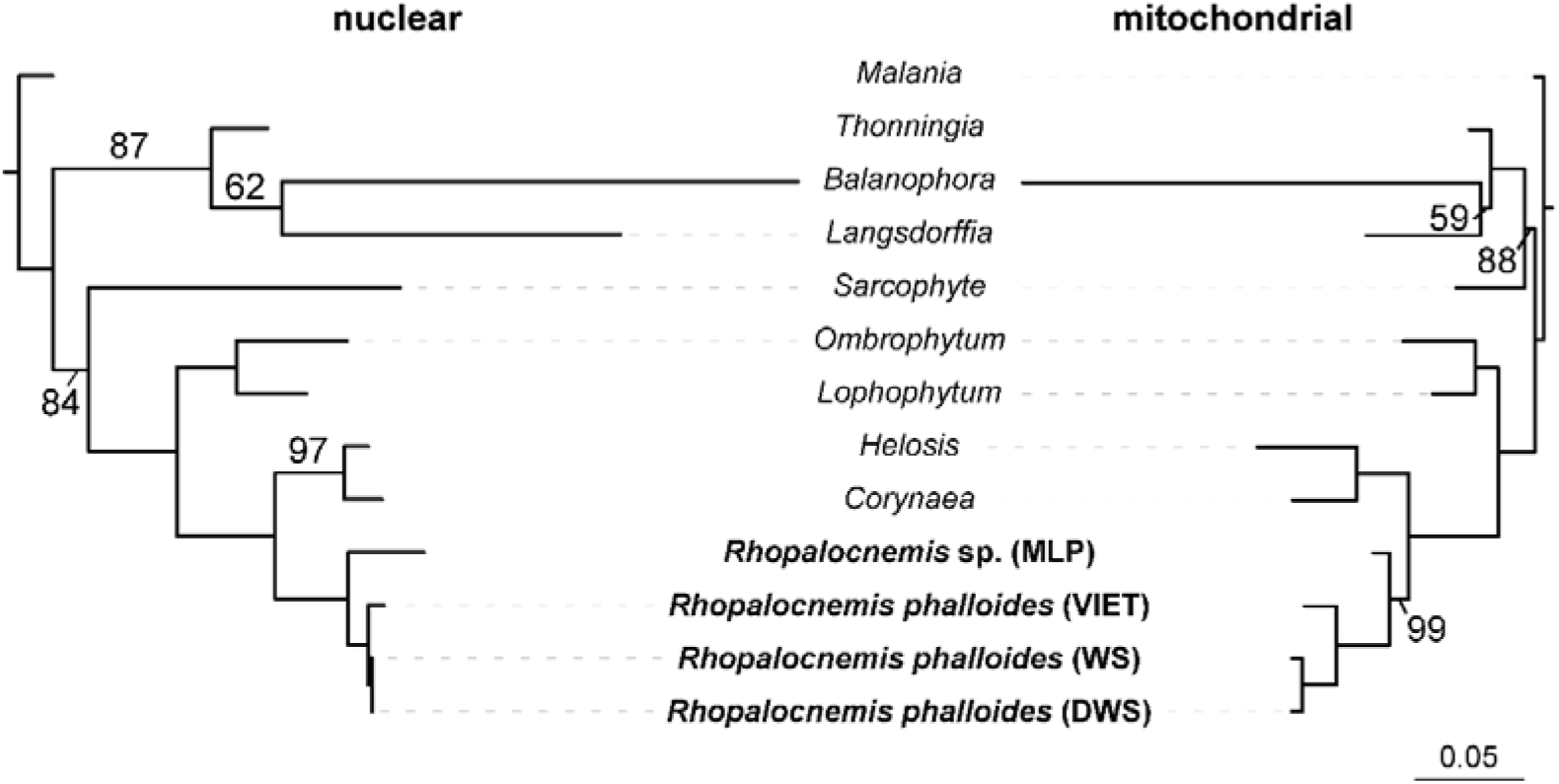
Maximum likelihood trees of *Rhopalocnemis* and eight other genera of Balanophoraceae based on concatenated sequences of two nuclear genes (18S rRNA and 28S rRNA, left) or three mitochondrial genes (*cox3*, *matR* and *rrnS*, right), respectively. *Malania oleifera* is used as an outgroup. The unlabeled branches have a bootstrap support value of 100. The scale below represents nucleotide substitutions per site.

The assembled plastomes of WS and MLP are 18,970 bp and 18,347 bp in size, respectively, which are close to the plastome sizes of DWS (19,137 bp, Yu *et al*., 2022) and VIET (18,622 bp, Schelkunov *et al*., 2019). All four plastomes of *Rhopalocnemis* exhibit exceptionally high AT content (84.2-86.8%) and contain 15 genes (Table S2). Pairwise comparisons of the protein-coding genes reveal the highest sequence identity (92.8% to 99.2%) between DWS and WS, lower identity (80.2%-95.2%) between VIET and DWS/WS and the lowest identity (64.2%-82.5%) between MLP and three other individuals (Figure S2), also indicating that MLP represents a distant taxonomic entity.

### Rapid interspecific and intraspecific divergence in mitogenome structure

Like the mitogenome of DWS characterized before (Yu *et al*., 2022), the mitogenomes of WS, VIET and MLP also comprise dozens of circular-mapping mini-chromosomes, with their chromosome numbers being 20, 18 and 39, respectively. The mitogenome sizes of WS (121,068 bp) and VIET (129,504 bp) are comparable to that of DWS (130,713 bp), while MLP has a larger size of 147,501 bp. The chromosome sizes of WS, VIET and MLP vary from 4,817 to 7,651 bp, 6,230 to 8,456 bp, and 2,480 to 5,853 bp, respectively (Table 1; Table S3-S4). The average chromosome size of MLP (3,782 bp) is the smallest. Overall GC contents of the four mitogenomes (44.7%, 44.7%, 45.0% and 46.3%) are similar and comparable to those of most other land plants.

**Table 1.** Features of *Rhopalocnemis* mitogenomes.

| Sample code | DWS<sup>a</sup> | WS | VIET | MLP |
| --- | --- | --- | --- | --- |
| Genome size (bp) | 130,713 | 121,068 | 129,504 | 147,501 |
| Chromosome number | 21 | 20 | 18 | 39 |
| Chromosome size range (bp) | 4,949 ~7,861 | 4,817~7,651 | 6,230~8,456 | 2,480~5,853 |
| Average chromosome size (bp) | 6,224.4 | 6,053.4 | 7,194.7 | 3,782.1 |
| Protein-coding genes<sup>b</sup> | 36(38) | 36(37) | 36(36) | 36(36) |
| tRNA genes<sup>b</sup> | 6(7) | 4(5) | 4(5) | 5(6) |
| rRNA genes | 3 | 3 | 3 | 3 |
| Group II introns<sup>b</sup> | 25(26) | 25(27) | 25 | 24(26) |
| *cis*-spliced<sup>b</sup> | 19(20) | 19(20) | 19 | 18 |
| *trans*-spliced<sup>b</sup> | 6 | 6(7) | 6 | 6(8) |
| Group I introns | 1 | 1 | 1 | 1 |
| GC content | 45.02% | 44.72% | 46.31% | 44.73% |
| Repeat content<sup>c</sup> | 10.81% | 11.21% | 5.20% | 10.51% |

<sup>a</sup> Adapted from Yu *et al.* 2022.
<sup>b</sup> Values in parentheses include duplicated copies.
<sup>c</sup> For DWS, WS and VIET, unique regions (URs) of the chromosomes were used for calculation.

The chromosome structure of WS, like that of DWS, features a conserved region (CR) of 792 bp across all chromosomes, plus two shorter hypervariable regions (HV1 and HV2) flanking the CR, two semi-conserved regions (SC1 and SC2) adjacent to HV1 and HV2, respectively, and a unique region (UR) between SC1 and SC2 (Figure 2A). The situation for VIET is slightly different in that its CR is divided into two parts [CR1 (537 bp) and CR2 (343 bp)] with one additional hypervariable region, HV3, separating the two parts (Figure 2A). In stark contrast, the MLP chromosomes have no such CR, HV, or SC regions; In other words, all 39 chromosomes of MLP contain only URs (Figure 2A).

**Figure 2.**
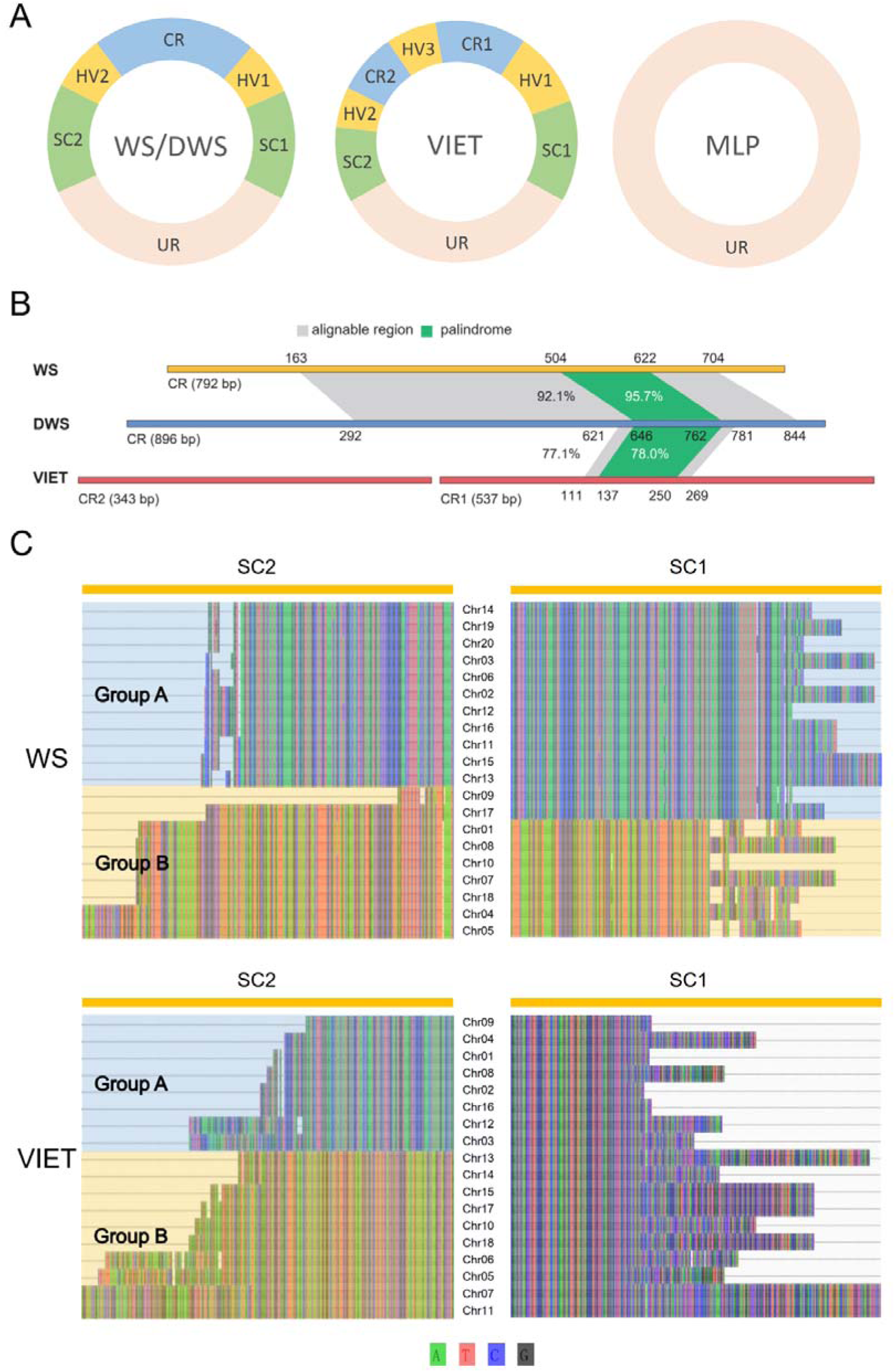
The mitochondrial chromosome structure of *Rhopalocnemis*. A. The schematic diagram of mitochondrial chromosome structure in three individuals of *R. phalloides* (DWS, WS and VIET) and one individual of the congeneric species (MLP). CR (conserved region), HV (hypervariable region), SC (semi-conserved region) and UR (unique region) are marked with different colors and connected clockwise. For better presentation, these regions are not shown in proportion to their sizes. B. Sequence divergence in the CR of the three individuals of *R. phalloides*. Grey and green indicate alignable regions and palindromic regions of the three individuals, respectively, and the corresponding nucleotide positions of these regions are shown. C. Alignments of SC1 and SC2 using the consensus sequences of all mitochondrial chromosomes of WS (above) and VIET (below).

The CRs of DWS and WS exhibit relatively high sequence similarity (92.1% identity for the 542 bp alignable sequence in the middle of the CR), whereas sequences at both ends of the CRs cannot be aligned with each other (Figure 2B). The CRs of DWS and WS have no obvious sequence similarity to, and cannot be aligned with, the CR2 of VIET. Only 159 bp sequence in the middle of the CR1 of VIET can be aligned with the CRs of DWS and WS, and the alignable region is less conserved (77.1% identity between DWS and VIET). Like the CR of DWS, the CR of WS and CR1 of VIET each have a palindrome (119 bp and 114 bp, respectively), which is the most conserved portion among the alignable regions of the three individuals and can fold into a stem-loop secondary structure (Figure S3). As suggested for DWS (Yu *et al*., 2022), these sequences with stem-loop secondary structure likely represent replication origins of the mitochondrial DNA of *R. phalloides*.

Similar to DWS, the HVs of WS and VIET comprise many short motifs of different copy numbers within and among chromosomes. Size variation of HVs among different chromosomes within each individual is mainly caused by copy number variation of these motifs, and there is no shared motif among HVs of the three individuals of *R. phalloides*.

SC regions (SC1: 179-322 bp long and SC2: 40-297 bp long) of WS can be divided into two sets of chromosomal groups (group A and group B, see Figure 2C) based on sequence similarity, as those of DWS. Two chromosomes (Chr09 and Chr17) appear to be a recombinant of the two groups. For VIET, while the two sets of chromosomal groups (group A and group B, see Figure 2C) can be applied to its SC2 (171–430 bp long), no further subdivision is required for its SC1 (233–644 bp long). Relatively high sequence similarity in SC1 and SC2 are found between DWS and WS, whereas nearly no sequence similarity is detected between VIET and either DWS or WS. Moreover, we found that while a large part of sequences of the same group (group A or group B) between SC1 and SC2 are identical or highly similar in both WS and DWS, those of the two groups (group A and group B) within either SC1 or SC2 show complete or nearly complete reverse complementation, a feature unnoticed in Yu *et al*. (2022). In VIET, a large part of sequences of SC1 and group A of SC2 show complete or nearly complete reverse complementation, while only a small fraction of sequences between group A and group B of SC2 are identical or highly similar (Figure S4).

### Few tRNA genes and no evidence of horizontal transfer of mitochondrial genes in *Rhopalocnemis*

Nearly all *R. phalloides* chromosomes of all three individuals carry between one and four genes per chromosome, and all mitochondrial genes are in the URs (Figure 3, Table S5). However, two chromosomes of DWS (Chr16 and Chr18), one of WS (Chr17) and eight of MLP (Chr5, Chr8, Chr12, Chr15, Chr17, Chr30, Chr33 and Chr35) lack any identifiable mitochondrial gene (Figure 3). Like the UR of Chr18 of DWS, the UR of Chr17 of WS contains fragments of DNA polymerase and RNA polymerase genes, which should be the remnant of an integrated mitochondrial plasmid (Yu *et al*., 2022).

**Figure 3.**
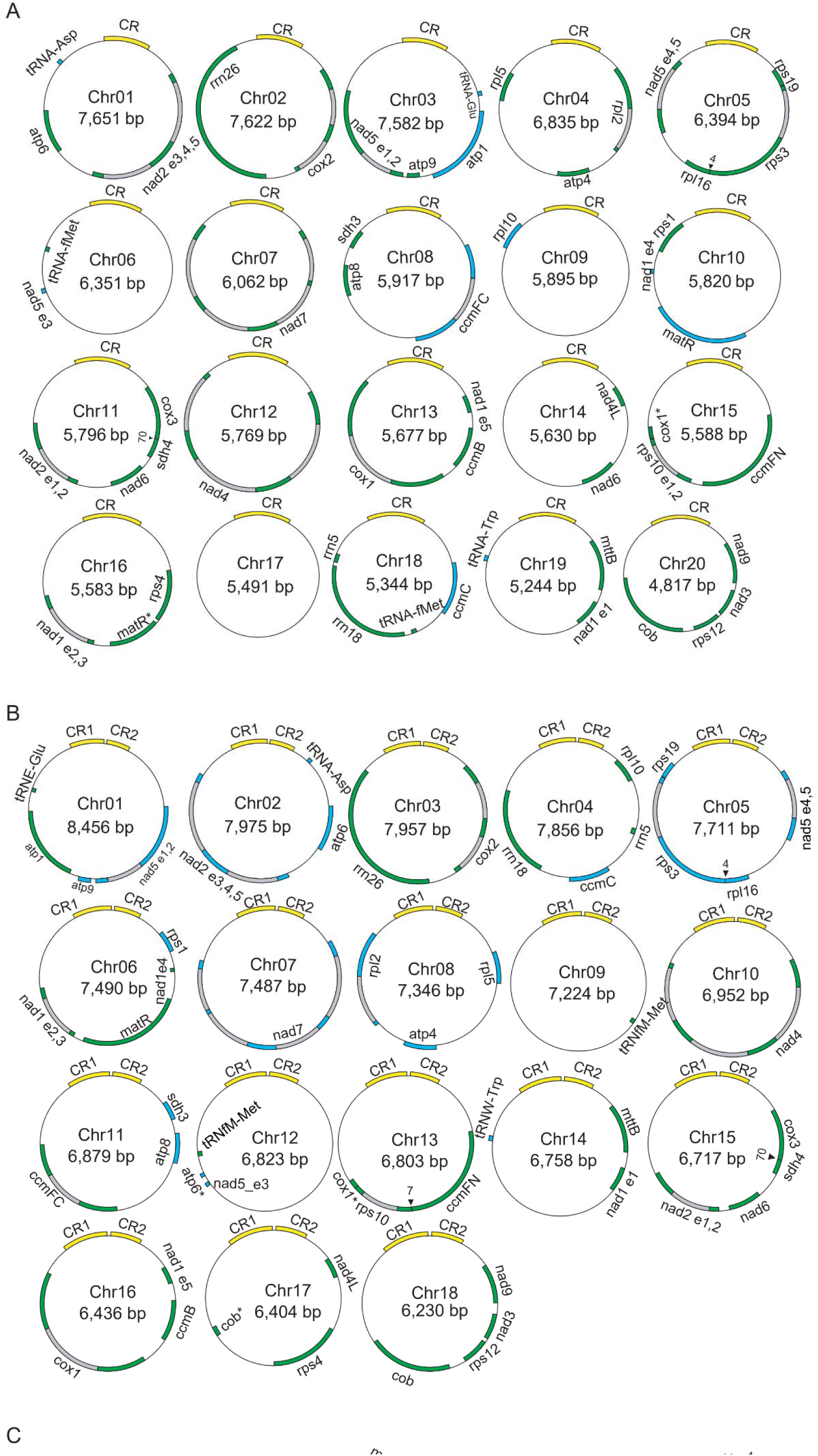
Minicircular mitochondrial chromosomes of *Rhopalocnemis*. The mitogenomes of the three individuals, WS (A), VIET (B) and MLP (C) consist of 20, 18, 39 chromosomes, respectively. When a gene contains *trans*-spliced intron(s), its exons are indicated with “e” followed by exon number. Gene fragments (>100 bp in length) are shown with an asterisk. Filled triangles show the positions of overlapping genes, with the length of overlap indicated.

The mitogenomes of all four individuals of *Rhopalocnemis* harbor a set of protein-coding genes and rRNAs typical of most angiosperms, containing 36 unique protein-coding genes and three rRNAs (Figure 3, Table S6). The only differences in protein-coding genes among the four individuals are one extra copy of *nad6* in the mitogenomes of DWS and WS, and one more copy of *rps10* in the mitogenome of DWS. All four *Rhopalocnemis* individuals have only four to six tRNAs, of which three tRNAs are shared among them (Table S6). These tRNA gene sets are among the smallest in angiosperm mitogenomes, compared with the highly reduced sets in *Viscum* and some species of *Silene* (Sloan *et al*., 2012; Skippington *et al*., 2015). For all three individuals of *R. phalloides*, bidirectional transcription is required for some chromosomes, while gene(s) on each chromosome of MLP are oriented on the same strand (not considering the *rps12* fragment on Chr36) suggesting unidirectional transcription (Figure 3).

All four individuals have 26 unique introns, including one group I (*cox1* intron) and 25 group II introns. The three individuals of *R. phalloides* have the same six *trans*-spliced introns (*nad1* i1, *nad1* i3, *nad1* i4, *nad2* i2, *nad5* i2 and *nad5* i3), while MLP also has an extra *trans*-spliced intron *rps3* i1. Compared with *rps3* of DWS, the shifting to *trans*-splicing of *rps3* i1 in MLP has stemmed from two inversions, one containing partial *rps3* i1 and the complete *rps3* e1, and the other containing the remaining *rps3* i1 and the complete *rps3* e2 (Figure S5). A shift from *cis*- to *trans*-splicing of this intron, which stems from one inversion, has also been found in *Tolypanthus maclurei* (Yu *et al*., 2021), a hemiparasitic species from the same order (Santalales), but a different family (Loranthaceae). For the shared group II *cis*-spliced introns, MLP has the largest average size (1,006.8 bp), while the three individuals of *R. phalloides* have similar average intron size (745.1-800.2 bp; Table S7). Each of the shared *cis*-spliced intron is also the largest in MLP. Based on the alignment of the shared *cis*-spliced introns of the four individuals, we found >30 bp deletions in almost all introns of *R. phalloides* compared with those of MLP (Table S8). Along with relatively large average intron sizes (1199.5-1371.4 bp) in other genera of Balanophoraceae (Zhou *et al*., 2023), this suggests that deletions contribute to smaller intron size of *R. phalloides*.

Phylogenetic analyses for each of these protein-coding genes for *Rhopalocnemis* and other selected angiosperms (Figure S6) showed that the four individuals of *Rhopalocnemis* clustered together for all genes with high bootstrap support. *Rhopalocnemis* is sister to *Ombrophytum* at 25 of 36 genes, mostly with strong bootstrap support. For the remaining 11 genes, *Rhopalocnemis* clustered either with species of Santalales or with non-Santalales species, but all with low bootstrap support (< 50%). Therefore, no convincing evidence for horizontal gene transfer was found for any protein-coding gene in the mitogenomes of *Rhopalocnemis*.

### Extensive intraspecific and interspecific chromosomal rearrangements

In general, the URs on most chromosomes are collinear among the three individuals of *R. phalloides*, especially between DWS and WS (Figure 4). Only one chromosome of DWS (Chr16), which lacks any identifiable mitochondrial gene, has no corresponding sequence in the UR of any chromosome of WS and VIET. One more chromosome of DWS (Chr18), which has also no identifiable mitochondrial gene, has no or nearly no corresponding sequence in the UR of any chromosome of VIET.

**Figure 4.**
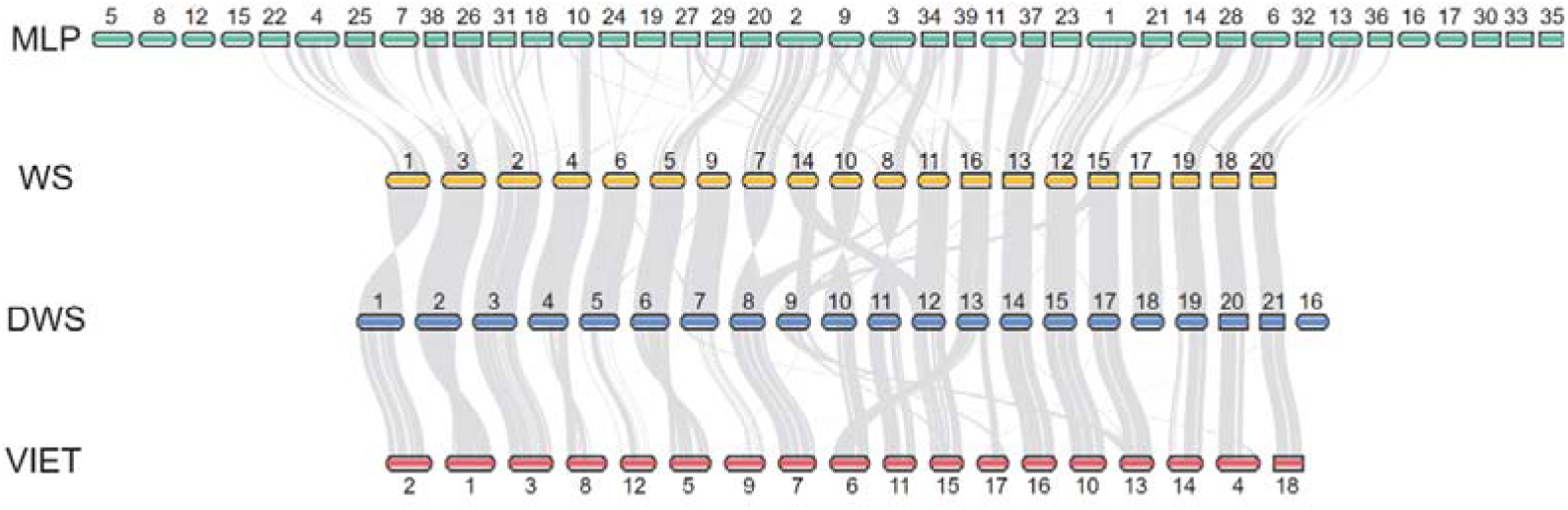
Mitogenome collinearity among three individuals of *Rhopalocenmis phalloides* (WS, DWS and VIET) and one individual of the congeneric species (MLP). For WS, DWS and VIET, unique regions (UR) on each chromosome were used for comparison.

Relative to the CR orientation, inversions have occurred in the URs of four chromosomes of WS (Chr01, Chr07, Chr08 and Chr10) and three chromosomes of VIET (Chr01, Chr05 and Chr08), when compared with the URs of Chr01, Chr08, Chr10, Chr11 of DWS and the URs of Chr02, Chr04, Chr06 of DWS, respectively. Chromosomal rearrangements have also occurred between DWS and WS, for example, one part of Chr09 of DWS corresponds to a region of Chr14 of WS and the other part of Chr09 of DWS matches with a region of Chr15 of WS. The other parts of Chr14 and Chr15 of WS match with a region of Chr12 and the whole UR of Chr17 of DWS, respectively. For some chromosomes of DWS, only partial region(s) have collinearity with the URs of VIET, and *vice versa*. For example, Chr13 of DWS corresponds to a region of Chr06 and a region of Chr17 of VIET. Chromosomal rearrangements between the two species are more extensive, as shown by multiple inversions and non-one-to-one correspondence for chromosomes between WS and MLP (Figure 4).

### Relaxed selection on mitochondrial protein genes in the *R. phalloides* lineage

In general, the sequence identity of these 36 protein-coding genes between DWS and WS is the highest (mean = 98.8%, Figure S7, Table S9), followed by those between VIET and WS/DWS (mean = 96.4%), while those between MLP and each of the three individuals of *R. phalloides* are the lowest (mean = 93.0%-93.4%, Figure S7, Table S9).

As shown in the tree of the concatenated 30 protein-coding genes shared by 12 species of Balanophoraceae (Figure 5), the ancestral branch of *Rhopalocnemis* has the largest values (*d_N_* = 3.06 × 10^-2^; *d_S_*= 8.30 × 10^-2^), which means that accelerated evolution has occurred in this genus relative to other genera. Note that these results are very conservative for *R. phalloides*, because we used the functional versions of these protein-coding genes of this species without considering the extreme sequence heteroplasmy which is largely individual-specific (see below).

**Figure 5.**
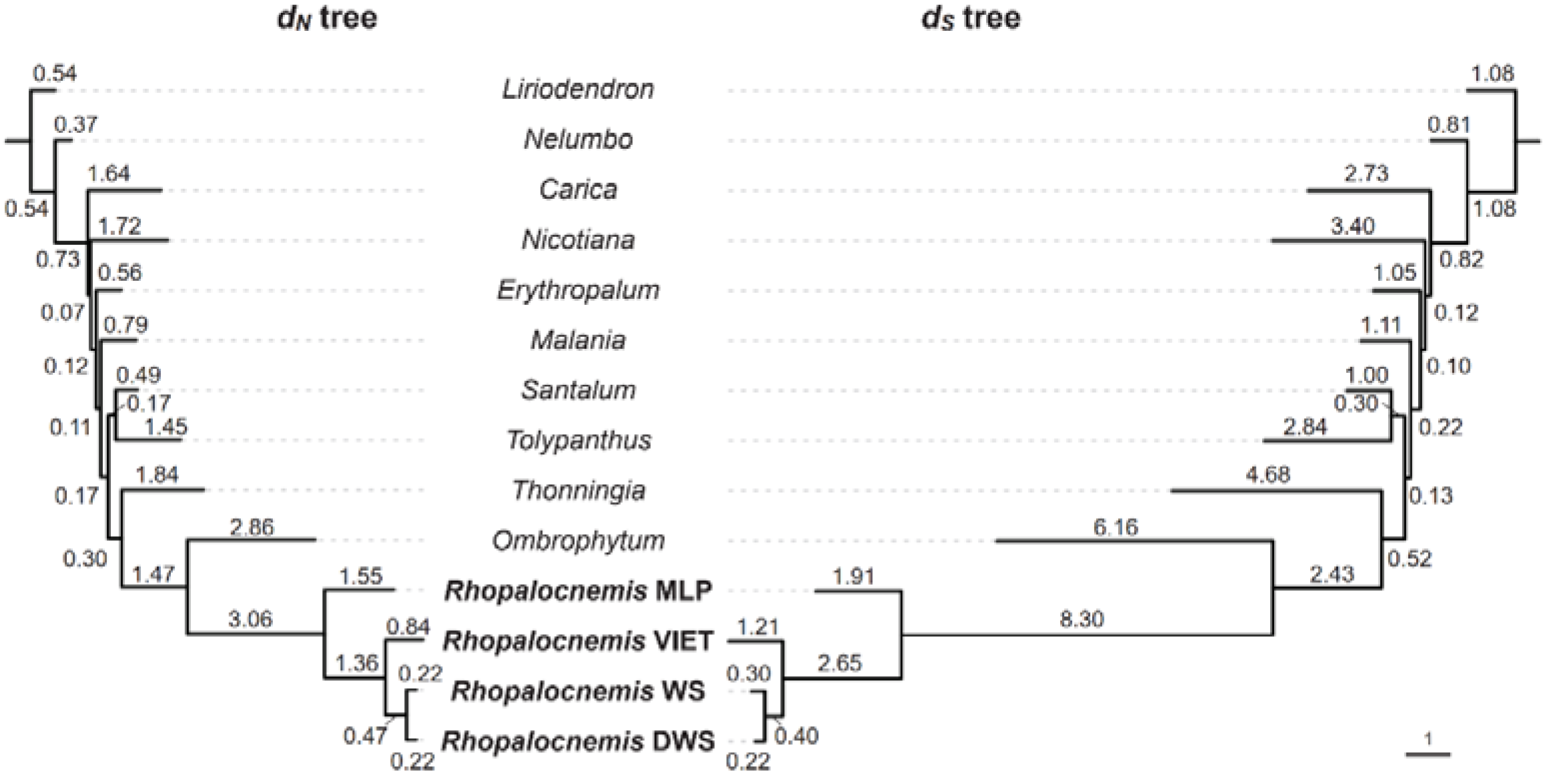
Variation in nucleotide substitution rate in *Rhopalocnemis* and 10 other angiosperms based on 30 shared, concatenated mitochondrial protein-coding genes. *d_N_* and *d_S_* represent nonsynonymous and synonymous nucleotide substitution rates, respectively. The numbers on the branches are *d_N_* × 100 (left panel) and *d_S_* × 100 (right panel). The topologically constrained tree is based on Nickrent (2020) and Figure 1 of this study.

The branches of *Rhopalocnemis* showed no significantly different *d_N_*/*d_S_* from other branches (0.52 vs 0.52, P = 0.95) by analyzing the shared, concatenated 30 mitochondrial protein-coding genes under the branch model. In contrast, the branches of *R. phalloides* have significantly larger *d_N_*/*d_S_* than other branches (0.67 vs 0.51, P = 3.26 × 10^-4^), suggesting that relaxed selection has occurred in the *R. phalloides* lineage after it diverges from its congener, and should contribute to its remarkable sequence heteroplasmy.

### Extremely high, individual-specific sequence heteroplasmy in *R. phalloides* but no heteroplasmy in its congener

Extremely high mitogenome sequence heteroplasmy has been observed in DWS and VIET (Yu *et al*., 2022). It is also the case for the mitogenomes of WS. A total of 446 variants were identified in the protein-coding genes of the three individuals of *R. phalloides* (Figure 6A; Table S10). The heteroplasmic variants in the protein-coding genes include 373 insertions/deletions, 51 complex variants (involving multi-nucleotide replacements) and 22 single nucleotide variants (SNVs), which exhibit varying frequencies from 0.05–0.83. Surprisingly, the majority of heteroplasmic variants are individual-specific (106 in DWS, 157 in WS and 146 in VIET) and only four heteroplasmic variants are shared by the three individuals of *R. phalloides* (Figure 6A). Pairwise comparison among the three individuals indicated that the number of shared heteroplasmic variants was the highest between WS and DWS (33), while that between DWS and VIET (7) and that between WS and VIET (5) were much lower (Figure 6A). Most heteroplasmic variants in the protein-coding regions can cause frameshifts and premature stop codons, resulting in abundant potentially non-functional variants. Heteroplasmic variants were also identified using RNA-seq data of WS and VIET, and all variants in these mitochondrial protein-coding genes were transcribed, consistent with the observation in DWS (Yu *et al*., 2022). The variant frequencies of protein-coding genes at the DNA level and the RNA level were highly correlated, with correlation coefficients of R^2^ = 0.816 (P < 0.001) and 0.704 (P < 0.001) in WS and VIET, respectively (Figure 6B and 6C; Table S11-S12). In contrast, no heteroplasmic variants were found in the whole mitogenome of MLP.

**Figure 6.**
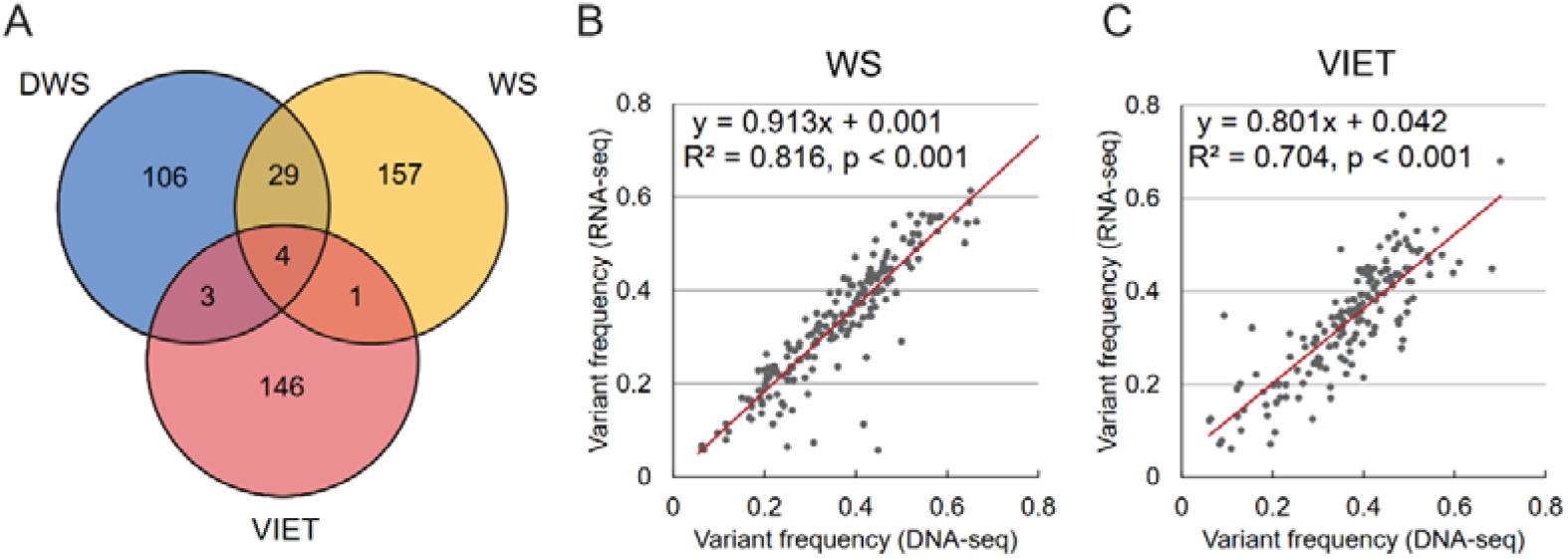
The heteroplasmic variants in mitochondrial protein-coding regions of *Rhopalocnemis phalloides*. A: A venn diagram showing shared and specific heteroplasmic variants in mitochondrial protein-coding genes of three individuals of *Rhopalocnemis phalloides* (WS, DWS and VIET). B: Correlation between variant frequencies in the DNA-seq and RNA-seq data for protein-coding genes in the mitogenome of WS. C: Correlation between variant frequencies in the DNA-seq and RNA-seq data for protein-coding genes in the mitogenome of VIET.

### Rolling circle replication and repeat-mediated recombination

Concatemers of multiple chromosome units have been observed in the mitogenome of DWS based on PacBio CLR sequencing (Yu *et al*., 2022). Here we examined PacBio CCS (HiFi) sequencing data of WS and MLP. Among the 224 mitochondrial PacBio HiFi reads for WS, 78 reads contain only one chromosome unit and the remaining 146 reads are concatemers composed of up to four intact chromosomal units. The 146 concatemers include 108 homo-concatemers (composed of the same chromosome unit) and 38 hetero-concatemers (composed of at least two different chromosome units). Among the 38 hetero-concatemers, four were reversed concatemers (Table S13; see Yu *et al*. (2022) for the definition of reversed concatemers). Among the 468 mitochondrial PacBio HiFi reads for MLP, 21 reads contain only one chromosome unit and the remaining 447 reads are concatemers composed of up to 10 intact chromosomal units in a head-to-tail orientation. These include 441 homo-concatemers and six hetero-concatemers (Table S13). Unlike the mitogenomes of DWS and WS, which have reversed concatemers, the mitogenome of MLP contains hetero-concatemers in only head-to-tail orientation.

We identified repeats in the URs of the mitogenomes of the three individuals of *R. phalloides* and the whole mitogenome of MLP. 63, 60 and 58 repeat units >= 30 bp, with the largest repeat units being 847, 812 and 746 bp, were detected among the URs of DWS, WS and VIET, respectively (Table S14). 113 repeat units >= 30 bp, were detected in the mitogenome of MLP, with the largest repeat unit being 1,171 bp (Table S14). Only one of the 38 hetero-concatemers from WS is produced by recombination mediated by a large repeat (Repeat_8 of WS) in the URs, suggesting that all other hetero-concatemers could result from recombination mediated by the CR or SC. Of the six hetero-concatemers of MLP, three are produced by Repeat_1-mediated recombination, while the other three are generated from recombination mediated by Repeat_2, Repeat_5 and Repeat_7 of MLP, respectively.

### Loss of mitochondrion-targeted DNA-RRR genes in *Rhopalocnemis*

The assembled transcriptomes of DWS, WS, VIET and MLP were evaluated by BUSCO based on eukarayota_odb10 database, and the gene completeness rates were 94.1%, 87.1%, 81.9% and 89.9%, respectively. These values are comparable to or even higher than those generated in other holoparasitic plants, such as 80.5% and 78.9% for *Cuscuta denticulata* and *C. nevadensis*, respectively (Frangione, 2020), and 90.8% for *Balanophora fungosa* (Schelkunov *et al*., 2021). Twenty-four genes involved in mitochondrial DNA replication, repair and recombination (DNA-RRR genes) were examined using the assembled transcriptomes of four individuals, and were also detected using PacBio long DNA reads (DWS, WS, MLP) or a draft genome of VIET assembled by Illumina reads. These genes had been examined in the closely related genus *Lophophytum* (Ceriotti *et al*., 2022). Two DNA-RRR genes, *PHR1* and *WHY2*, were not detected in both *Rhopalocnemis* and *Lophophytum* (Figure 7; Table S15). *FPG1* was not detected in all four individuals of *Rhopalocnemis* but is present in *Lophophytum*, and *OSB2-4* were not detected in *Lophophytum* but are present in all four individuals of *Rhopalocnemis* (Figure 7; Table S15). Although no orthologous transcript of *OSB1* was detected in VIET’s transcriptome (which has the lowest gene completeness rate based on the BUSCO assessment), we identified the intact *OSB1* gene from the draft genome assembled with Illumina reads. Some genes, such as *OSB1*, *ODB1*, *RADA*, *RECA2* and *RECA3*, were detected in the transcriptomes of some samples, but not in the PacBio HiFi reads (Table S15). This should result from giant genomes of *Rhopalocnemis* (>30 Gb, Schelkunov *et al*. 2021), which might not be covered by our limited HiFi reads.

**Figure 7.**
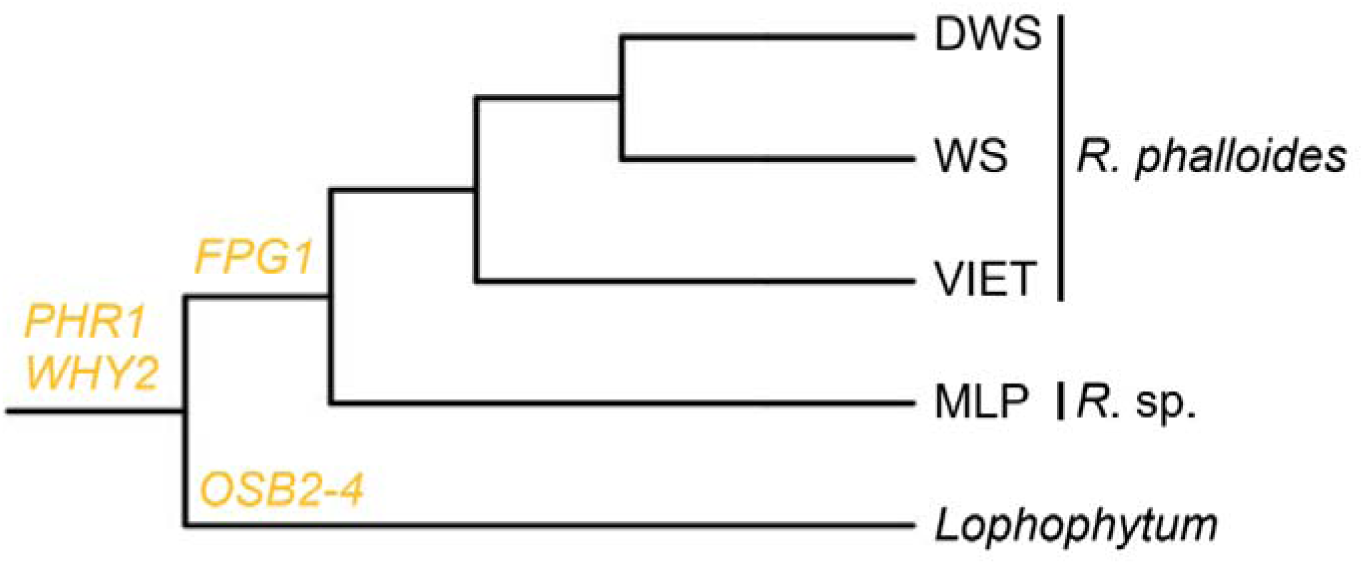
The loss of mitochondrion-targeted DNA-RRR genes in *Rhopalocnemis* and *Lophophytum.* The gene status of *Lophophytum* is from Ceriotti *et al*. (2022). The phylogenetic tree is based on Figure 1.

## Discussion

### The timing of the origin of all-minicircular chromosomes, the CR and sequence heteroplasmy in *Rhopalocnemis*

In this study, we assembled the mitogenomes from four individuals representing two species of *Rhopalocnemis*, and all mitochondrial chromosomes are minicircular, with a very small size of only 2.5-8.9 kb. Mitogenomes comprising all-minicircular chromosomes with such a small size (< 10 kb) have not been reported in other plants. Other than *Rhopalocnemis*, mitogenomes of three additional Balanophoraceae genera (*Lophophytum, Ombrophytum* and *Thonningia*) are also available (Sanchez-Puerta *et al*., 2017; Roulet *et al*., 2020; Sanchez-Puerta *et al*., 2023; Zhou *et al*., 2023), all of which show a highly multi-chromosomal structure, albeit with fairly different ranges of chromosomal size. The mitogenomes of *Thonningia sanguinea*, *Ombrophytum subterraneum* and *Lophophytum spp.* consist of 18 circular chromosomes of 8.7 to 19.2 kb (Zhou *et al*., 2023), 54 circular chromosomes of 4.9 to 27.2 kb (Roulet *et al*., 2020) and 54-81 circular chromosomes of 5.5 to 58.3 kb (Sanchez-Puerta *et al*., 2017; Roulet *et al*., 2024), respectively, all containing non-minicircular chromosomes (> 10 kb). Other plants with a highly multi-chromosomal mitogenome structure are the same, including 26 chromosomes of 6.0 to 32.3 kb for *Paphiopedilum micranthum* (Yang *et al*., 2023), and two species of *Silene*, with 128 chromosomes of 44.0 to 163.1 kb for *S. conica* (Sloan *et al*., 2012) and 59 chromosomes of 64.9 to 191.2 kb for *S. noctiflora* (Wu *et al*., 2015).

Considering that the available mitogenomes of all four genera in Balanophoraceae are multi-chromosomal, it is probable that the multi-chromosomal mitogenome originated in the most recent common ancestor of extant genera of Balanophoraceae, although mitogenome sequencing from other genera is needed to test this hypothesis. Because *Rhopalocnemis* is closely related to the clade comprising *Lophophytum* and *Ombrophytum* (Sanchez-Puerta *et al*., 2023), the origin of all-minicircular chromosomes in *Rhopalocnemis* must have occurred after it diverged from the clade comprising *Lophophytum* and *Ombrophytum*. The closely related congeneric species (MLP) as well as other plants with a multichromosomal mitogenome lacks a shared CR on their mitochondrial chromosomes, suggesting the shared CR in *R. phalloides* must have originated very recently, after it diverged from MLP. Similarly, mitogenome sequence heteroplasmy in *R. phalloides* must also have originated very recently after diverging from MLP.

### Mechanistic models for the CR-containing all-minicircular mitogenomic structure

The origin of all-minicircular mitochondrial chromosomes in *Rhopalocnemis* was a remarkable and unprecedented event in plants. Shao *et al*. (2009) suggested that minicircular chromosomes were likely generated from a series of events that involved the excision of fragments containing the conserved noncoding region and one or several genes near the noncoding region from a typical mitochondrial chromosome, and the rejoining of fragments over a long period. However, this model is inconsistent with our observations in *Rhopalocnemis*, because MLP has already possessed the multi-minicircular mitogenome, albeit without a conserved non-coding region.

Recently, both Lee *et al*. (2023) and Zhou *et al*. (2023) proposed that the formation of multiple smaller mitochondrial chromosomes was driven by alterations in repeat-mediated recombination due to the loss or dysfunction of mitochondrion-targeted DNA-RRR genes. However, they differ because the former hypothesized the increase of microhomology-mediated recombination while the latter proposed the decrease of homologous recombination. The latter predicts repeat degradation, which is supported by population-level mitogenome sequence comparison (Zhou *et al*., 2023). The origin of all-minicircular mitochondrial chromosomes in *Rhopalocnemis* can be also explained by the mechanism proposed by Zhou *et al*. (2023). However, we can not exclude the former hypothesis (Lee *et al*., 2023).

Surprisingly, the multi-minicircular mitogenome structure of *Rhopalocnemis* closely resembles that of some metazoan, protists and red algae, such as parasitic lice and booklice (Shao *et al*., 2009; Shao *et al*., 2012; Wei *et al*., 2012; Song *et al*., 2019), the nematode *Globodera pallida* (Armstrong *et al*., 2000; Gibson *et al*., 2007), the mesozoan (Watanabe *et al*., 1999), myxozoans (Yahalomi *et al*., 2017) and Stylonematophyceae red algae (Lee *et al*., 2023), as well as plastid genomes of free-living dinoflagellates (Zhang *et al*., 1999; Barbrook & Howe, 2000), implying convergent evolution of organellar genome structure across diverse eukaryotes (Yu *et al*., 2022). Interestingly, most of these organisms with a multi-minicircular mitogenome structure are parasites, implying that relaxed selection associated with a parasitic lifestyle may facilitate the fragmentation of ancestral mitogenomes with a single circular structure. The advantage of rapid replication for much shorter chromosomes could then be the major selective factor maintaining the multi-minicircular mitogenome structure (Shao *et al*., 2009).

How did the CR originate? One model suggests that minicircular plastid chromosomes and the shared non-coding region originate from the repeated transposition of the replication origin sequences and subsequent intra- and inter-chromosomal recombination (Zhang *et al*., 2002). Another model was proposed by Shao *et al*. (2009), as mentioned above. However, these models predict the simultaneous origin of minicircular chromosomes and the CR, inconsistent with the fact that the origin of the CR is later than the origin of all-minicircular mitogenome in *Rhopalocnemis*. Moreover, the CR sequences in *R. phalloides* have no homology with any available plant sequences, making rapid turnover of native mitochondrial non-coding sequences in a relatively short evolutionary time less likely.

Here we propose a new model to explain the origin of the CR on the minicircular chromosomes of *R. phalloides*. After diverging from MLP, *R. phalloides* likely integrated an autonomous linear plasmid containing a replication origin into one of its minicircular chromosomes. Linear mitochondrial plasmids in plants are characterized by terminal inverted repeats, reduced GC content, the presence of reading frames encoding DNA and/or RNA polymerases, and the occurrence of autonomous replication (Handa, 2008; Warren *et al*., 2016). This integration could have enhanced the chromosome’s replication rate, allowing it to outcompete and replace other chromosomal copies lacking this replication origin. The replication origin and its surrounding sequences may then spread to other minicircular chromosomes via repeat-mediated recombination or transposon-mediated transposition.

There are several pieces of evidence supporting this new model. In the *R. phalloides* mitogenomes, 1) UR of Chr18 of DWS (corresponding to UR of Chr17 of WS) contains the remnants of a mitochondrial plasmid, indicated by DNA and RNA polymerase gene fragments and the lowest GC content (39.9%) relative to other chromosomes (Yu *et al*., 2022). The absence of this region in VIET may suggest subsequent loss of these DNA and RNA polymerase gene fragments. This is not surprising as these degraded gene fragments are no longer functional. 2) SC1 and SC2 of *R. phalloides* are inverted repeats. 3) The CR on each chromosome of *R. phalloides* includes a potential replication origin (Yu *et al*., 2022). These features of the *R. phalloides* mitogenome are consistent with the plasmid incorporation model.

Additionally, there is no hit when CR and SC sequences of *R. phalloides* are searched against the GenBank nr database and the MLP mitogenome using BLASTN, suggesting they arose from recent unknown foreign sequence incorporation after diverging from MLP. The larger chromosome sizes of *R. phalloides* (4.8-8.9 kb) compared to MLP (2.5-5.9 kb) further support this model, as foreign sequence integration would increase chromosomes size.

Meanwhile, the CR is extremely conserved across all mitochondrial chromosomes within *R. phalloides* individuals, likely due to concerted evolution, similar to the “core” region of dinoflagellate chloroplast genomes (Zhang *et al*., 2002) and in the “constant” region of Stylonematophyceae red algae mitogenomes (Lee *et al*., 2023). As proposed in Zhang *et al*. (2002), gene conversion may be the primary molecular mechanism for maintaining this conservation.

Additionally, MLP lacks reversed hetero-concatemers, suggesting that in individuals like DWS and WS, where these structures are present, the CR mediates homologous recombination between different chromosomes and the palindrome in the CR causes the formation of reversed concatemers, as detailed by Yu *et al*. (2022). Meanwhile, MLP lacks large palindromic sequence, suggesting that such structures are not essential for replication origin of mitochondrial DNA.

### Rapid intraspecific variability and interspecific divergence in the mitogenomes of *Rhopalocnemis*

Rapid intraspecific variability of *R. phalloides* is manifested in mitogenome structure, sequence and heteroplasmy. For mitogenome structure, DWS and WS each have one CR, while VIET has two CRs, CR1 and CR2, separated by HV3. Although the CR is well conserved among all the minicircular chromosomes in each of the three individuals of *R. phalloides*, the CRs are divergent or highly divergent among the three individuals, especially between VIET and two other individuals. Even for DWS and WS, which are closest to each other, ∼40% sequences within the CR of DWS cannot be aligned to those of WS and even in the alignable region sequence identity between them is only 92.1%. Greater sequence divergence occurs in the CRs of VIET: one of its two CRs can not be alignable with the CRs of DWS and WS. In addition, no shared motifs in HV among the three individuals, nearly no alignable sequences in SC between VIET and two other individuals, and relatively high sequence divergence in protein coding genes among the three individuals are found. Moreover, all three individuals exhibit extremely high sequence heteroplasmy throughout the mitogenomes, including mitochondrial protein-coding genes, and very few heteroplasmic variants are shared between the three individuals, also suggesting rapid intraspecific evolution in heteroplasmy.

Relaxed selection, as detected in the *R. phalloides* branches, allows the accumulation of variation, resulting in high sequence differentiation, which can explain the extremely high heteroplasmy among the three individuals of *R. phalloides*. Relaxed selection was also used to explain the extremely high substitution rate in mitochondrial genes of *Viscum scurruloideum*, another parasitic plant (Skippington *et al*., 2015).

Rapid interspecific divergence between *R. phalloides* and its congener (MLP) is also manifested in mitogenome structure, sequence and heteroplasmy, as well as chromosomal rearrangement. MLP has about twice more mitochondrial chromosomes than *R. phalloides* and quite a few chromosomal rearrangements from *R. phalloides*. Unlike *R. phalloides*, MLP has no CR, SC, HV sequences and no sequence heteroplasmy in all its chromosomes. The two species also exhibit higher sequence divergence in protein coding genes.

Rapid intraspecific and interspecific evolution of mitogenomes in *Rhopalocnemis* makes this genus an ideal system for studying evolutionary dynamics of mitogenome size, structure and heteroplasmy, and evolutionary forces driving these changes. Other interesting questions include, 1) What are the replication origins of the mitogenome of MLP, which lacks the CR on its chromosomes? 2) What is the function of the abundant “non-functional” transcripts due to the existence of heteroplasmic variants? 3) Can the “non-functional” transcripts be translated into proteins? 4) The minicircular chromosomes can ensure rapid replication, but what is the lower limit of chromosome size? Will the introns and intergenic spacers be contracted further in *Rhopalocnemis*?

### The striking heteroplasmy in the *R. phalloides* mitogenome cannot be explained by the loss of available mitochondrion targeted DNA-RRR genes

Both *Rhopalocnemis* and its closely related genus *Lophophytum* have lost two DNA-RRR genes, *PHR1* and *WHY2*, which typically repairs the majority of UV-generated DNA lesions (Dündar *et al*., 2020) and blocks microhomology-mediated end joining (García-Medel *et al*., 2019), respectively. The loss of *WHY2* or both genes may contribute to the formation of multi-chromosomal mitogenome structure, based on the hypothesis of Lee *et al*. (2023) mentioned above. Compared with *Lophophytum*, *Rhopalocnemis* has lost only one more DNA-RRR gene, *FPG1*, which encodes a DNA glycosylase and is involved in DNA repair through excision of 8-oxoG residues resulting from oxidative damage (Córdoba-Cañero *et al*., 2014; Ferrando *et al*., 2019). Given that this gene is not implicated in repeat-mediated recombination, loss of *FPG1* in *Rhopalocnemis* may not contribute to the formation of all-minicircular mitochondrial chromosomes. No *R. phalloides*-specific loss was found in the detected DNA-RRR genes. Thus, the striking difference in mitochondrial sequence heteroplasmy between *R. phalloides* and its congener could not be explained by DNA-RRR gene loss identified in this study. Loss of other undiscovered DNA-RRR genes in the common ancestor of *R. phalloides* or differential expression of available and/or undiscovered DNA-RRR genes may result in its striking heteroplasmy, a hypothesis in need of further testing.

## Materials and Methods

### Plant sampling and sequencing

Two newly collected individuals of *Rhopalocnemis*, WS and MLP, were sampled from Bozhu, Wenshan and Malipo, Wenshan, Yunnan, China, respectively. The inflorescence tissues were used for DNA extraction, RNA extraction and sequencing. The procedures of DNA and RNA extraction, library construction and Illumina sequencing followed Yu *et al*. (2022). For PacBio sequencing, a DNA library with an insert size of 15 kb was constructed for each of the two individuals and sequenced on a PacBio Sequel II platform using the CCS (HiFi) mode. Plant sampling and sequencing of two other individuals of *R. phalloides*, DWS and VIET, were described in Yu *et al*. (2022) and Schelkunov *et al*. (2019), respectively. Sampling and sequencing data details were shown in Table S1.

### Mitogenome assembly

Mitogenomes of WS, VIET and MLP were assembled using their Illumina DNA-seq reads in GetOrganelle v1.7 (Jin *et al*., 2020) with the parameters “-F embplant_mt -k 127”. Mitochondrial contigs for each individual were extracted in Bandage v0.8.1 (Wick *et al*., 2015) following the method of Yu *et al*. (2022). Mitogenome structures of WS and VIET (Figure S8) highly resemble that of DWS (Yu *et al*., 2022), and 20 and 18 circular chromosomes could be resolved, respectively. The mitogenome structure of MLP is relatively simple (no CR, HV and SC, see Results) and was assembled into 39 circular chromosomes.

To extract mitochondrial PacBio sequencing reads for WS, we mapped all PacBio sequencing reads to its assembled mitogenome using NGMLR v0.2.7 (Sedlazeck *et al*., 2018) with parameters “-x pacbio -i 0.90 -R 0.7”. We used the consensus sequences of mitochondrial PacBio reads for each chromosome to resolve the issue of extremely high sequence heteroplasmy. For VIET, which has no available PacBio sequencing data, we mapped Illumina reads to its assembled mitogenome and used the consensus sequences of the unique regions (URs), conserved regions (CRs) and semi-conserved regions (SCs), respectively, for each chromosome. We randomly selected one of the many alternative paths for each of the three hypervariable regions (HVs) from the assembled mitogenome graph as the sequences of HVs for all its chromosomes. The orders of all parts on the chromosomes of WS and VIET used for presentation and submission to GenBank are CR-HV1-SC1-UR-SC2-HV2 and HV2-CR2-HV3-CR1-HV1-SC1-UR-SC2, respectively.

### Mitogenome annotation

To annotate protein-coding and rRNA genes in the mitogenomes of WS, VIET and MLP, we searched the assembled mitochondrial chromosomes of WS, VIET and MLP with a custom angiosperm mitochondrial gene dataset as a query using BLASTN 2.2.28 (Altschul *et al*., 1990) with the e-value set to 1e-3. Due to the high sequence heteroplasmy for almost all genes of WS and VIET, the “functional” version of each gene was used for the two individuals. The “functional” versions of the genes were obtained by mapping Illumina sequencing reads of the two individuals to their assembled mitochondrial chromosomes and then comparing the mitochondrial genes of five other angiosperms, namely, *Liriodendron tulipifera*, *Arabidopsis thaliana*, *Vitis vinifera*, *Malania oleifera* and *Ombrophytum subterraneum* (Table S16). When necessary, manual adjustments were made for start and stop codon positions to ensure the accuracy of gene annotation. At each heteroplasmic position, variant sequences identical to most other species were used. tRNA genes were annotated using tRNAscan-SE v2.0 (Lowe & Chan, 2016) with the “organelle” mode. Gene map of the mitochondrial genomes were drawn using GenomeVx (http://wolfe.ucd.ie/GenomeVx/).

### Plastome assembly and annotation of WS and MLP

Plastomes of WS and MLP were assembled with their Illumina DNA-seq reads using GetOrganelle v1.7 (Jin *et al*., 2020) and setting the parameters “-F embplant_pt -k 127”. The protein-coding genes of WS and MLP plastomes were annotated via PGA (Qu *et al*., 2019) under default parameters and subsequently manually verified.

### Calculation of pairwise sequence identity

To compare the sequence similarity of mitochondrial protein-coding genes among four individuals of *Rhopalocnemis*, we calculated pairwise sequence identity for each of the 36 genes using the “*align of two or more sequences*” function of BLASTN. We also calculated the average identity of the 36 genes and plotted a boxplot to show pairwise identity distribution. The same method was used to calculate pairwise sequence identity of 13 plastid protein-coding genes for four *Rhopalocnemis* individuals (plastid genes of DWS and VIET are from Yu *et al*. (2022) and Schelkunov *et al*. (2019), respectively).

### Phylogeny reconstruction

We downloaded sequences of nuclear 18S and 28S rRNA genes and three available mitochondrial genes (*cox3*, *matR* and *rrn18*) of eight species of Balanophoraceae and one species of Olacaceae (*Malania oleifera*) from GenBank (Table S17). Sequence alignment for the above nine species and four individuals of *Rhopalocnemis* was performed for the two concatenated nuclear genes and three concatenated mitochondrial genes separately using MUSCLE v3.8.31 (Edgar, 2004). The alignments were then modified manually and were trimmed using Gblocks 0.91b (Castresana, 2000). The maximum likelihood (ML) trees were constructed for the nuclear and mitochondrial sequence matrix separately using RAxML (Stamatakis, 2014) with the GTR + Gamma model and 1000 bootstrap replications. The trees were then visualized in the iTOL website (https://itol.embl.de/).

### Identification of horizontal gene transfer

To infer potential horizontal gene transfer in the *Rhopalocnemis* mitogenomes, we extracted mitochondrial protein-coding genes from 121 diverse angiosperms, including the four individuals of *Rhopalocnemis*, the five previously sequenced Balanophoraceae species, six species from other families of Santalales, and 106 other angiosperms (Table S16). Foreign genes in the mitogenome of *Ombrophytum* (Roulet *et al*., 2020) were excluded. The alignment, trimming of poor-alignment regions, and phylogenetic analysis for each of the 36 genes followed the method in the “Phylogeny reconstruction” section above. We used *Amborella* as the outgroup for all mitochondrial genes except for *rpl10*, for which we used *Liriodendron* as the outgroup.

### Calculation of nucleotide substitution rates and testing of relaxed selection

To calculate nucleotide substitution rates, we concatenated 30 shared mitochondrial protein-coding genes (*atp1*, *atp4*, *atp6*, *atp8*, *atp9*, *ccmB*, *ccmC*, *ccmFC*, *ccmFN*, *cob*, *cox1*, *cox2*, *cox3*, *matR*, *mttB*, *nad1*, *nad2*, *nad3*, *nad4*, *nad4L*, *nad5*, *nad6*, *nad7*, *nad9*, *rpl5*, *rps10*, *rps12*, *rps3*, *rps4*, *sdh4*) for the four *Rhopalocnemis* individuals and 10 other angiosperms (Table S16). Sequence alignment and trimming of unreliable alignment regions for the 30 genes followed the method in the “Phylogeny reconstruction” section above. The synonymous and non-synonymous nucleotide substitution rates (*d_N_*and *d_S_*) were calculated using codeml in PAML (Yang, 2007), with the free ratio mode and F3×4 codon frequency. The topologically constrained tree is based on Nickrent (2020) and Figure 1 of this study.

To test whether the 30 concatenated mitochondrial genes showed relaxed selection in the *R. phalloides* branches than other branches, we calculated ω values under the branch model using codeml in PAML. We performed likelihood ratio tests (LRT) to see if there is significant difference between two models: (1) one *d_N_*/*d_S_* ratio across all branches as the null hypothesis (model = 0; NSsites = 0), (2) one ratio for the *R. phalloides* clade (all branches of *R. phalloides*) as foreground branch and the other ratio for all other branches as background branch, as the alternative hypothesis (model = 2; NSsites = 0). The likelihood values of the two models were used to test statistical significance with a Chi-square test. If the ω value of the foreground branch is significantly larger than that of the background branch, and both are less than 1, then it will be considered as evidence for relaxed selection on the foreground branch. The same test was also conducted to test relaxed selection in the *Rhopalocnemis* clade (all branches of *Rhopalocnemis*).

### Mitogenome collinearity analysis

Homologous regions of > 50 bp in length among the chromosomes of MLP and URs of DWS, WS and VIET were identified using BLASTN with the parameters “e-value and perc_identity” set to 1e-5 and 85, respectively, and plotted using MCscan in JCVI v1.2.7 (Tang *et al*., 2008). Homologous sequences between the CR and SC regions in the mitogenomes of DWS, WS and VIET were also identified using BLASTN with the same parameters. We made a dot plot to show the homology of the SC regions using Flexidot v1.06 (Seibt *et al*., 2018) with the parameters “-k 15 -S 1”.

### Identification of heteroplasmic variants

Mitogenome heteroplasmy for WS and VIET was analyzed using the same method of Yu *et al*. (2022). Here we focused on protein-coding genes only for comparison among different individuals, because their sequences are more conserved. Briefly, Illumina DNA sequencing reads of WS and VIET were mapped to their own assembled mitogenomes using BWA-mem (Li, 2013), respectively. Single nucleotide variants (SNVs), insertions/deletions and complex variants were then identified using VarDict (Lai *et al*., 2016) with the parameters “-X 5 -f 0.05”.

To examine whether these heteroplasmic variants transcribe or not, Illumina RNA-seq reads of WS and VIET were mapped to their own mitogenomes using HISAT2 v2.1.0 (Kim *et al*., 2015) with the parameters “--sensitive --no-mixed --no-discordant”. Heteroplasmic variants were also identified using VarDict with the parameters “-X 5 -f 0.05 -m 20”. Correlation analyses of variant frequencies at the DNA level and at the RNA level were performed using ggplot2 (Villanueva & Chen, 2019).

### Extraction of mitochondrial PacBio HiFi reads

We extracted mitochondrial PacBio reads of WS and MLP by mapping all their PacBio reads to their own assembled mitogenomes, using NGMLR v0.2.7 (Sedlazeck *et al*., 2018) with the parameters “-x pacbio -i 0.90 -R 0.7”. Then, we counted the number of chromosomes contained in the mitochondrial PacBio reads and divided the reads into homo-concatemers (composed of the same chromosome unit) and hetero-concatemers (composed of at least two different chromosome units).

### Repeat identification and characterization of repeat-mediated recombination in the formation of hetero-concatemers

Repeat pairs in the entire mitochondrial chromosomes of MLP and the URs of DWS, WS and VIET were identified using a perl script ROUSFinder2.0.py (Wynn & Christensen, 2019) with the minimum repeat size set to 30 bp. The CRs, SCs and HVs of the latter three are excluded from this analysis because they are completely or largely shared among different chromosomes.

To verify if a hetero-concatemer in the mitogenome of WS and MLP results from repeat-mediated recombination, we examined the sequences flanking the putative recombination breakpoints. If the immediate flanking sequences contain a repeat we identified above, this hetero-concatemer is considered to result from repeat-mediated recombination.

### Characterization of mitochondrion-targeted DNA-RRR genes

For each of the four individuals of *Rhopalocnemis*, Illumina RNA-seq reads were used to assemble the transcriptome using Trinity v2.11.0 (Grabherr *et al*., 2011) with default parameters except for --min_kmer_cov set to 2. The completeness of the assembled transcriptomes was evaluated using Benchmarking Universal Single-Copy Ortholog (BUSCO) v5.1.3 (Simao *et al*., 2015) with the transcriptome mode based on the eukaryota_odb10 datasets.

24 available mitochondrion-targeted DNA-RRR genes used in Ceriotti *et al*. (2022) were searched in the transcriptomes and PacBio/Illumina DNA reads of the four individuals of *Rhopalocnemis* to determine whether they are present or absent. For the assembled transcriptomes, we predicted open reading frame of transcripts, removed redundant transcripts, and clustered pre-computed orthogroups based on the built-in 22G v1.1 dataset using AssemblyPostProcessor module with targeted gene family model and GeneFamilyClassifier module of PlantTribes2 (Wafula *et al*., 2022). If a transcript or multiple transcripts of an individual of *Rhopalocnemis* was classified into the same orthogroup as an *Arabidopsis* DNA-RRR gene, we considered the presence of this gene in this individual. In two cases where multiple DNA-RRR genes are in the same orthogroup (*OSB1*, *OSB2-4* and *OSBX*; *RECA2* and *RECA3*), we aligned the sequences of multiple genes using MAFFT in GeneFamilyAligner module and constructed the maximum likelihood tree using RAxML in GeneFamilyPhylogenyBuilder module of PlantTribes2 to determine the orthologous transcripts for each gene (Figure S9).

Moreover, we also searched the 24 DNA-RRR genes of *Arabidopsis* against the PacBio long reads of DWS, WS and MLP, using TBLASTN with e-value set to 1e-3, to infer their status in the genomes. Because VIET has no available PacBio long reads, we assembled and used its nuclear draft genome with Illumina reads using SPAdes v4.0.0 (Bankevich *et al*., 2012). Only those genes that are undetected in both the transcriptome and the genome were considered to be truly lost.

## Acknowledgments

This study was financially supported by the National Natural Science Foundation of China (31811530297 and 32170217).

## Author Contributions

R.Z. and Y.L. conceived the study and supervised the project. Samples were collected by R.Z., and R.Y. Y.Z., R.Y., C.S., Y.L., J.P.M. and M.V.S. analyzed the data. R.Z. and Y.Z. drafted the manuscript. R.Z., Y.Z., J.P.M. and M.V.S. revised the manuscript. All authors reviewed and approved the final manuscript.

## Data Availability

The mitogenome assemblies and annotations for *Rhopalocnemis* have been deposited in National Center for Biotechnology Information (NCBI) under the accession numbers: PQ773375-PQ773394 for WS, PQ773357-PQ773374 for VIET, and PQ801611-PQ801649 for MLP. The plastome assemblies and annotations for WS and MLP are available in NCBI with accession numbers PQ849602 and PQ849603. The raw Illumina DNA-Seq and RNA-Seq reads, and the raw PacBio reads of WS and MLP are accessible under NCBI BioProject accession number PRJNA1195246. See Table S1 for details.

---

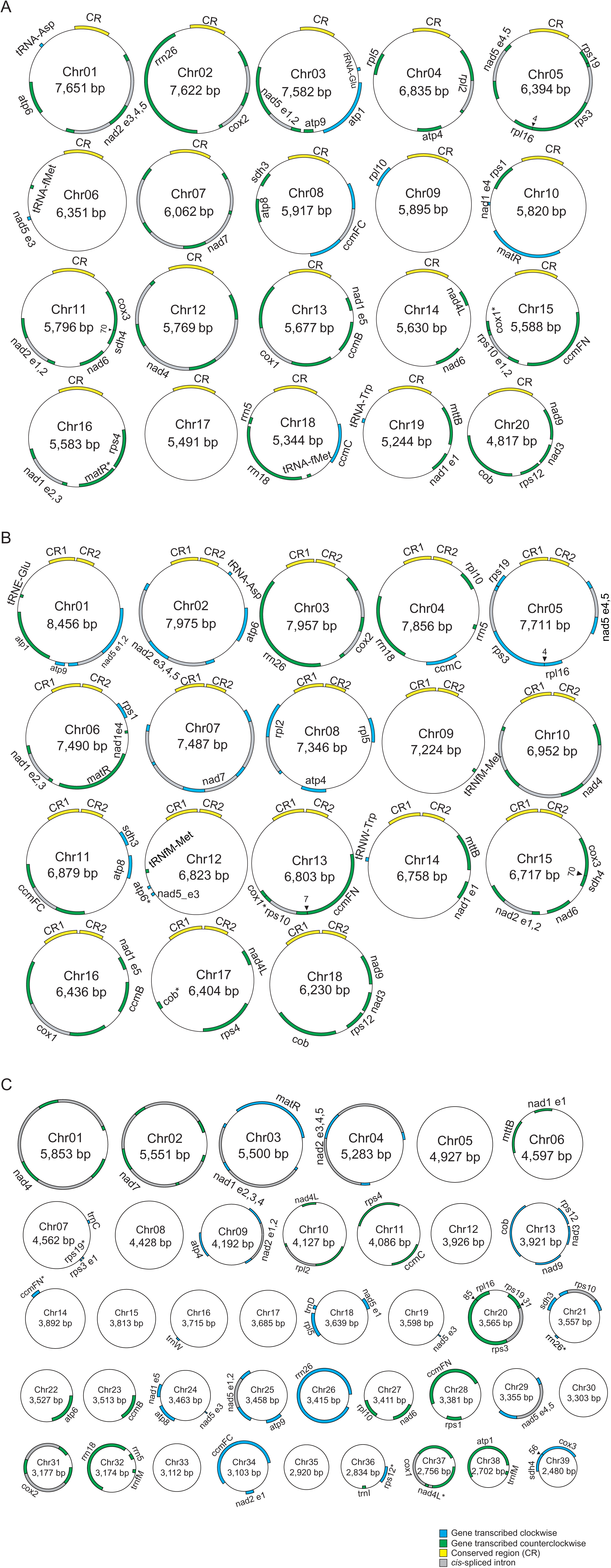

