## Supplemental information for "Origin and rapid evolution of minicircular and highly heteroplasmic mitogenome in the holoparasitic plant genus *Rhopalocnemis*"

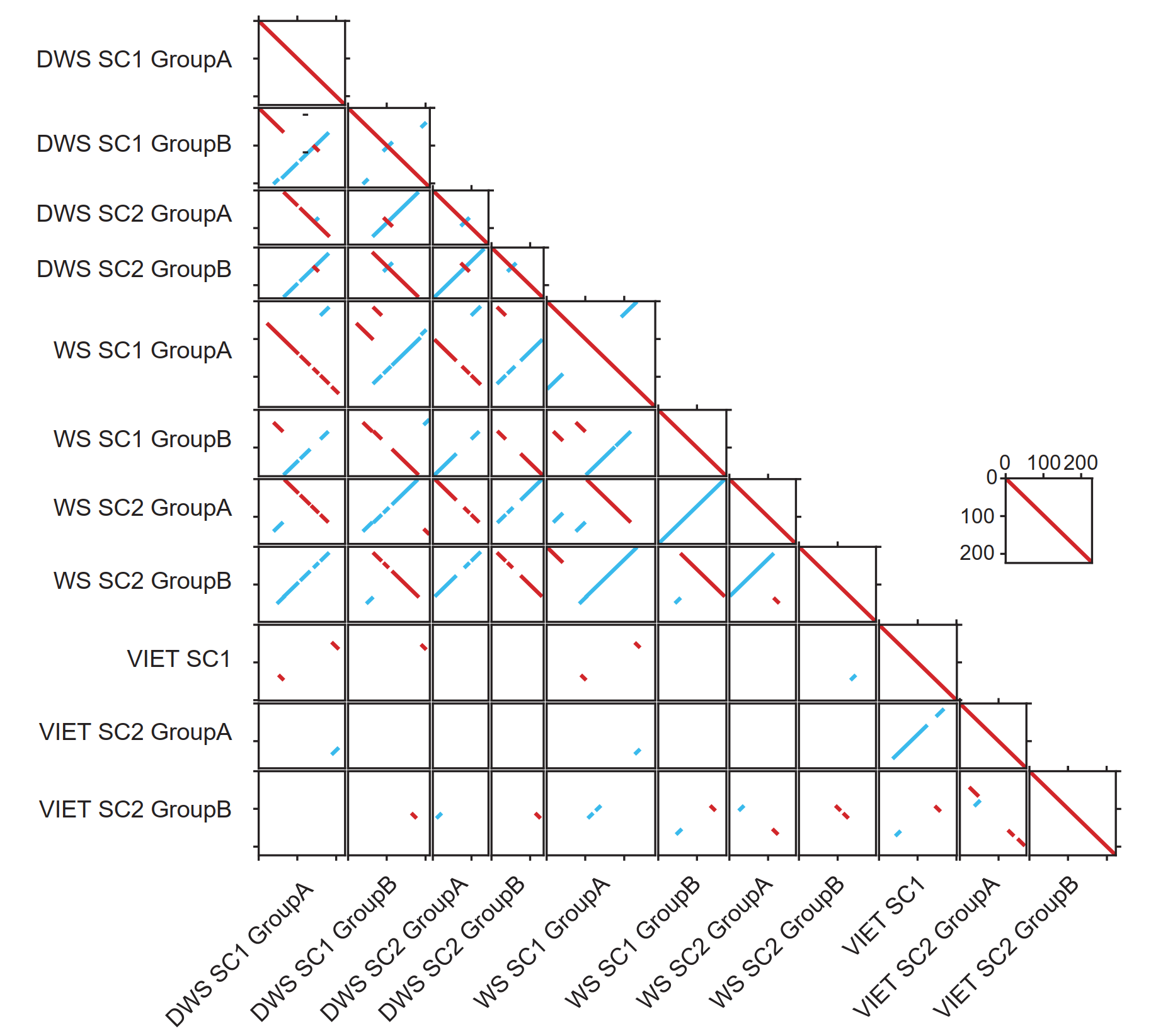

**Figure S2.** Dotplot analysis for mitochondrial semi-conserved regions (SC1 and SC2) among three individuals of *Rhopalocnemis phalloides* (DWS, WS and VIET). Identical or highly similar regions in the same orientation and in reverse compliment are marked in red and blue, respectively.

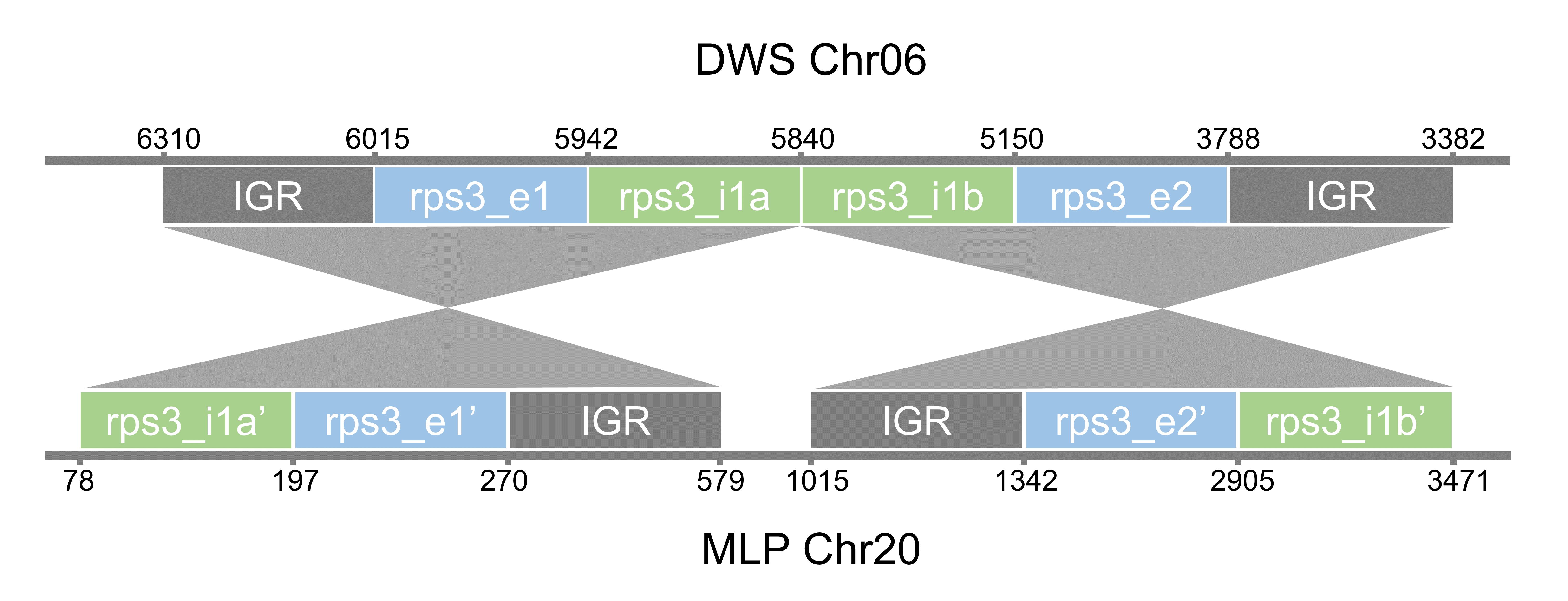

**Figure S3.** **Shift from *cis*-splicing to *trans*-splicing of the mitochondrial *rps3* gene shown by synteny analysis of this gene between *Rhopalocnemis phalloides* (DWS, *cis*-splicing) and its congener (MLP, *trans*-splicing).**

Two exons (marked by e1 and e2) and one intron (marked by two parts: i1a and i1b) of this gene are shown as blue and green rectangles with genomic coordinates on their own chromosomes, respectively. Intergenic regions (IGR) are shown as gray rectangles. Gray shading indicates homologous regions between DWS and MLP.

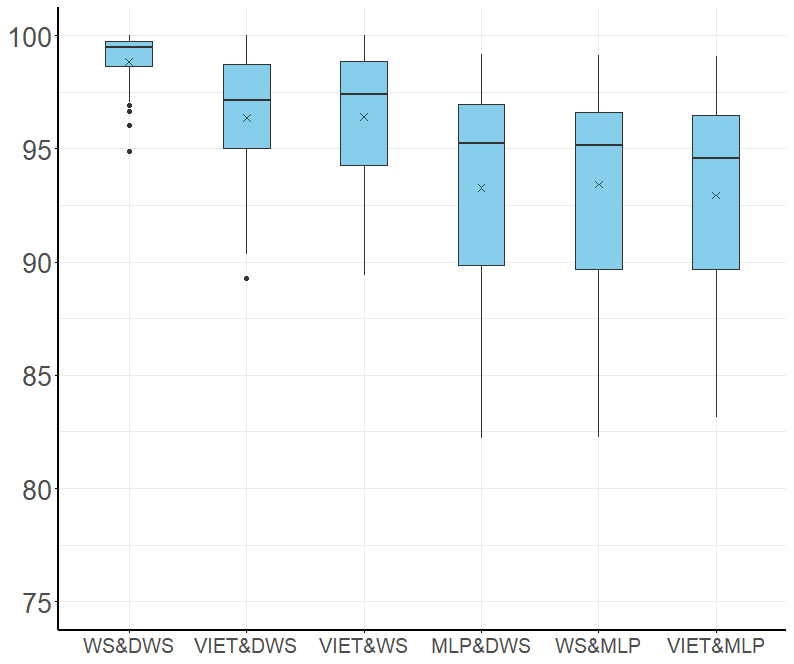

**Figure S5.** Pairwise sequence identity of 36 mitochondrial protein-coding genes among four individuals of *Rhopalocnemis* (WS, DWS, VIET and MLP). The vertical ordinate shows sequence identity (%). The boxes correspond to the first and third quartiles of sequence identity of 36 genes. The horizontal line and cross symbol inside the boxes correspond to median and mean of sequence identity of 36 genes. The whiskers extend to the smallest and largest values within 1.5-fold the interquartile range, respectively. Data that fall outside of the range of the whiskers are marked with dots.

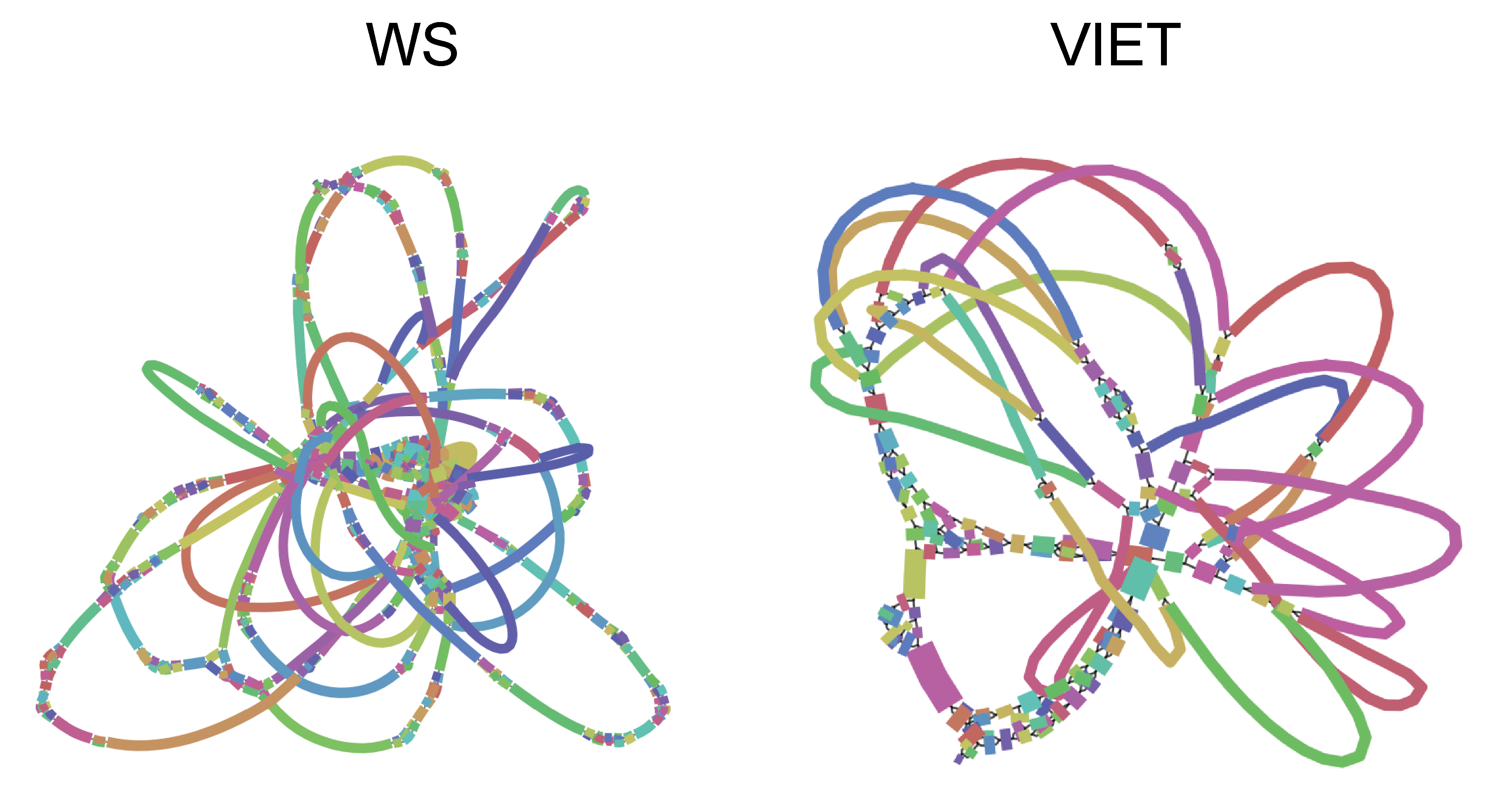

**Figure S7.** The mitogenome assembly graph of two individuals of *Rhopalocnemis phalloides* (WS and VIET) visualized in Bandage based on the raw GFA file produced by GetOrganelle. The contigs colors were displayed randomly.
