## Supplementary material for "Origin and rapid evolution of minicircular and highly heteroplasmic mitogenome in the holoparasitic plant genus *Rhopalocnemis*": FigureS1-S9

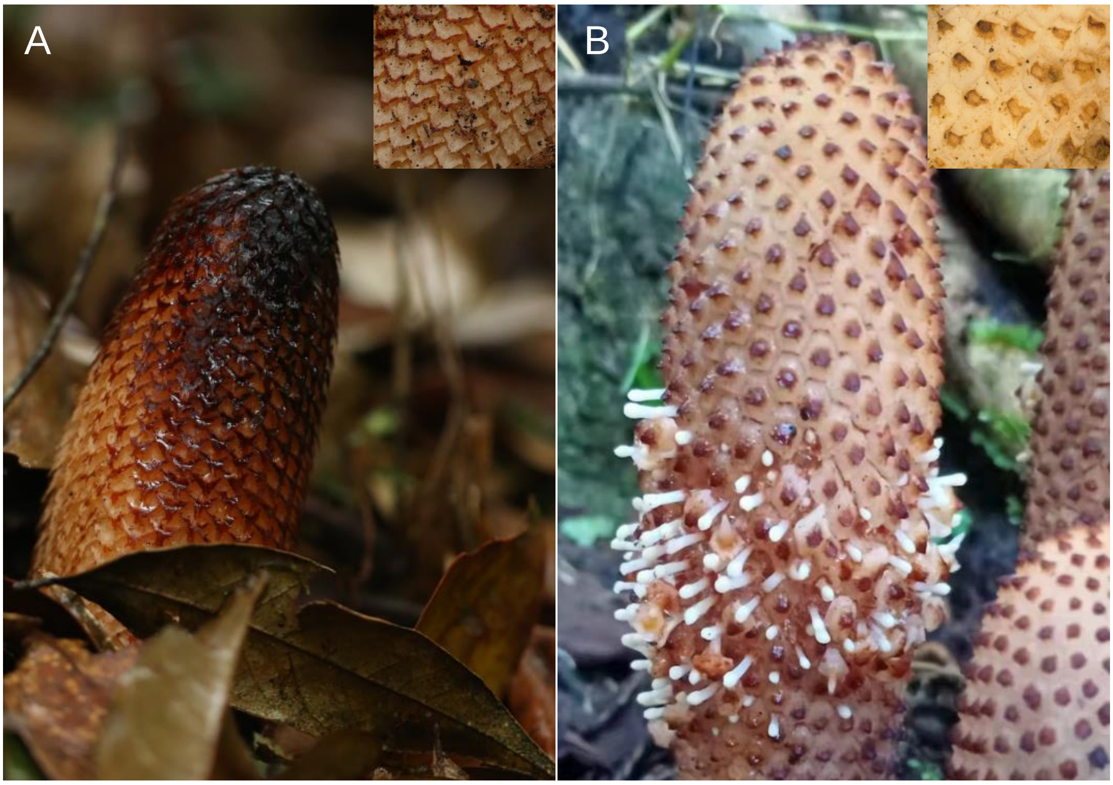


**Figure S1**. Photos of inflorescences of *Rhopalocnemis phalloides* (A) and its congener (B). The insets are magnified scales on their inflorescence surfaces.

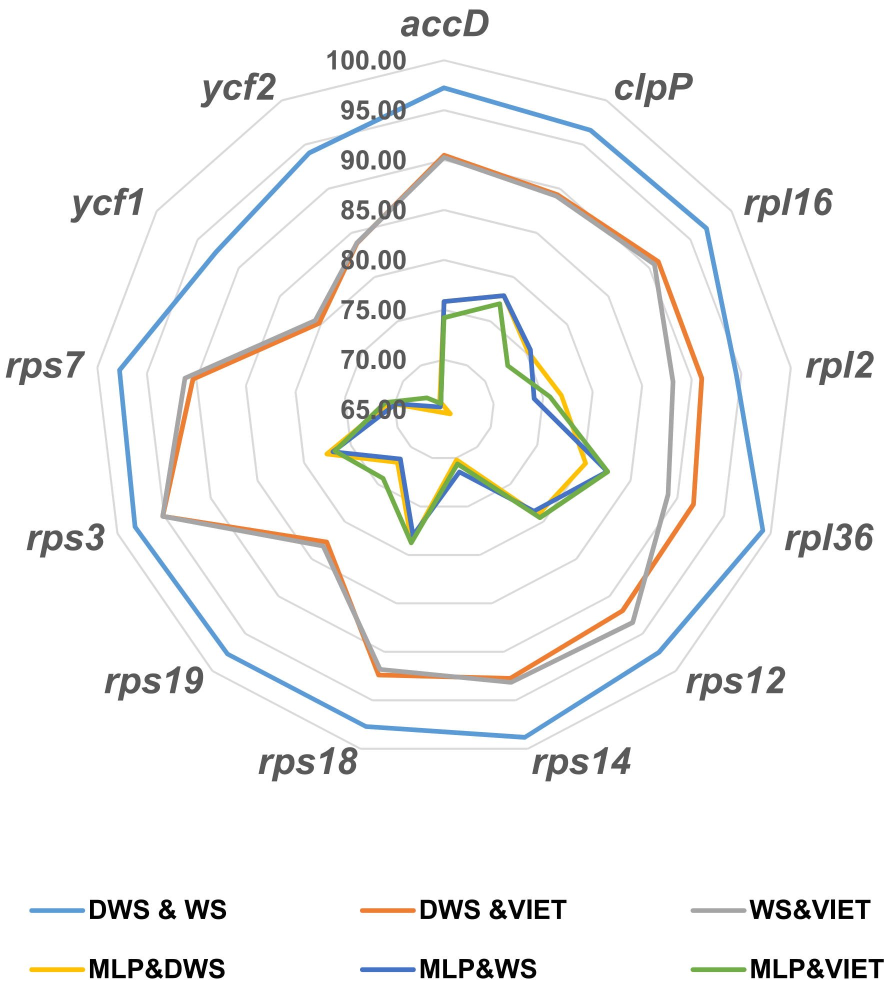


**Figure S2.** The radar map of pairwise identity between plastid genes of four *Rhopalocnemis* individuals.

WS

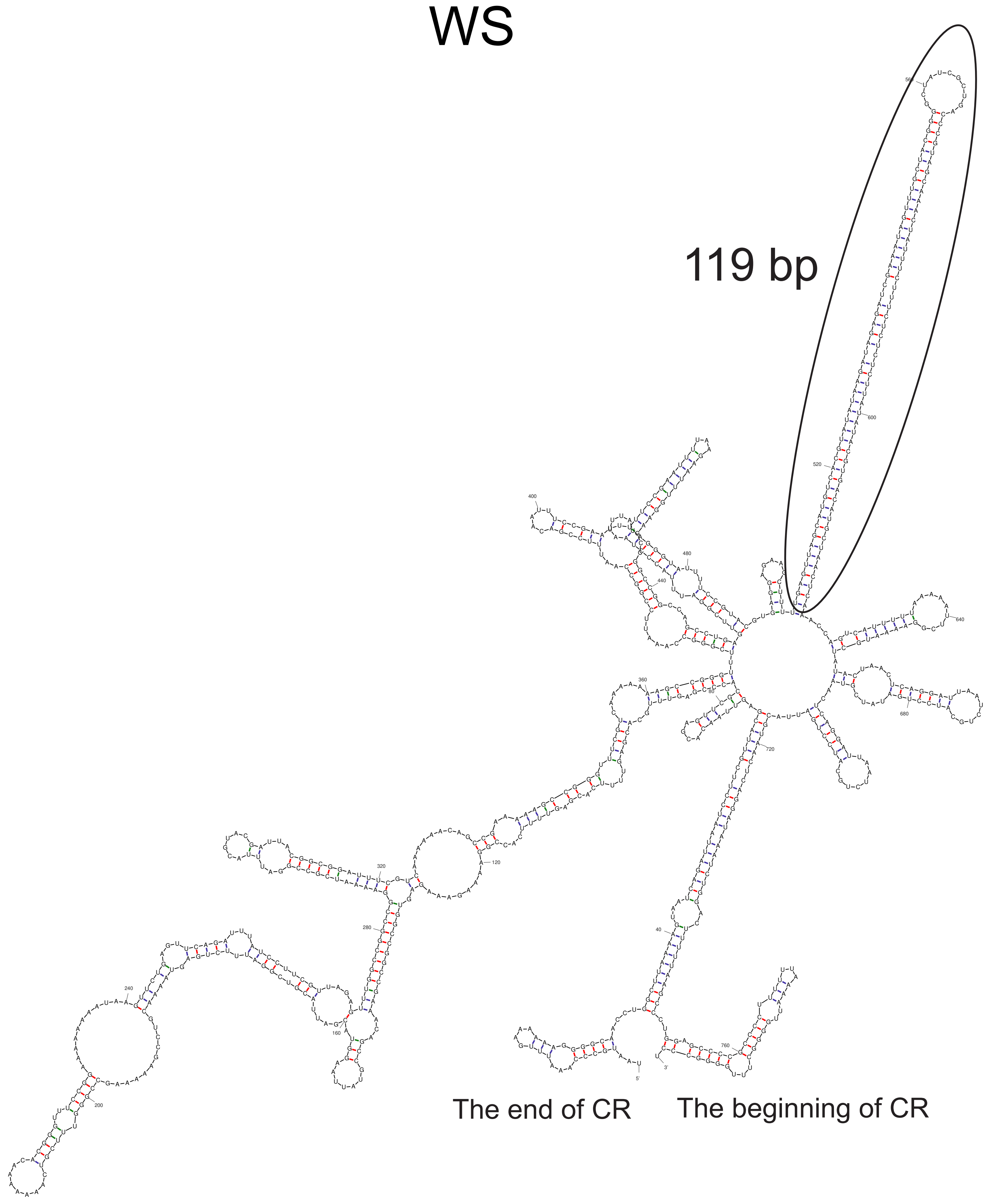


VIET

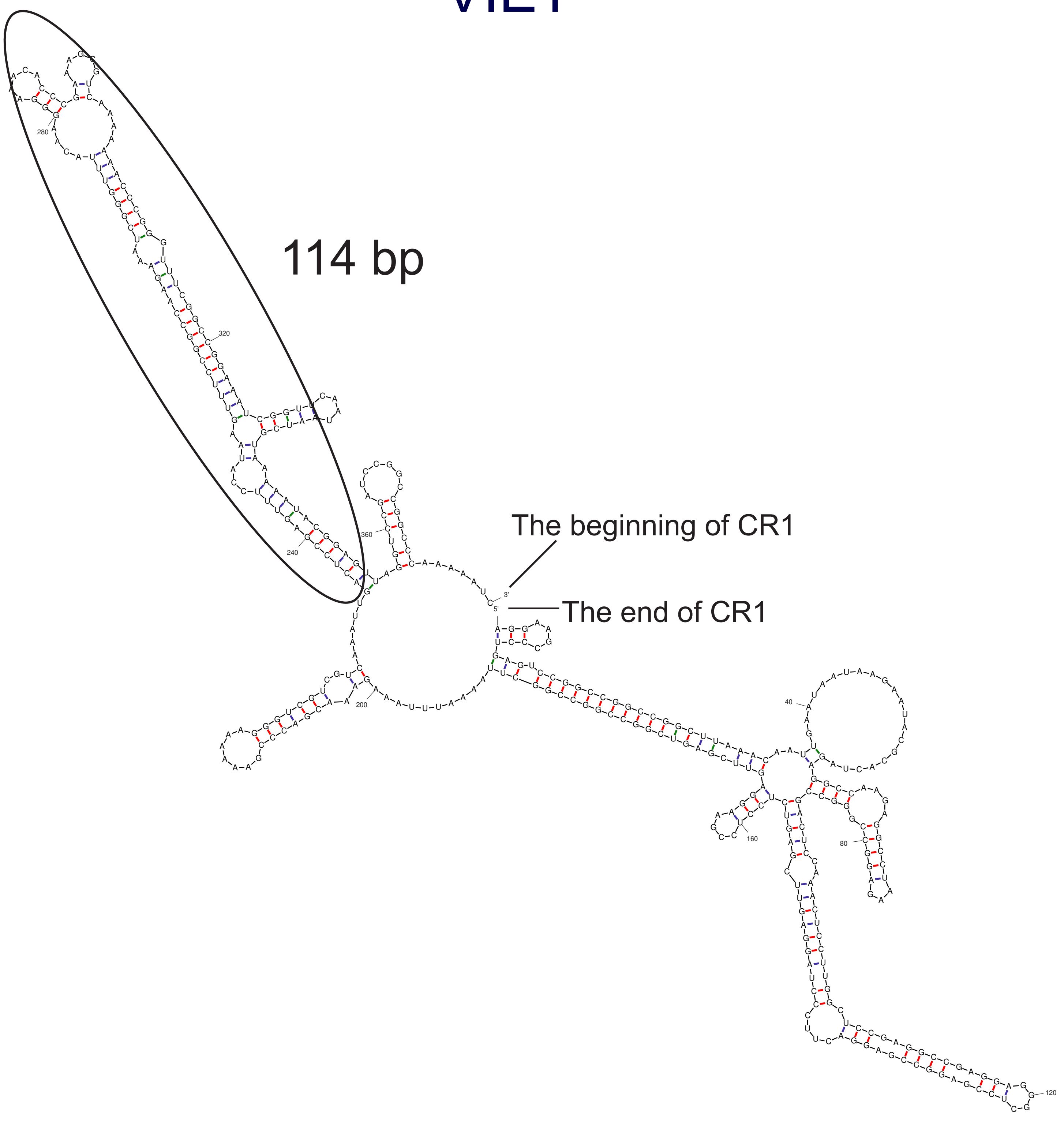


**Figure S3**. Single-stranded mRNA secondary structure of the CR in the mitogenomes of two individuals (WS and VIET) of *Rhopalocnemis phalloides*. For VIET, only CR1 was used here because its CR2 does not contain a palindrome. Regions enclosed with black lines denote the large palindromes.

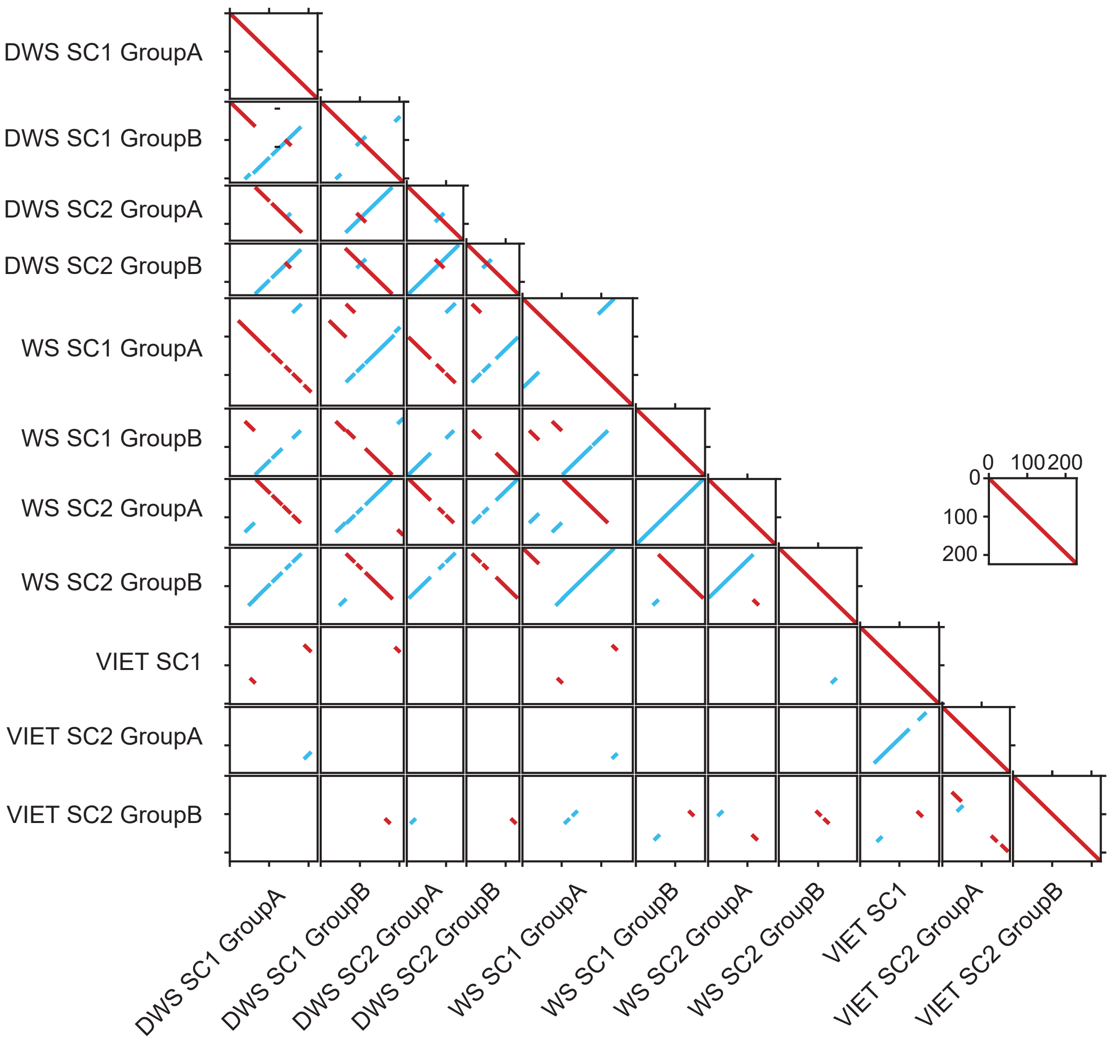


**Figure S4**. Dotplot analysis for mitochondrial semi-conserved regions(SC1 and SC2) among three individuals of *Rhopalocnemis phalloides*(DWS, WS and VIET). Identical or highly similar regions in the same orientation and in reverse compliment are marked in red and blue, respectively.

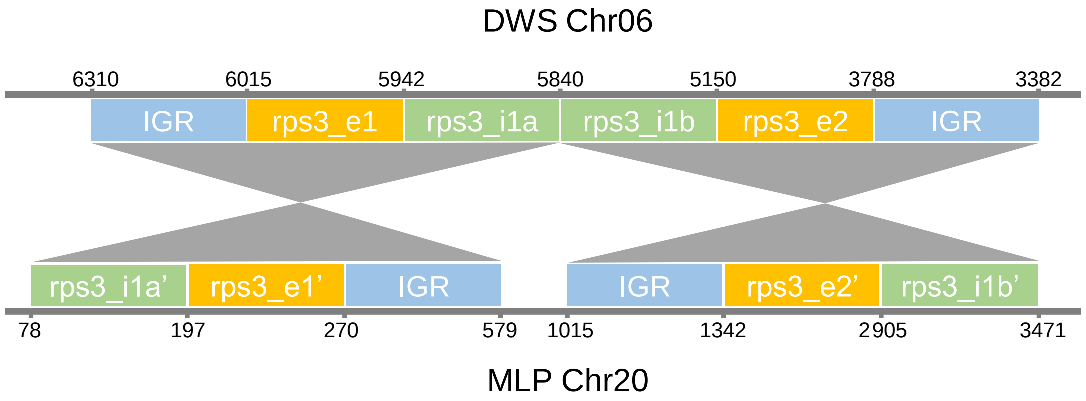


**Figure S5**. Shift from *cis*-splicing to *trans*-splicing of the mitochondrial *rps3* intron in MLP (*Rhopalocnemis* sp.) based on synteny analysis with DWS (*Rhopalocnemis phalloides*). *Cis*-splicing of this intron is observed in DWS. Two exons (e1 and e2) and one intron (divided into two parts: i1a and i1b) of this gene are shown as yellow and green rectangles, respectively, with nucleotide positions shown on their own chromosomes. Blue rectangles represent Intergenic regions (IGR). Gray shading indicateshomologous regions between DWS and MLP.

#### ***atp1***

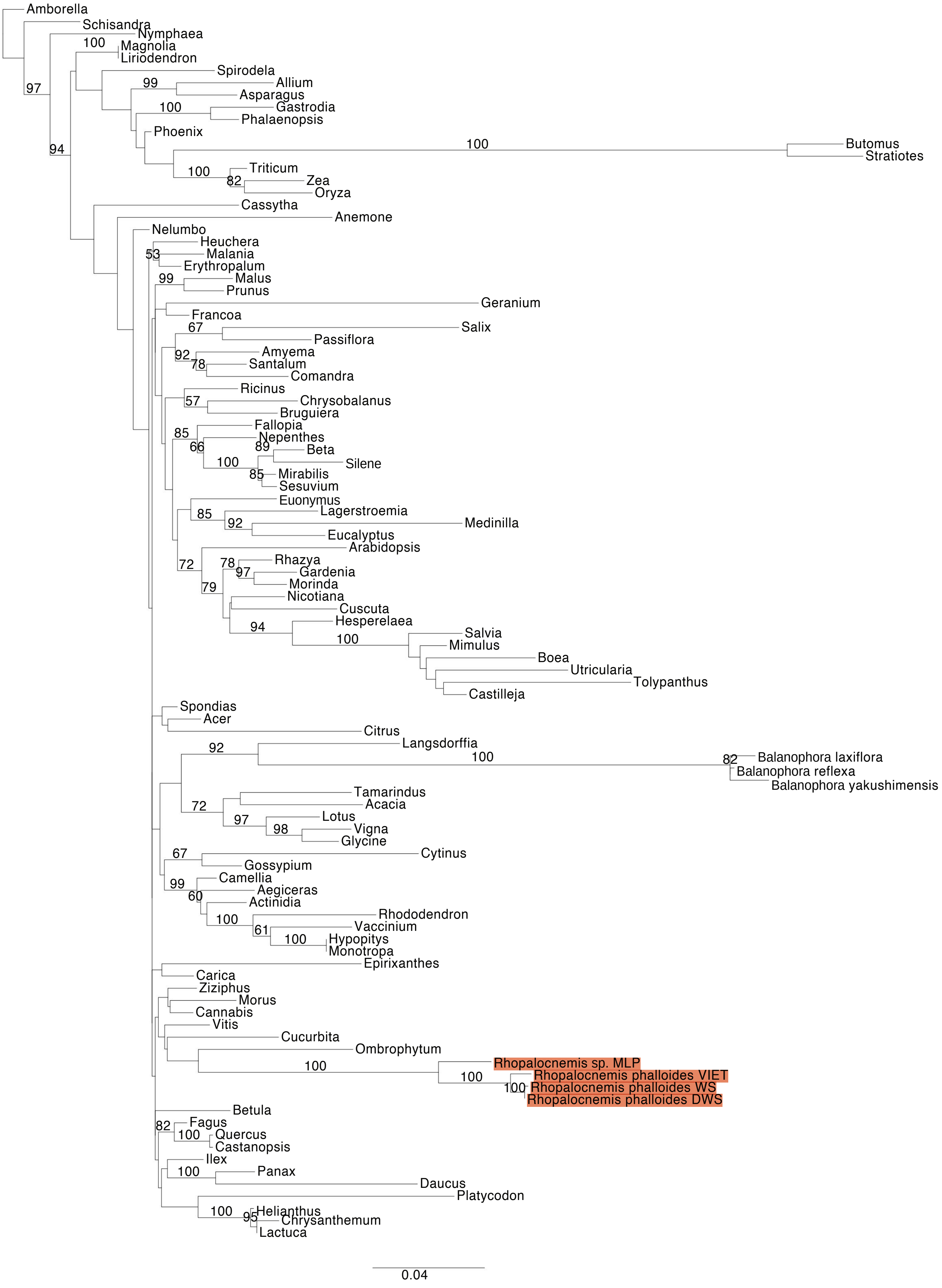


**Figure S6.** Maximum Likelihood (ML) phylogenetic analyses of protein-coding gene sequences in the mtDNAs of *Rhopalocnemis*. ML bootstrap support values ≥50% are shown. Scale bars correspond to substitutions per site.

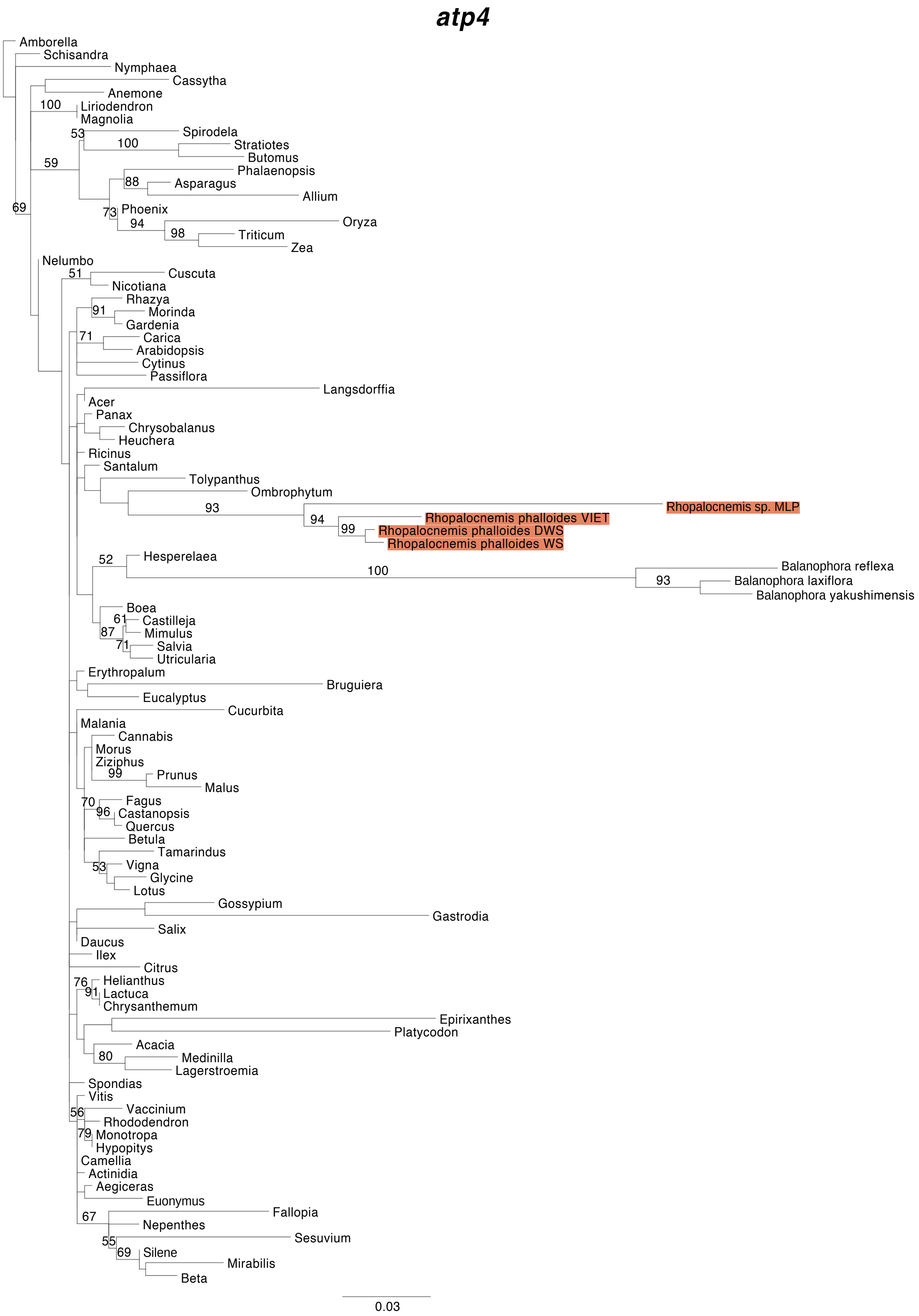


**Figure S6.** Maximum Likelihood (ML) phylogenetic analyses of protein-coding gene sequences in the mtDNAs of *Rhopalocnemis*. ML bootstrap support values ≥50% are shown. Scale bars correspond to substitutions per site.

#### *atp6*

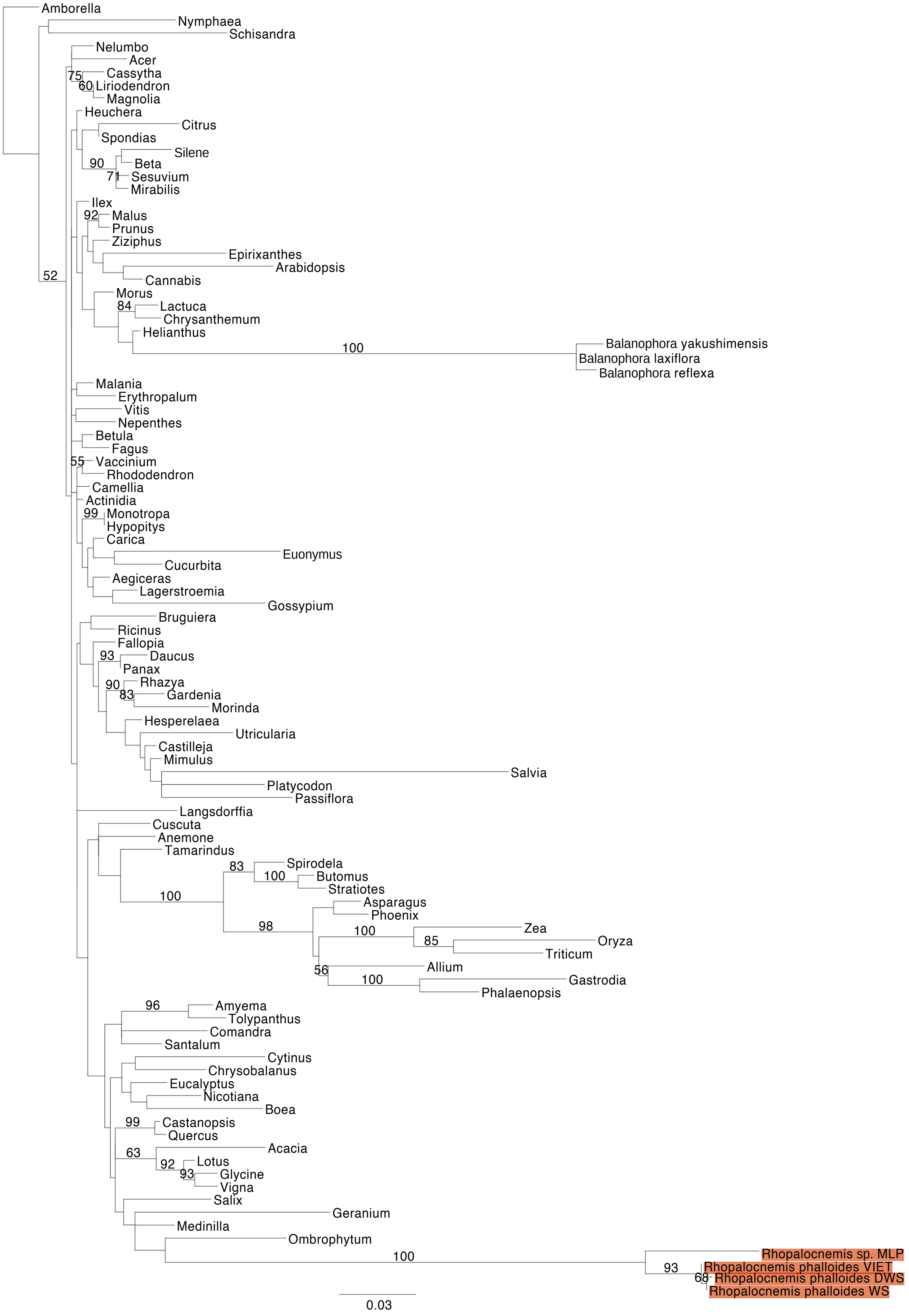


**Figure S6.** Maximum Likelihood (ML) phylogenetic analyses of protein-coding gene sequences in the mtDNAs of *Rhopalocnemis*. ML bootstrap support values ≥50% are shown. Scale bars correspond to substitutions per site.

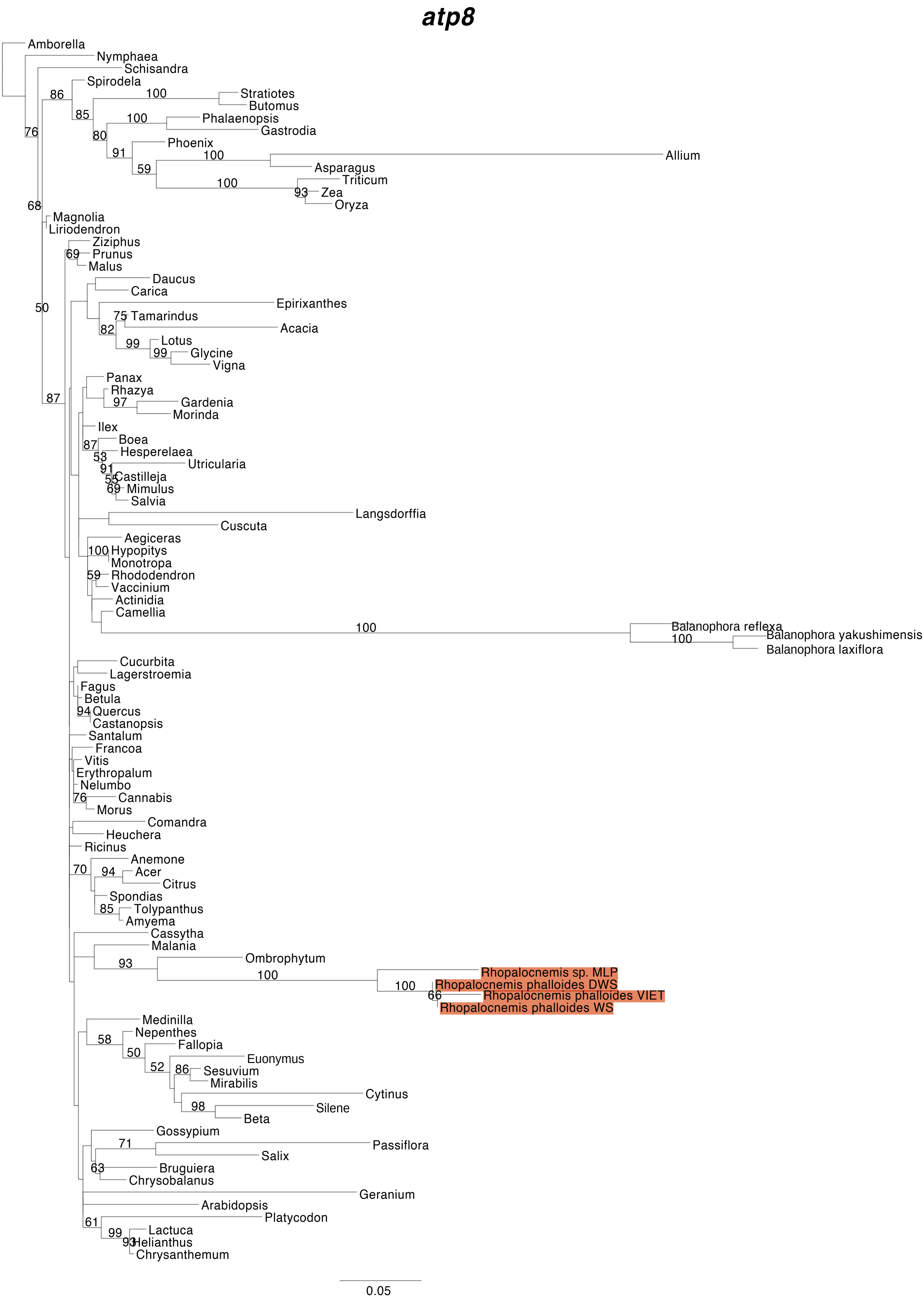


**Figure S6.** Maximum Likelihood (ML) phylogenetic analyses of protein-coding gene sequences in the mtDNAs of *Rhopalocnemis*. ML bootstrap support values ≥50% are shown. Scale bars correspond to substitutions per site.

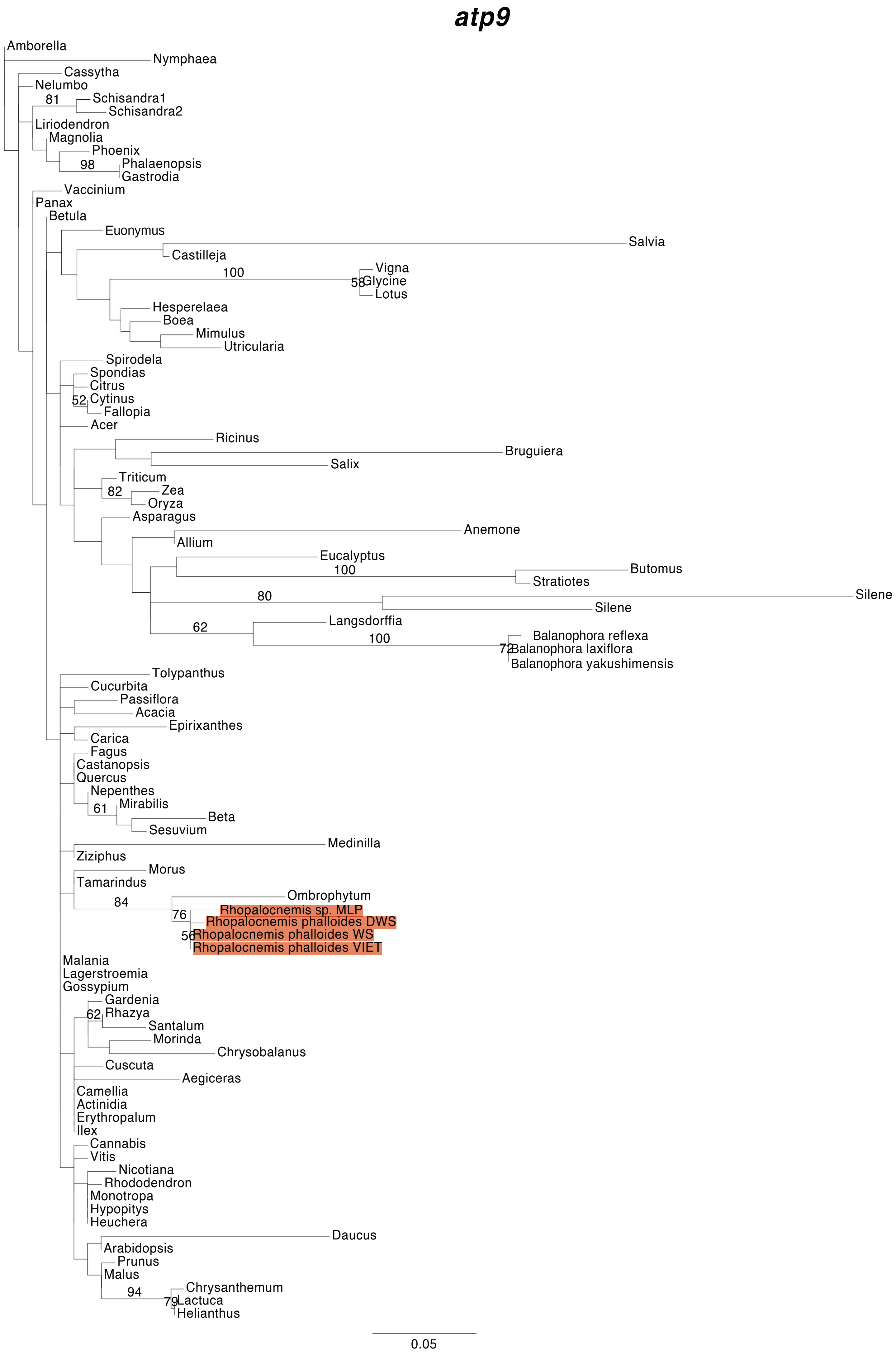


**Figure S6.** Maximum Likelihood (ML) phylogenetic analyses of protein-coding gene sequences in the mtDNAs of *Rhopalocnemis*. ML bootstrap support values ≥50% are shown. Scale bars correspond to substitutions per site.

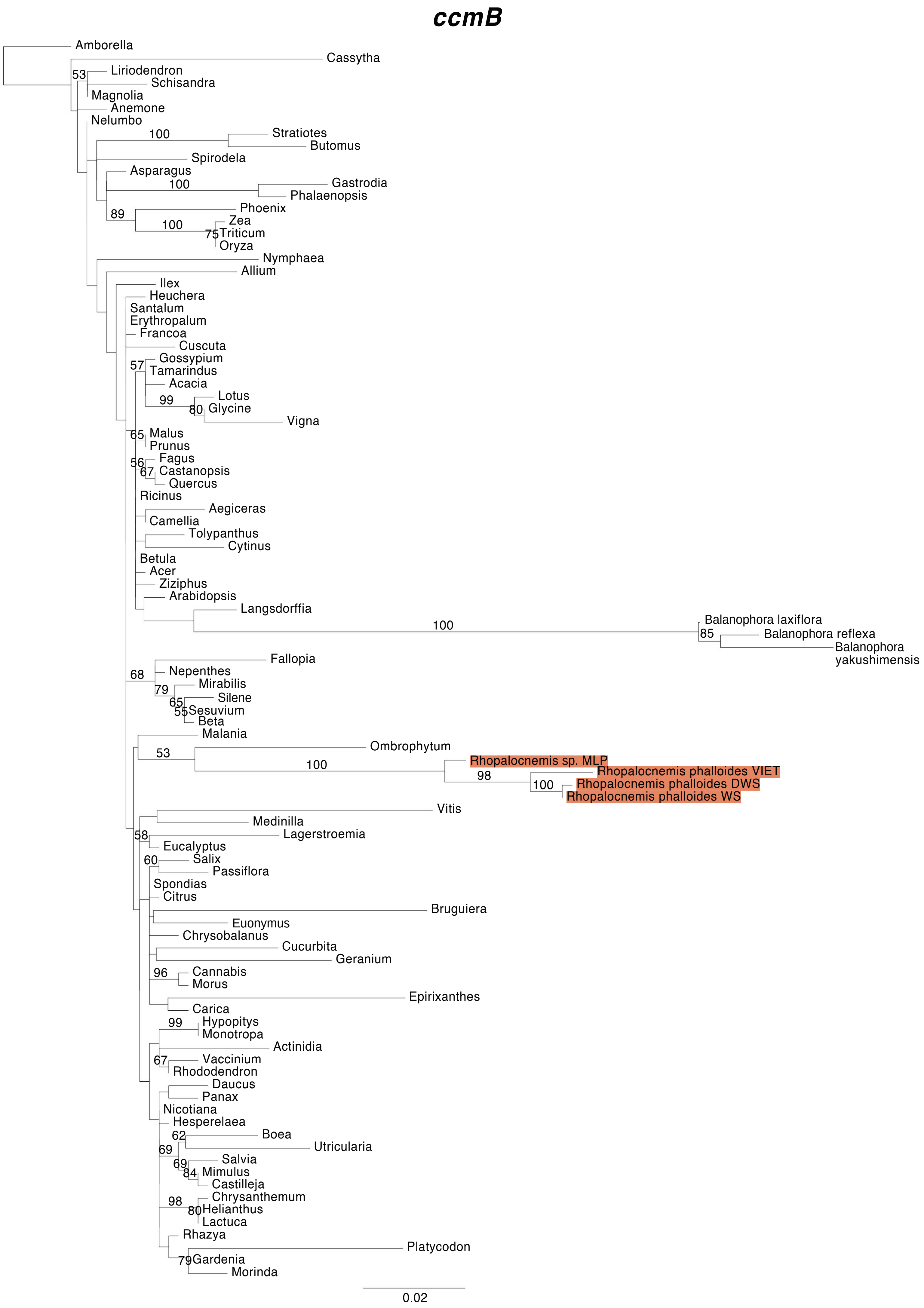


**Figure S6.** Maximum Likelihood (ML) phylogenetic analyses of protein-coding gene sequences in the mtDNAs of *Rhopalocnemis*. ML bootstrap support values ≥50% are shown. Scale bars correspond to substitutions per site.

#### *ccmC*

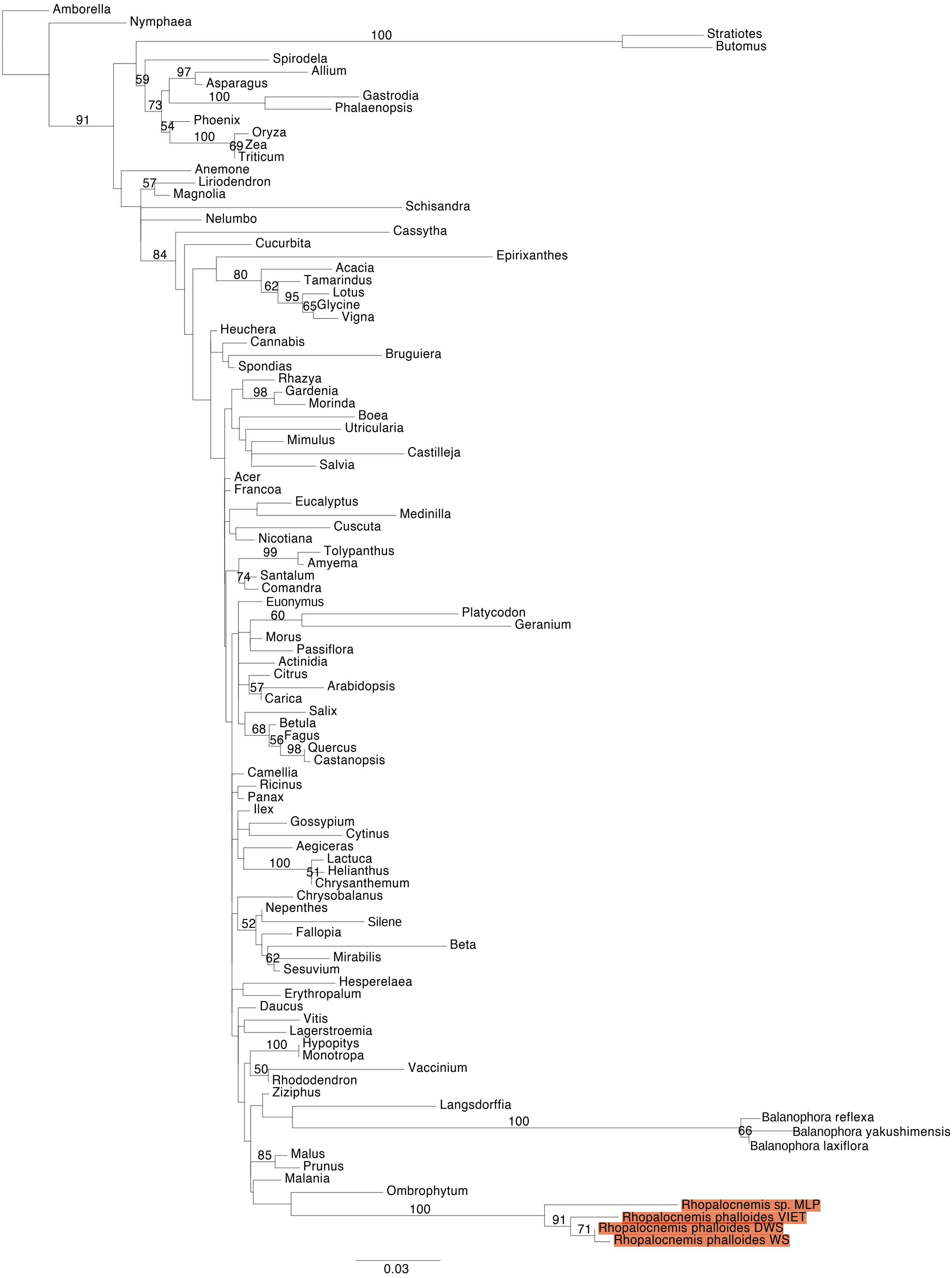


**Figure S6.** Maximum Likelihood (ML) phylogenetic analyses of protein-coding gene sequences in the mtDNAs of *Rhopalocnemis*. ML bootstrap support values ≥50% are shown. Scale bars correspond to substitutions per site.

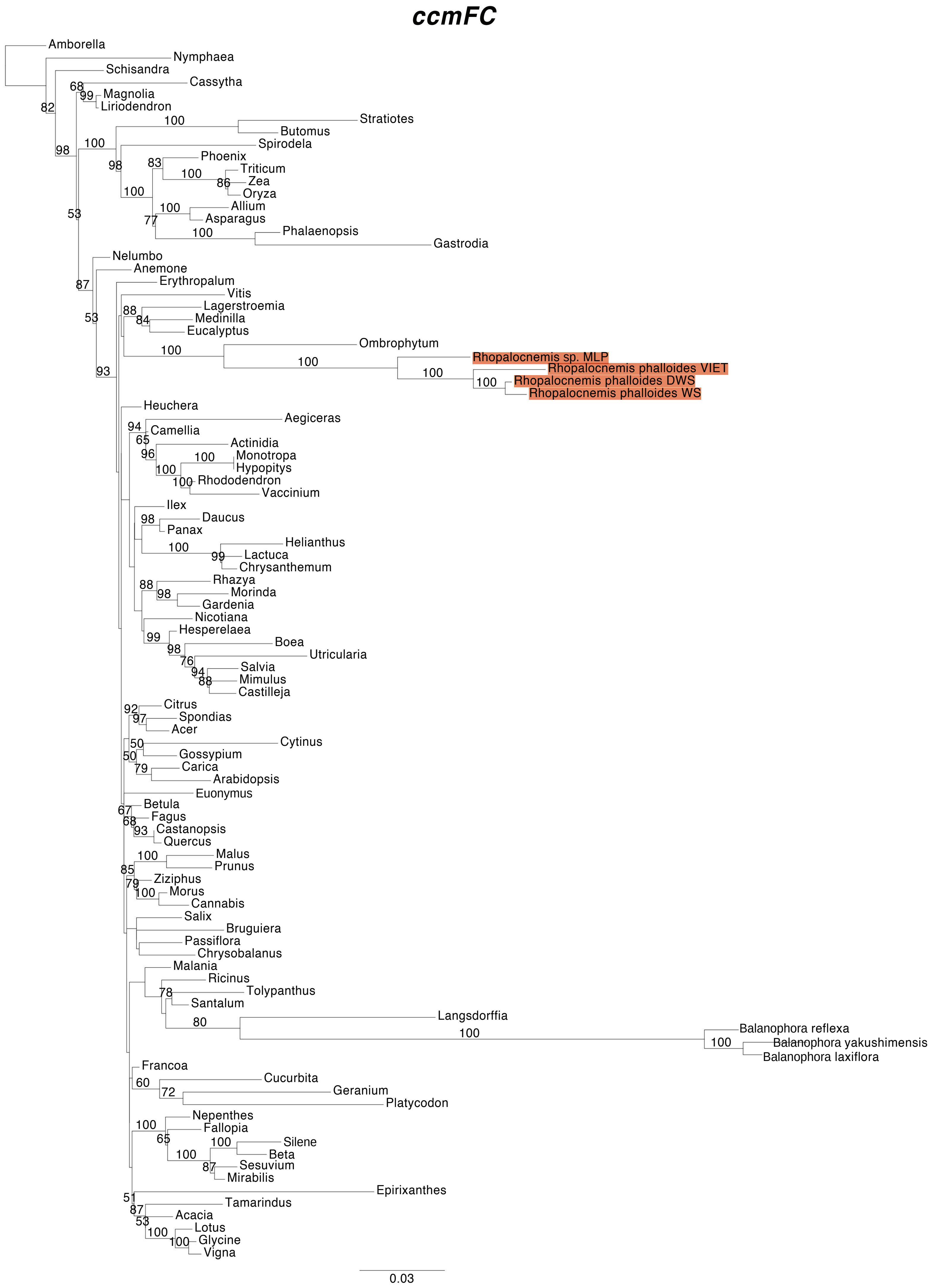


**Figure S6.** Maximum Likelihood (ML) phylogenetic analyses of protein-coding gene sequences in the mtDNAs of *Rhopalocnemis*. ML bootstrap support values ≥50% are shown. Scale bars correspond to substitutions per site.

### ccmFN

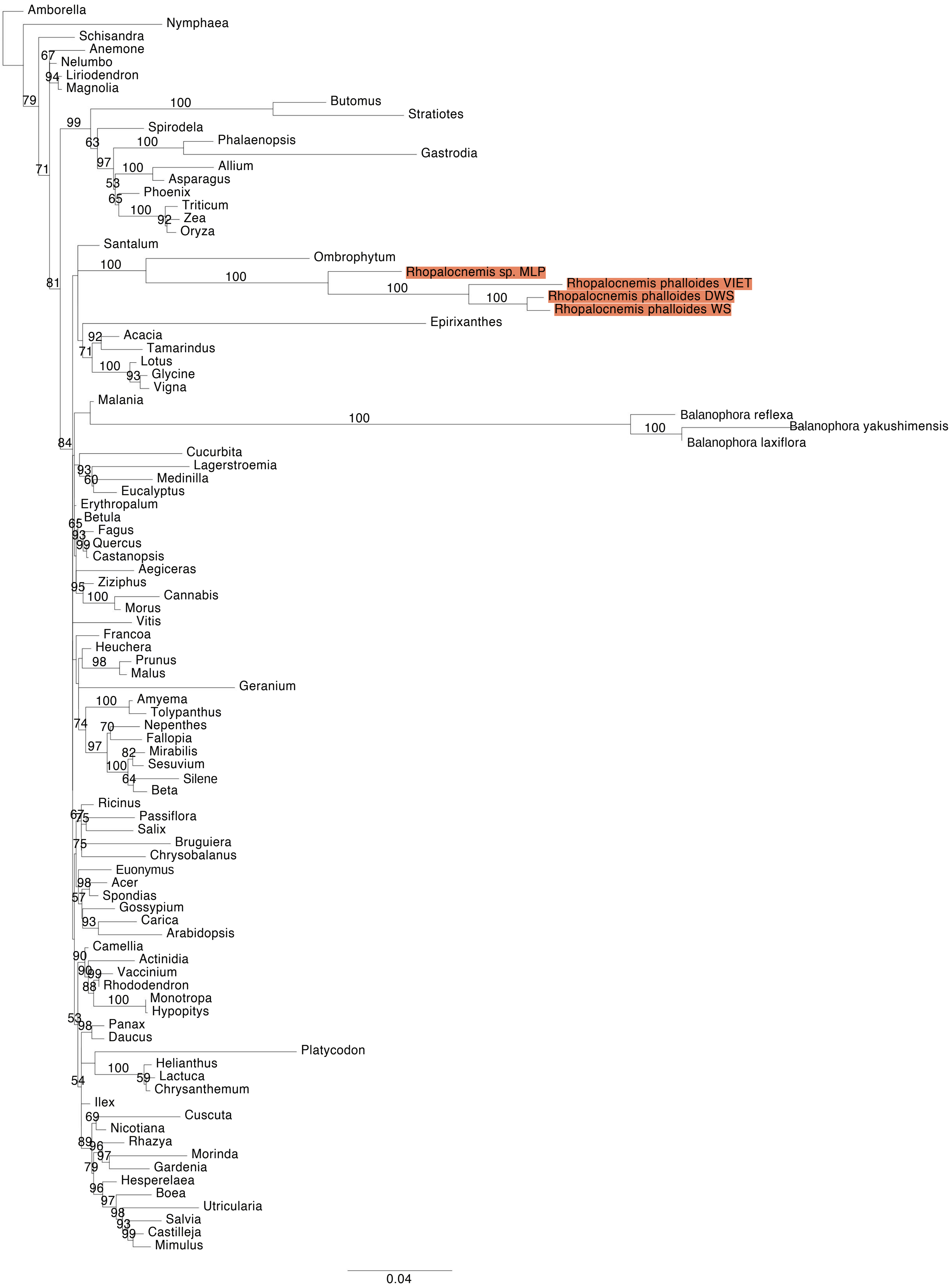


**Figure S6.** Maximum Likelihood (ML) phylogenetic analyses of protein-coding gene sequences in the mtDNAs of *Rhopalocnemis*. ML bootstrap support values ≥50% are shown. Scale bars correspond to substitutions per site.

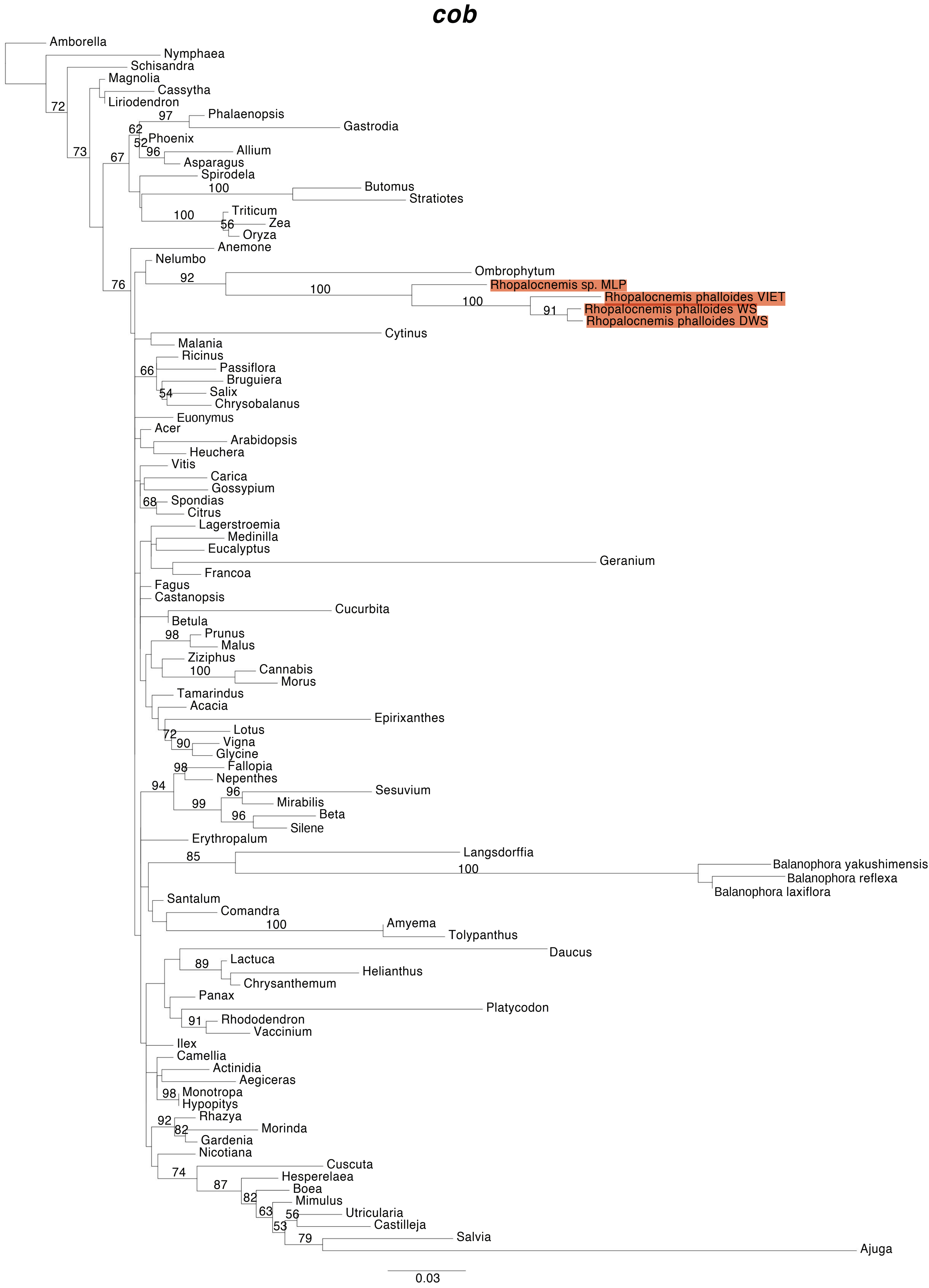


**Figure S6.** Maximum Likelihood (ML) phylogenetic analyses of protein-coding gene sequences in the mtDNAs of *Rhopalocnemis*. ML bootstrap support values ≥50% are shown. Scale bars correspond to substitutions per site.

***cox1***

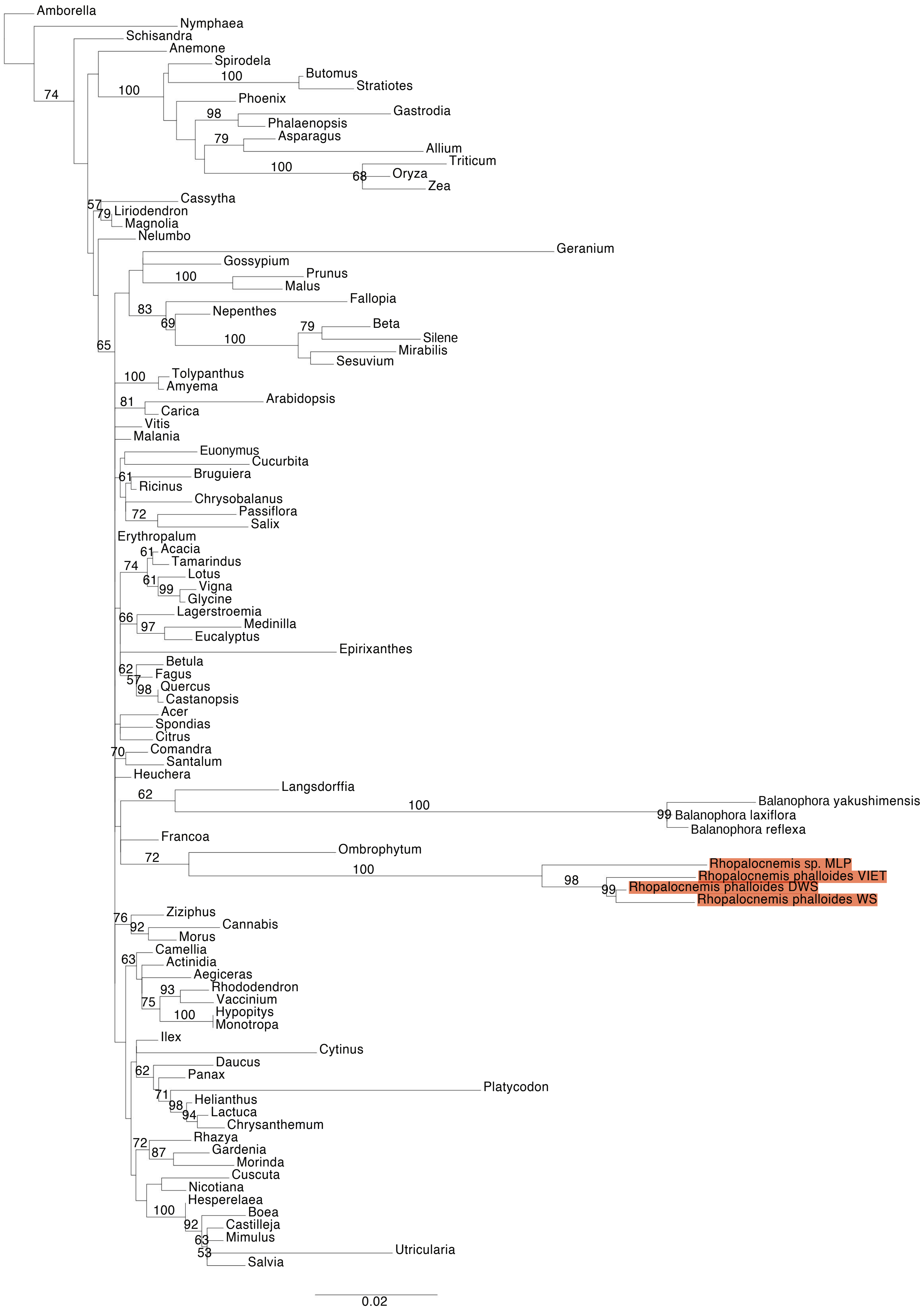


**Figure S6.** Maximum Likelihood (ML) phylogenetic analyses of protein-coding gene sequences in the mtDNAs of *Rhopalocnemis*. ML bootstrap support values ≥50% are shown. Scale bars correspond to substitutions per site.

### *cox2*

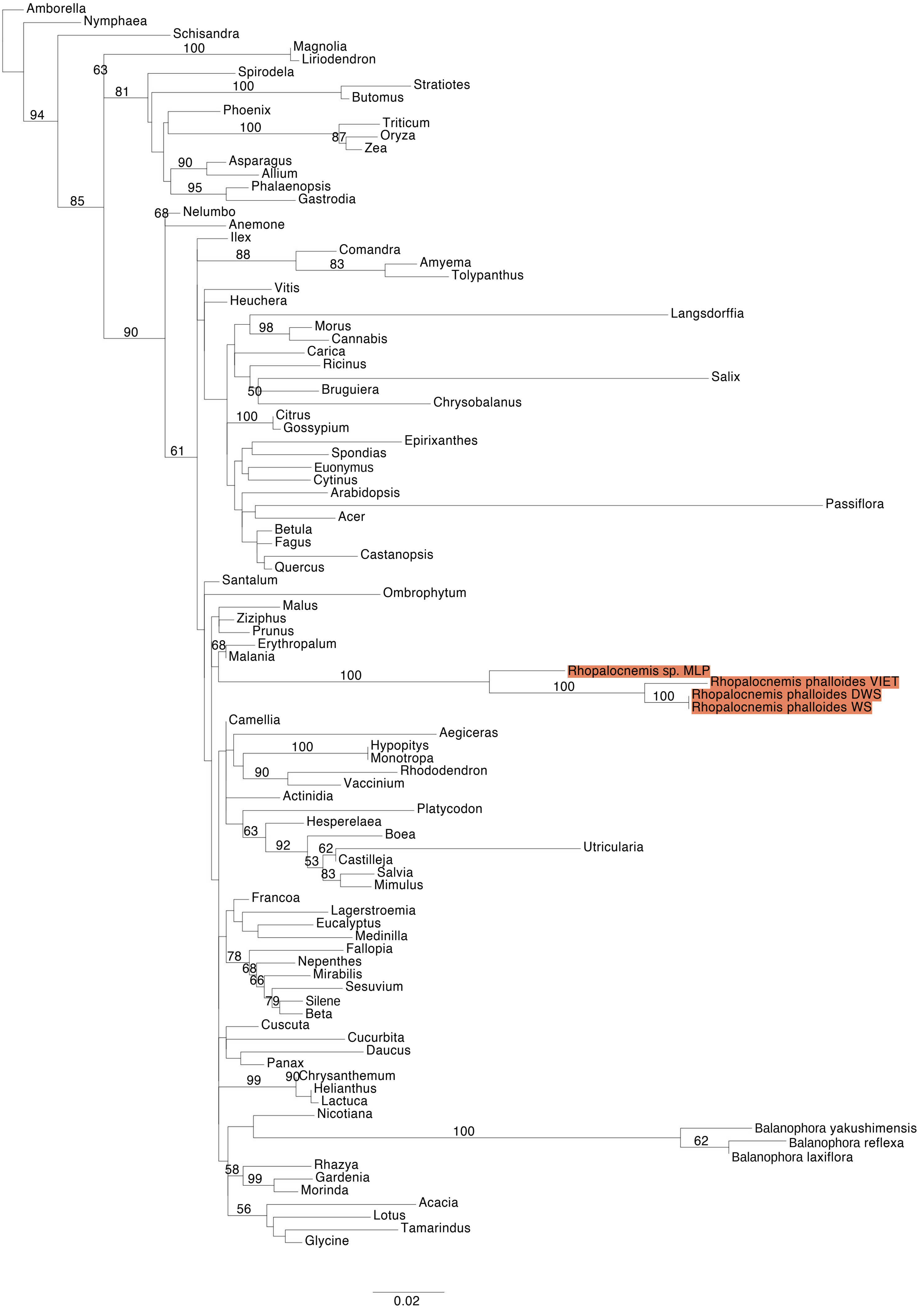


**Figure S6.** Maximum Likelihood (ML) phylogenetic analyses of protein-coding gene sequences in the mtDNAs of *Rhopalocnemis*. ML bootstrap support values ≥50% are shown. Scale bars correspond to substitutions per site.

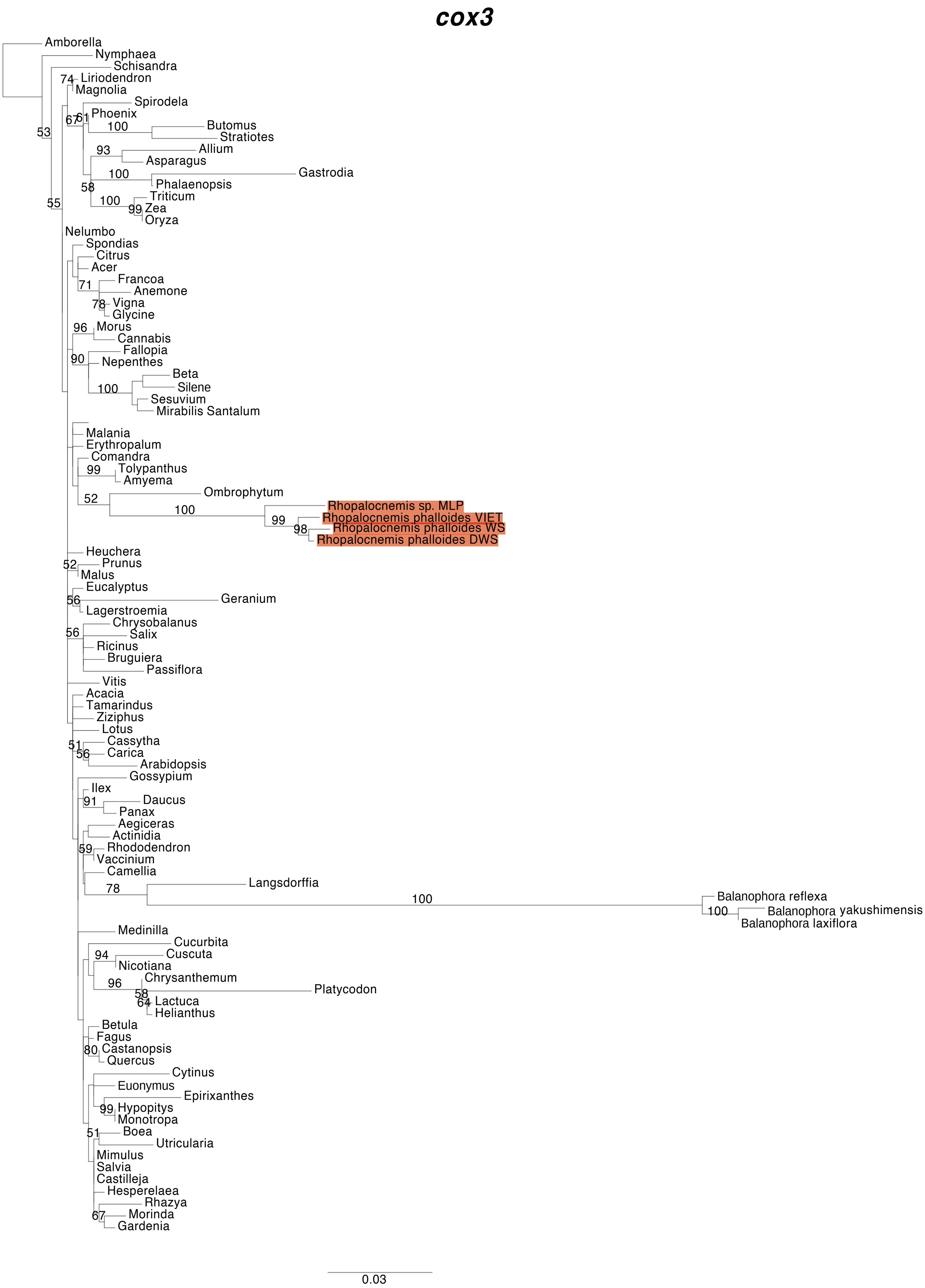


**Figure S6.** Maximum Likelihood (ML) phylogenetic analyses of protein-coding gene sequences in the mtDNAs of *Rhopalocnemis*. ML bootstrap support values ≥50% are shown. Scale bars correspond to substitutions per site.

#### *matR*

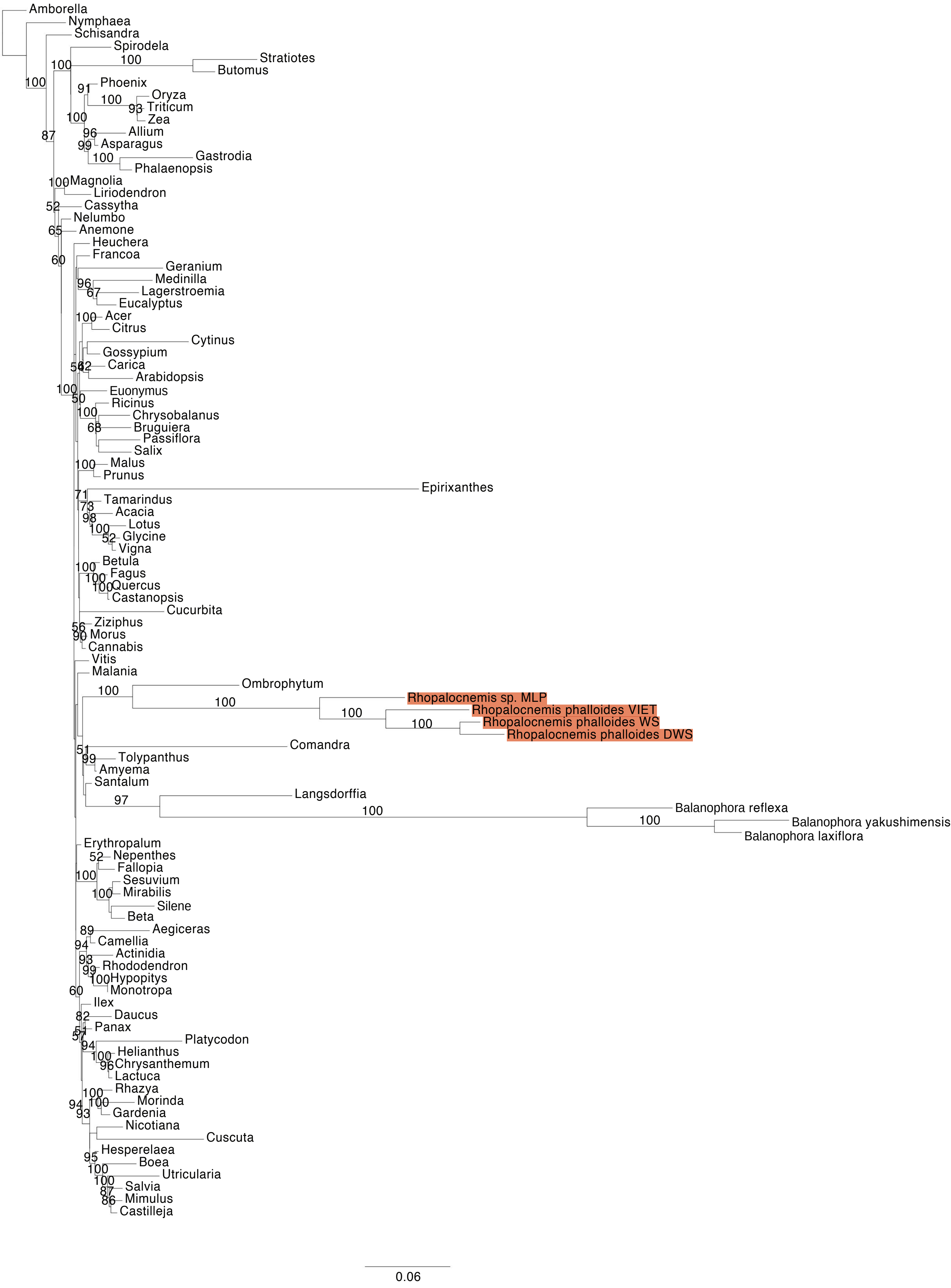


**Figure S6.** Maximum Likelihood (ML) phylogenetic analyses of protein-coding gene sequences in the mtDNAs of *Rhopalocnemis*. ML bootstrap support values ≥50% are shown. Scale bars correspond to substitutions per site.

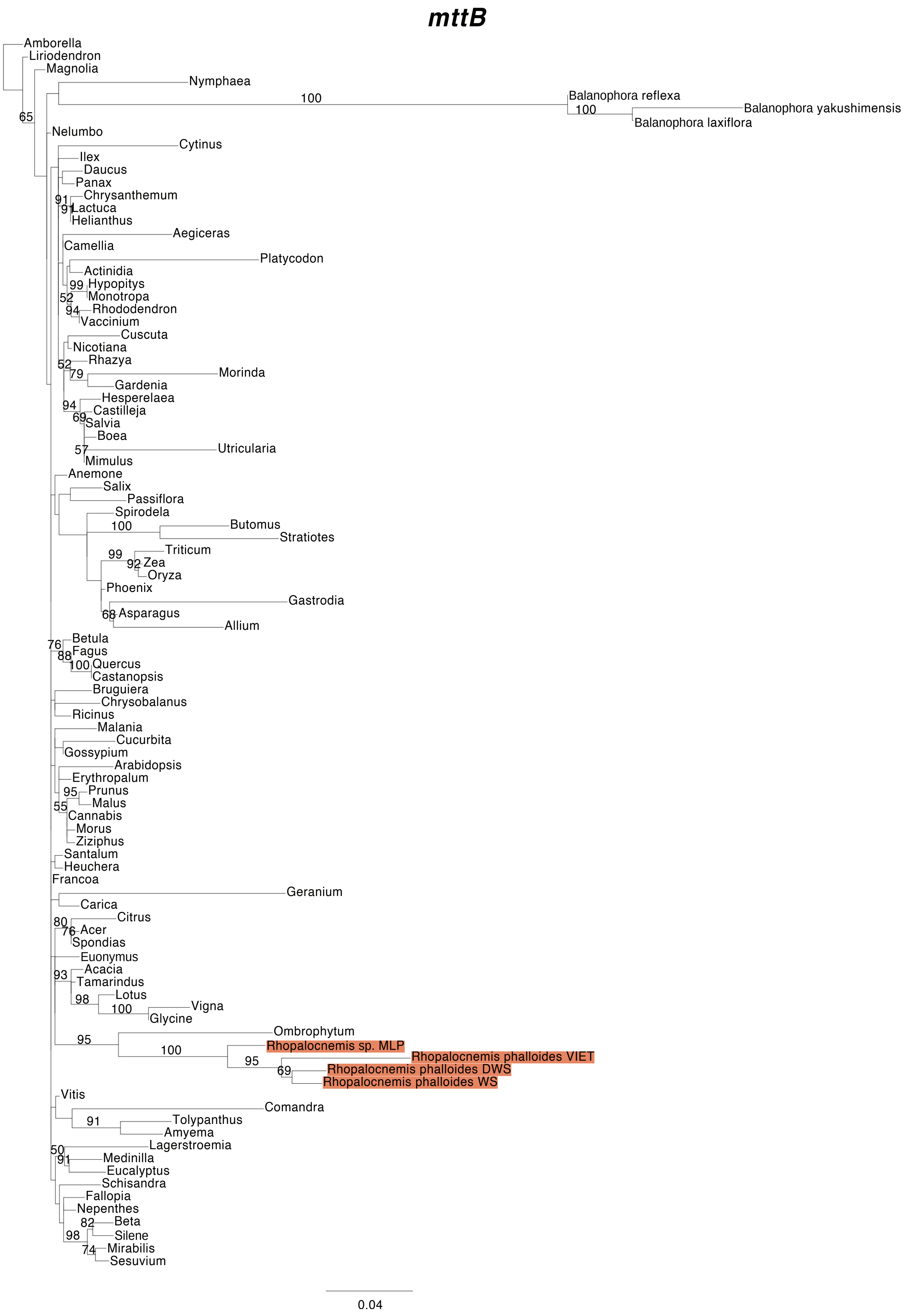


**Figure S6.** Maximum Likelihood (ML) phylogenetic analyses of protein-coding gene sequences in the mtDNAs of *Rhopalocnemis*. ML bootstrap support values ≥50% are shown. Scale bars correspond to substitutions per site.

#### *nad1*

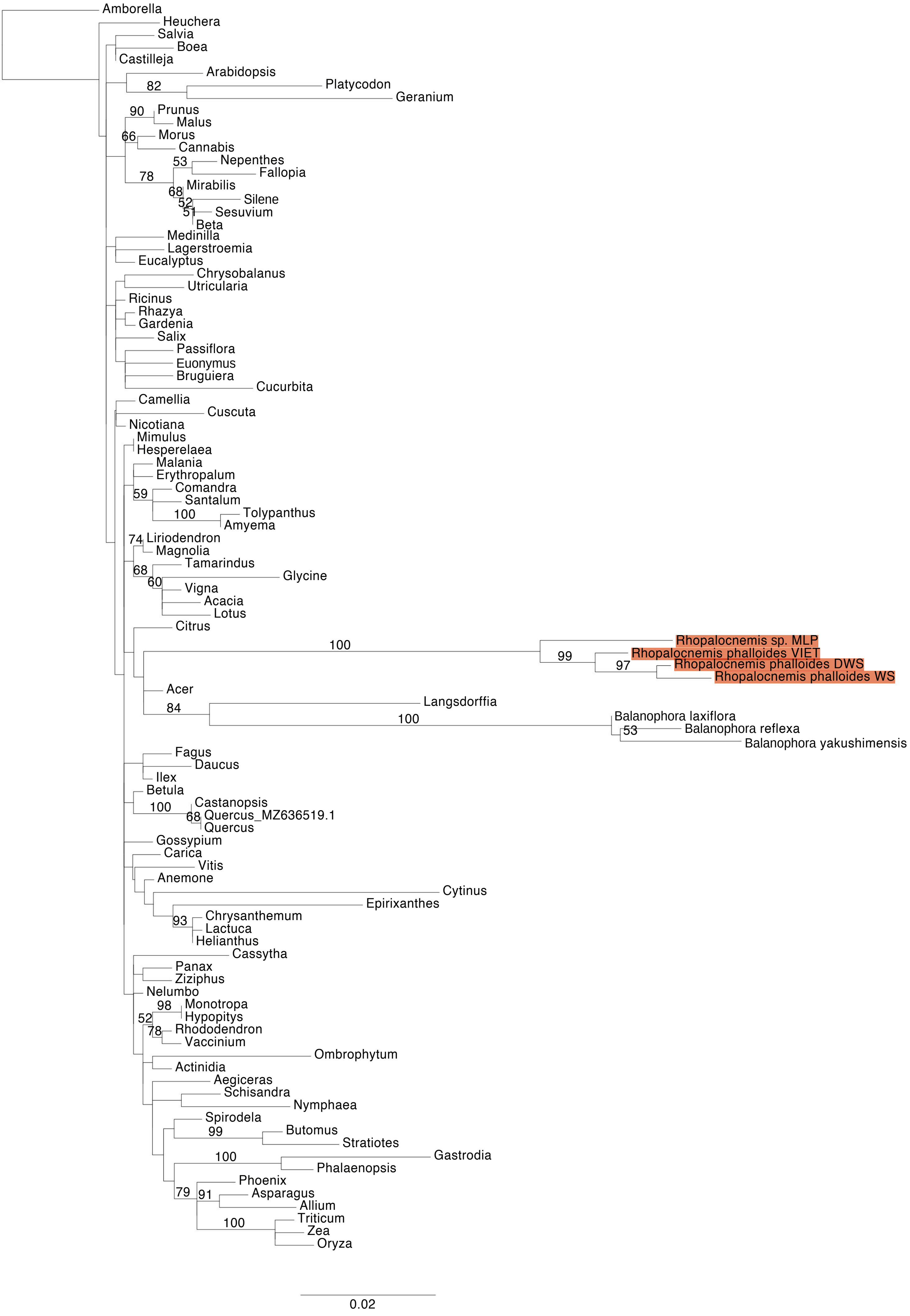


**Figure S6.** Maximum Likelihood (ML) phylogenetic analyses of protein-coding gene sequences in the mtDNAs of *Rhopalocnemis*. ML bootstrap support values ≥50% are shown. Scale bars correspond to substitutions per site.

#### *nad2*

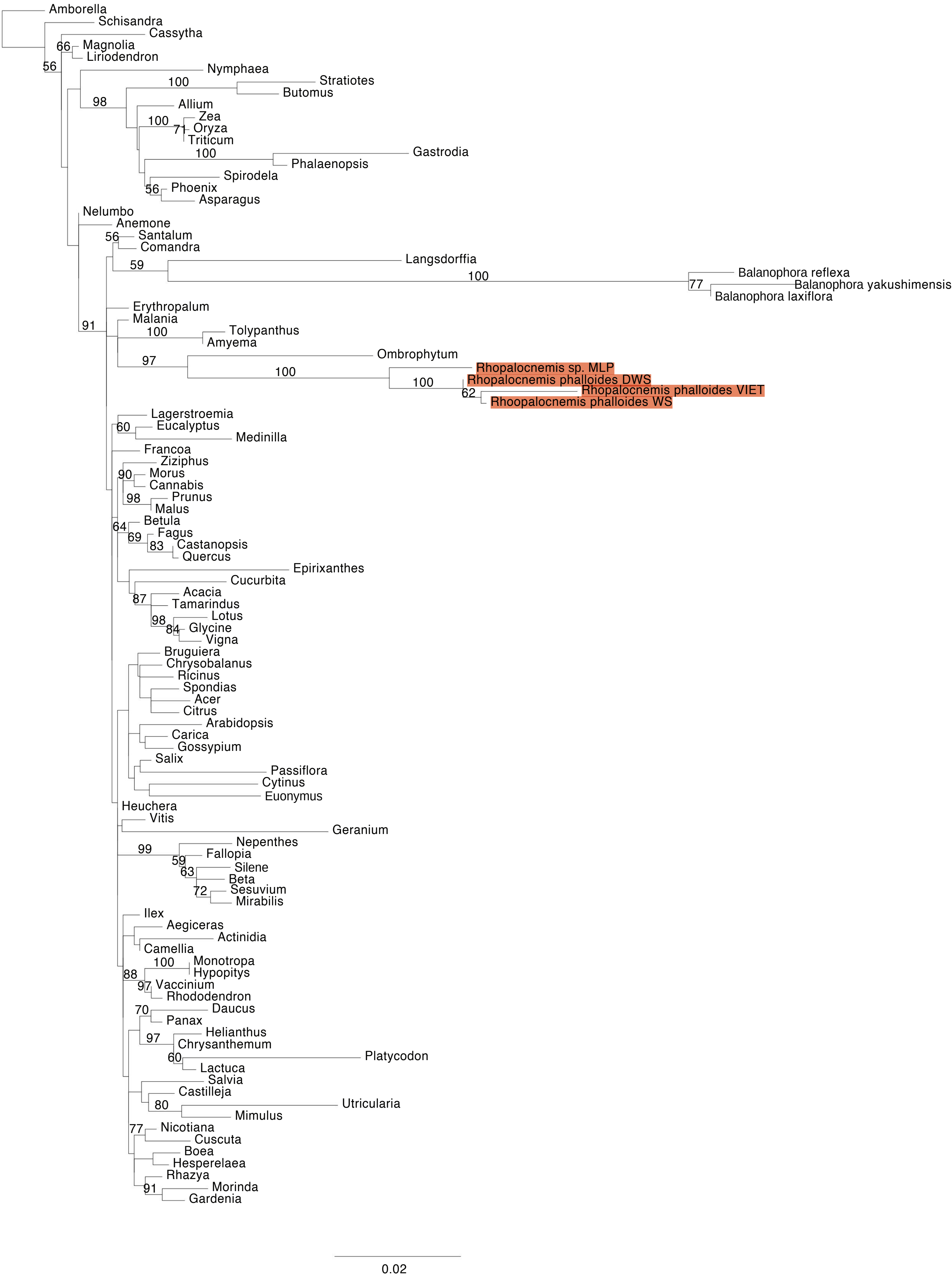


**Figure S6.** Maximum Likelihood (ML) phylogenetic analyses of protein-coding gene sequences in the mtDNAs of *Rhopalocnemis*. ML bootstrap support values ≥50% are shown. Scale bars correspond to substitutions per site.

### *nad3*

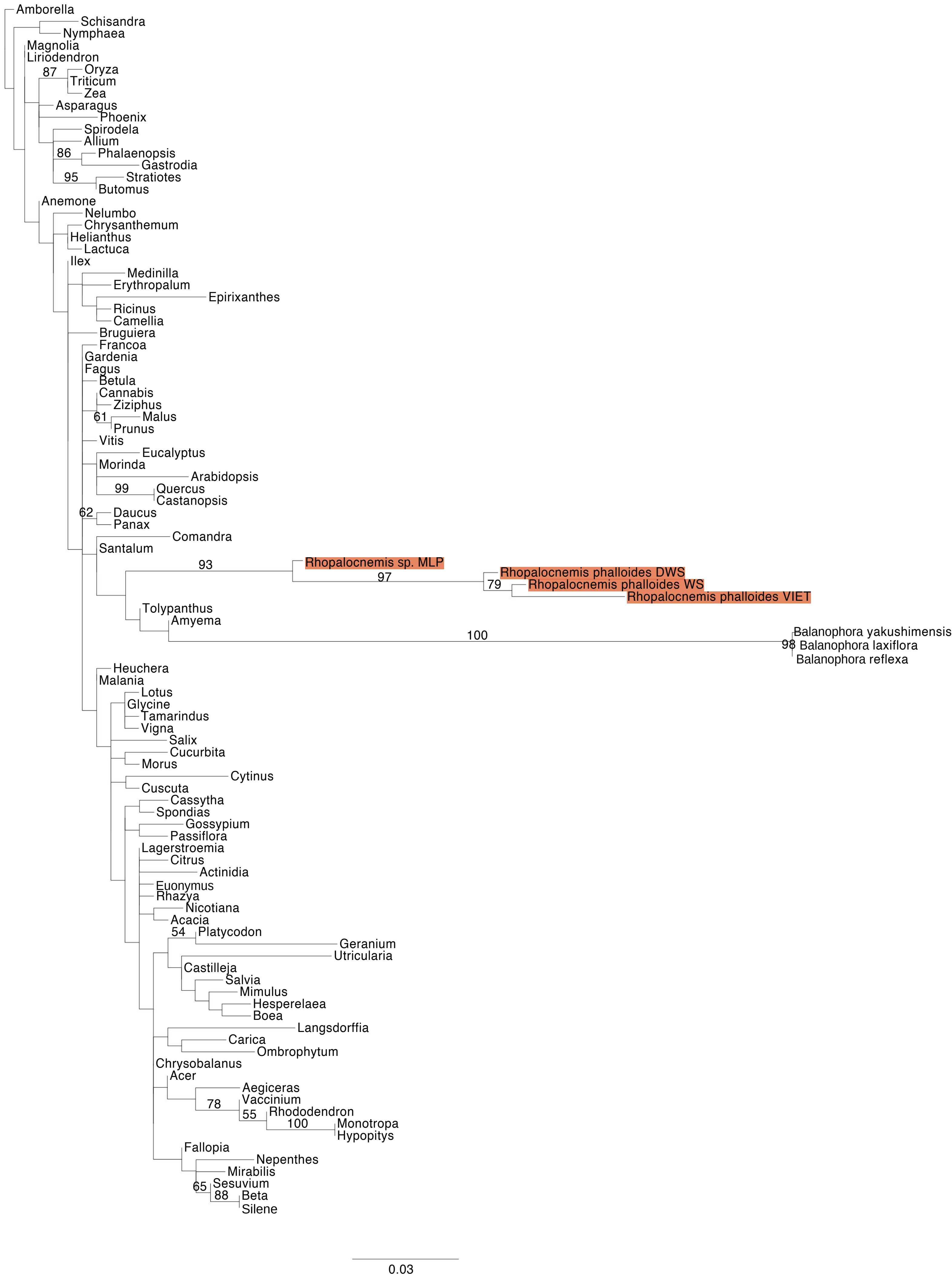


**Figure S6.** Maximum Likelihood (ML) phylogenetic analyses of protein-coding gene sequences in the mtDNAs of *Rhopalocnemis*. ML bootstrap support values ≥50% are shown. Scale bars correspond to substitutions per site.

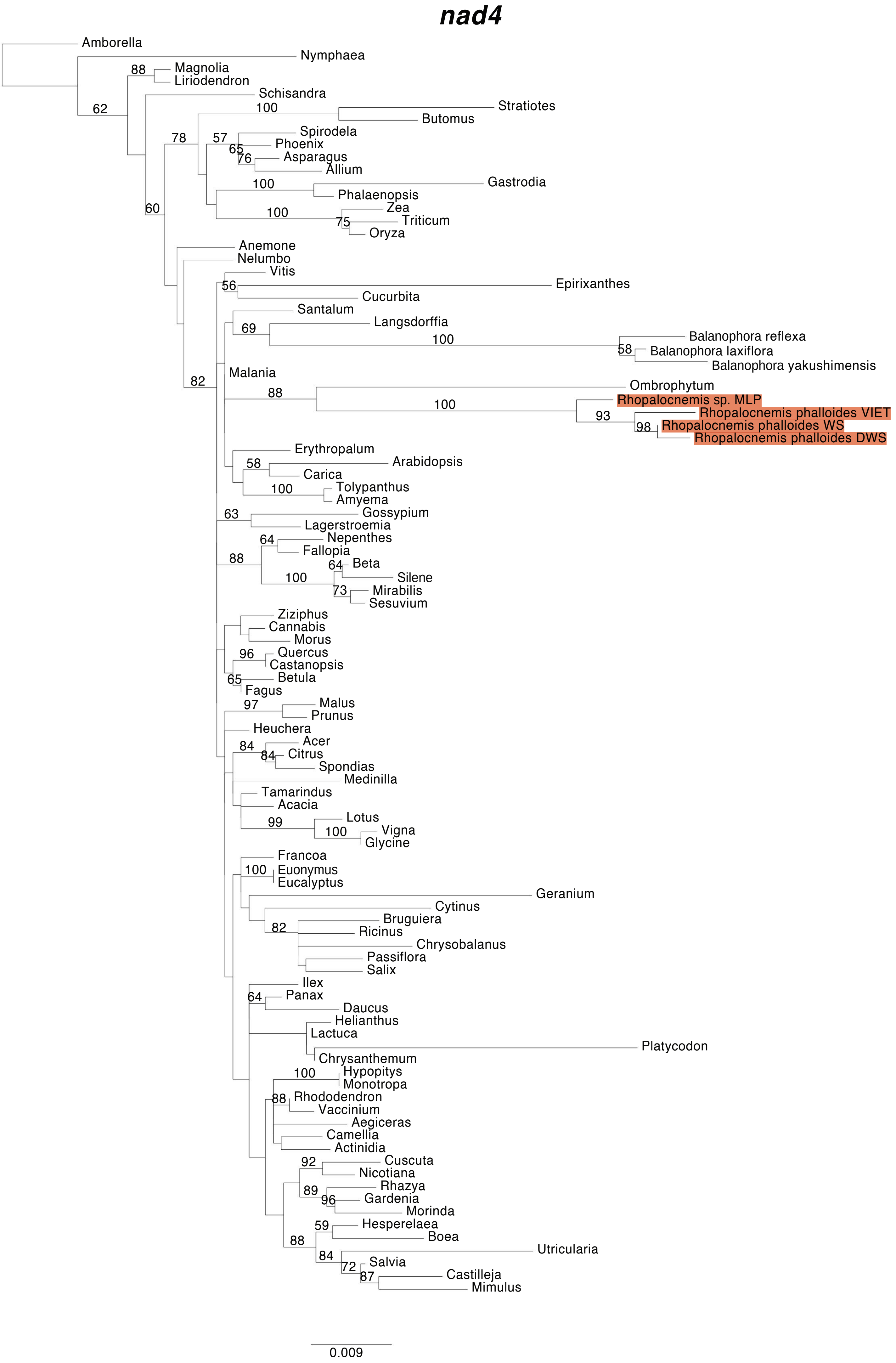


**Figure S6.** Maximum Likelihood (ML) phylogenetic analyses of protein-coding gene sequences in the mtDNAs of *Rhopalocnemis*. ML bootstrap support values ≥50% are shown. Scale bars correspond to substitutions per site.

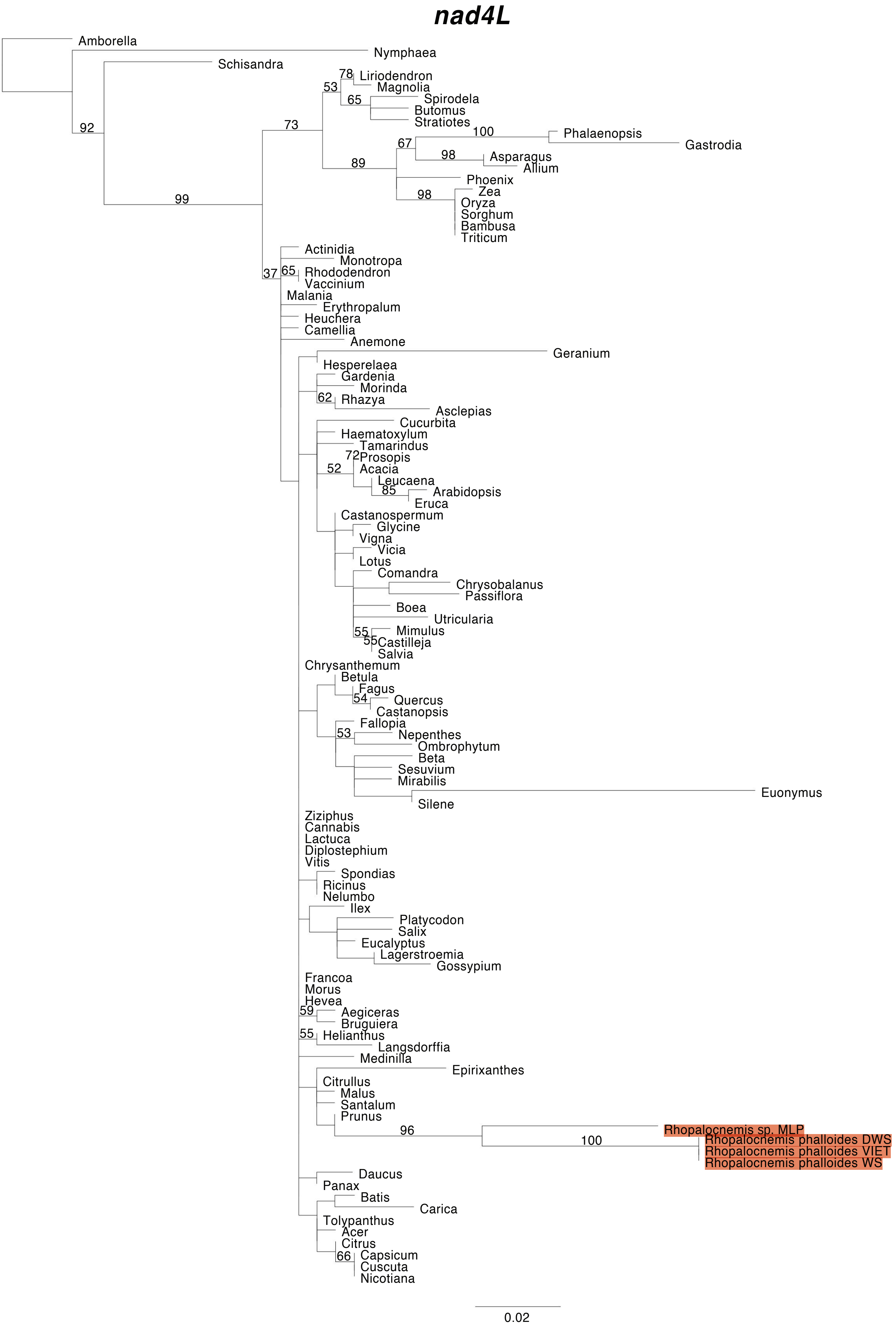


**Figure S6.** Maximum Likelihood (ML) phylogenetic analyses of protein-coding gene sequences in the mtDNAs of *Rhopalocnemis*. ML bootstrap support values ≥50% are shown. Scale bars correspond to substitutions per site.

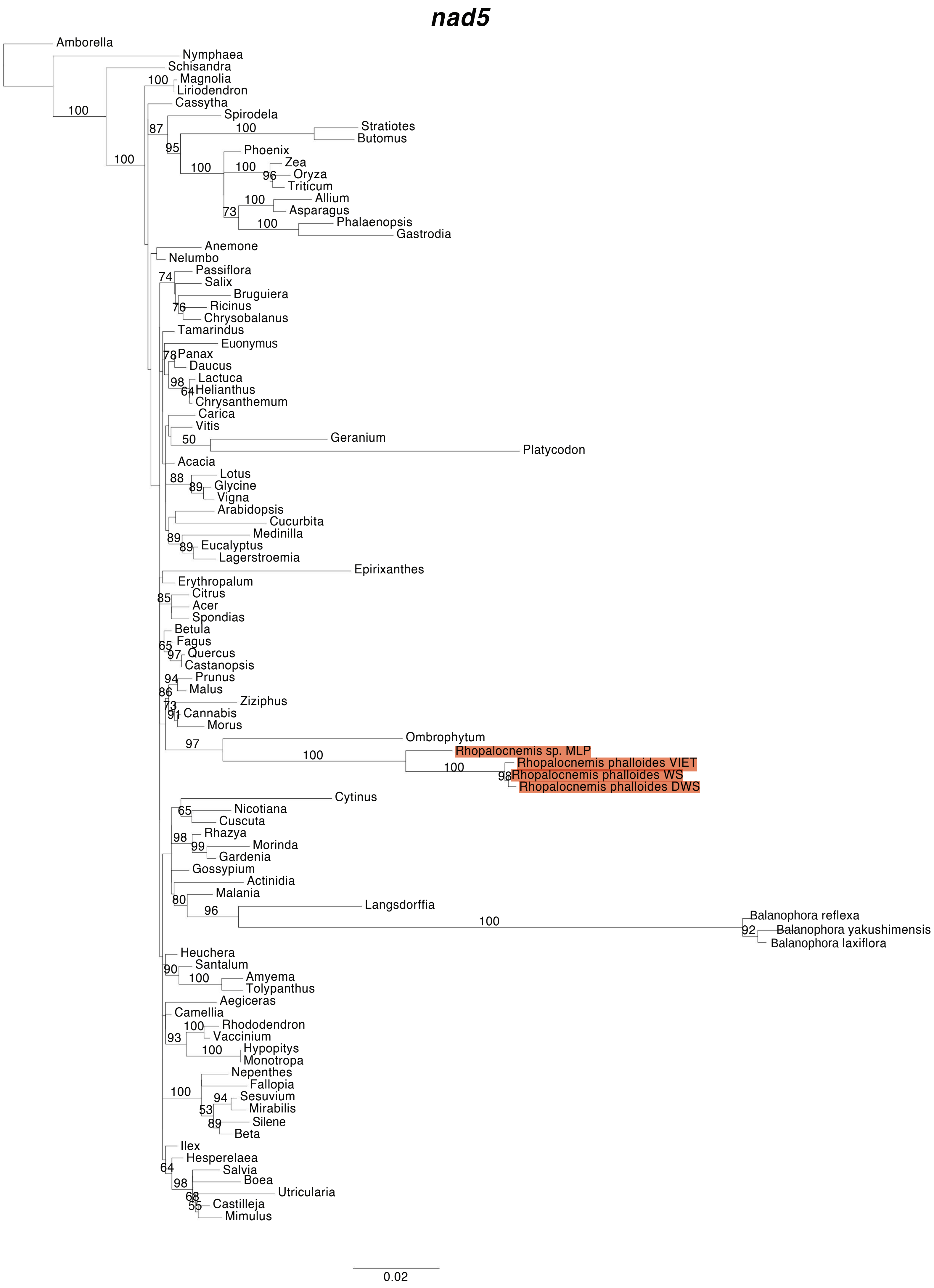


**Figure S6.** Maximum Likelihood (ML) phylogenetic analyses of protein-coding gene sequences in the mtDNAs of *Rhopalocnemis*. ML bootstrap support values ≥50% are shown. Scale bars correspond to substitutions per site.

***nad6***

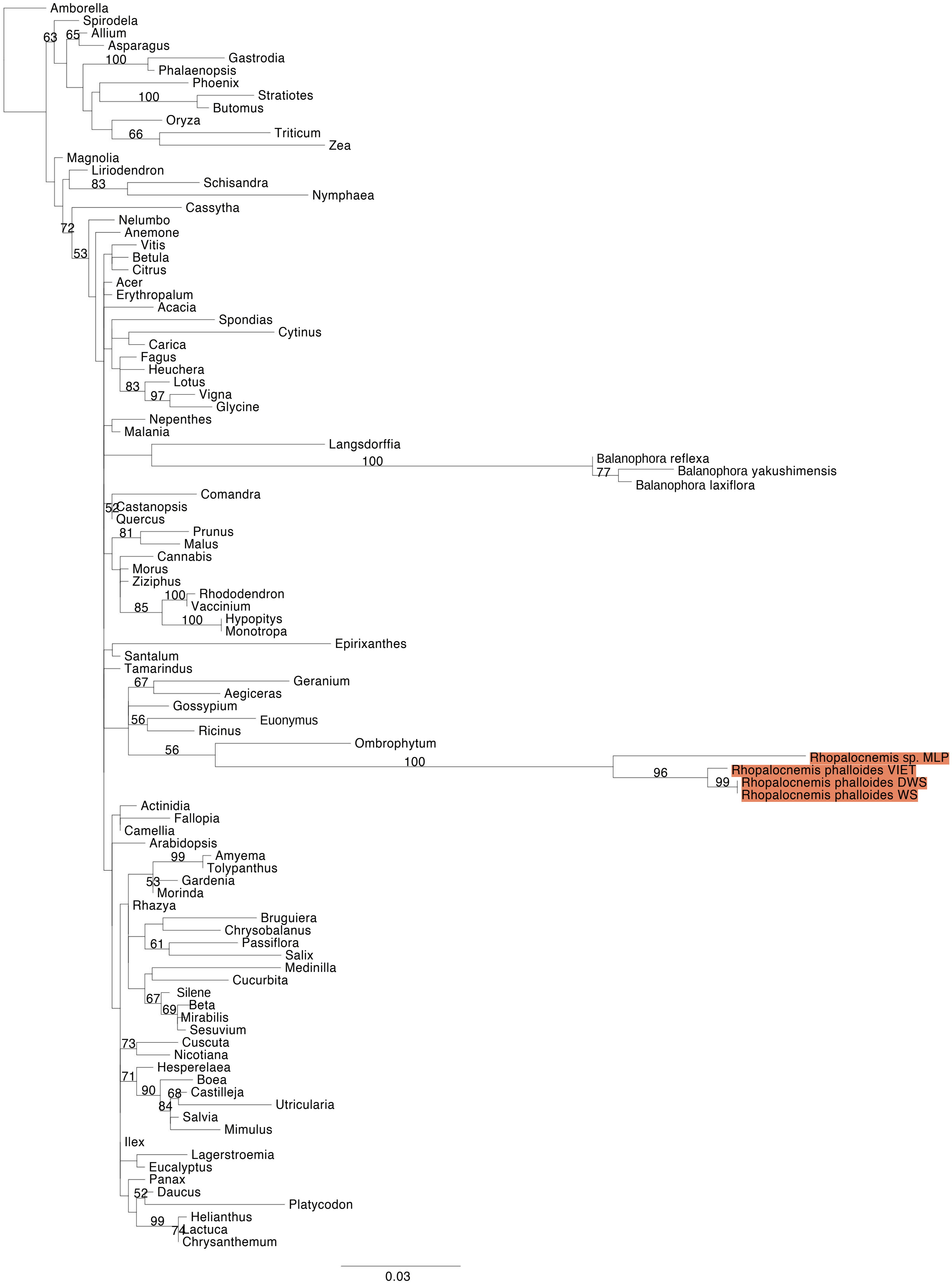


**Figure S6.** Maximum Likelihood (ML) phylogenetic analyses of protein-coding gene sequences in the mtDNAs of *Rhopalocnemis*. ML bootstrap support values ≥50% are shown. Scale bars correspond to substitutions per site.

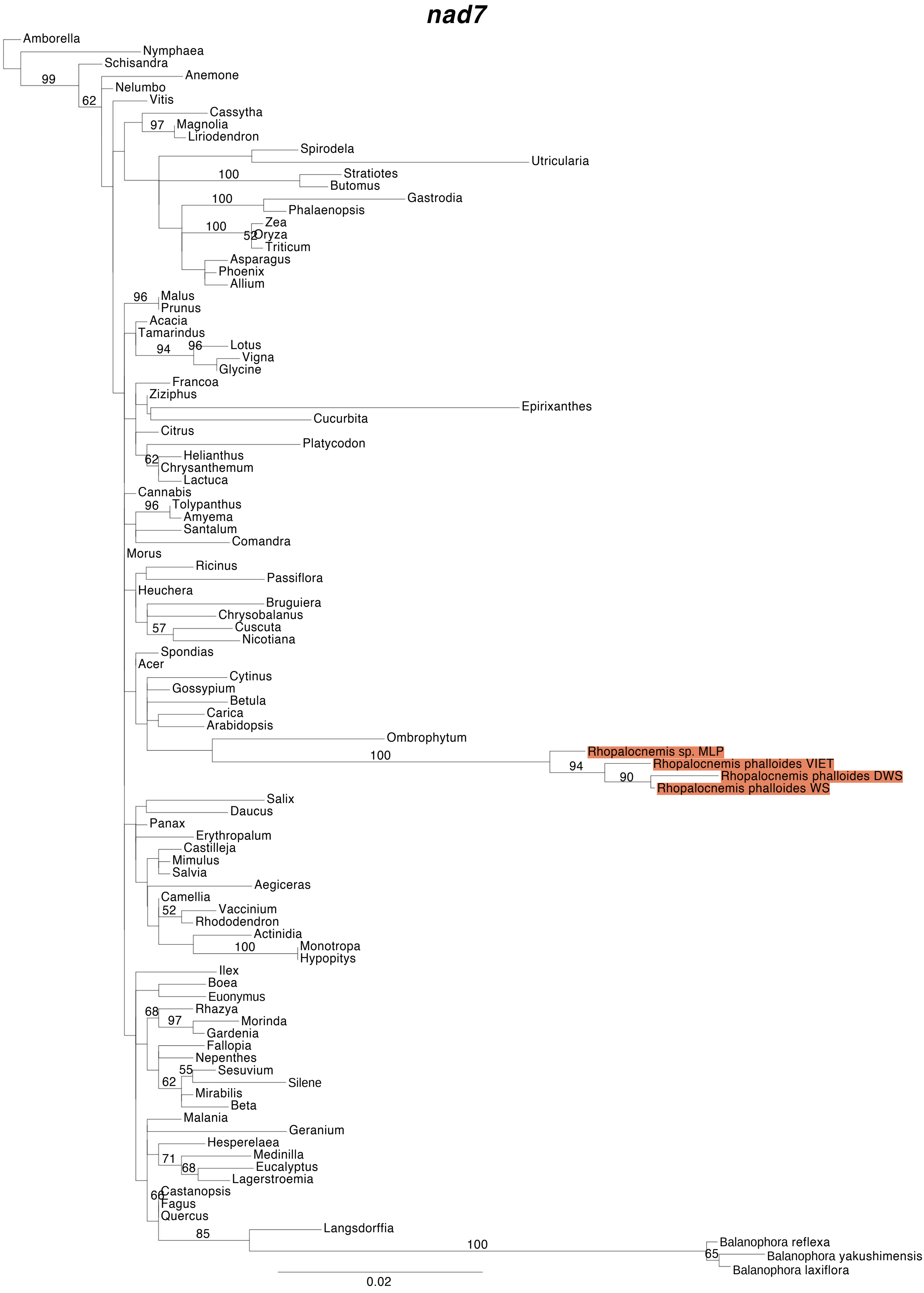


**Figure S6.** Maximum Likelihood (ML) phylogenetic analyses of protein-coding gene sequences in the mtDNAs of *Rhopalocnemis*. ML bootstrap support values ≥50% are shown. Scale bars correspond to substitutions per site.

***nad9***

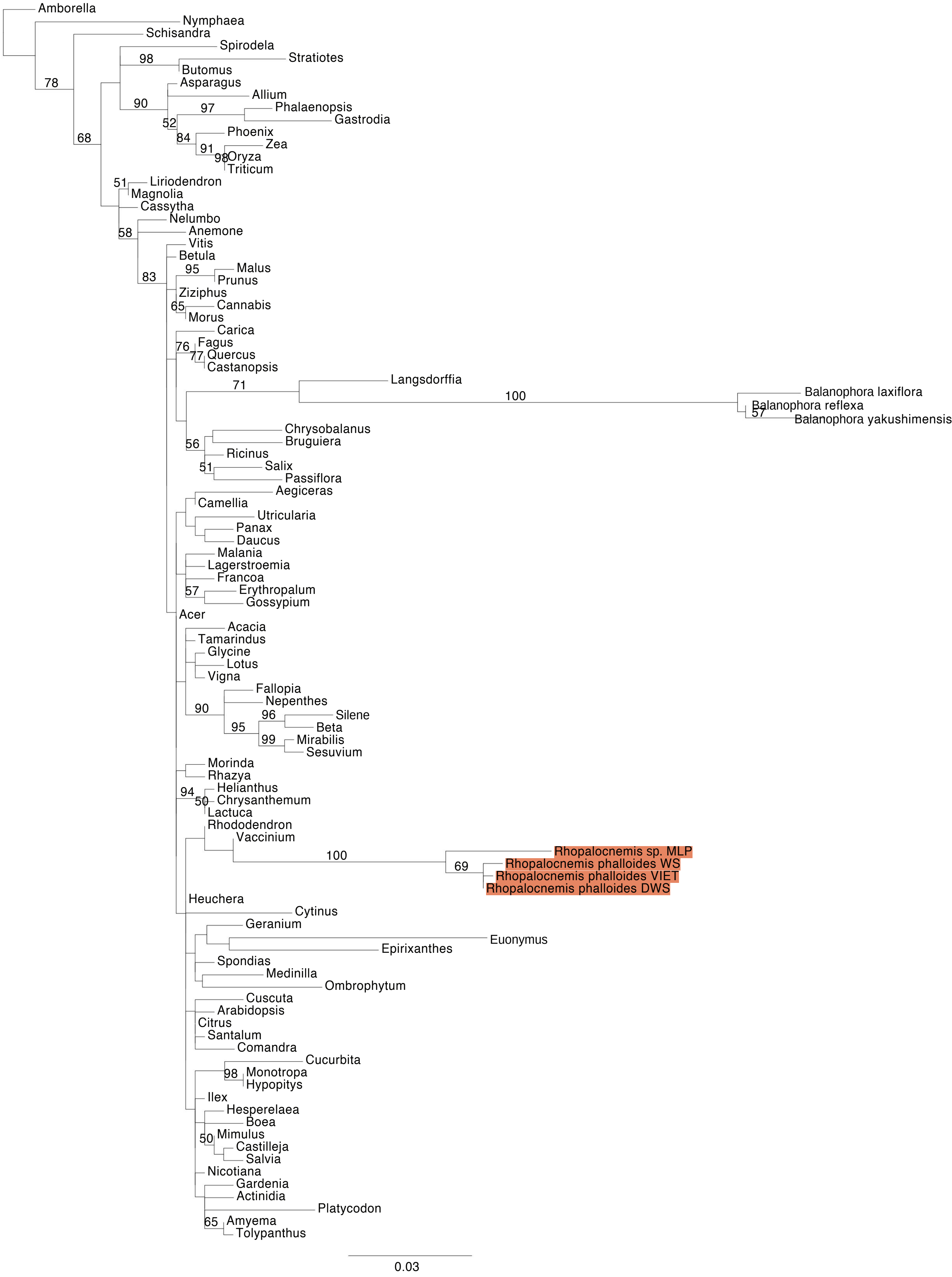


**Figure S6.** Maximum Likelihood (ML) phylogenetic analyses of protein-coding gene sequences in the mtDNAs of *Rhopalocnemis*. ML bootstrap support values ≥50% are shown. Scale bars correspond to substitutions per site.

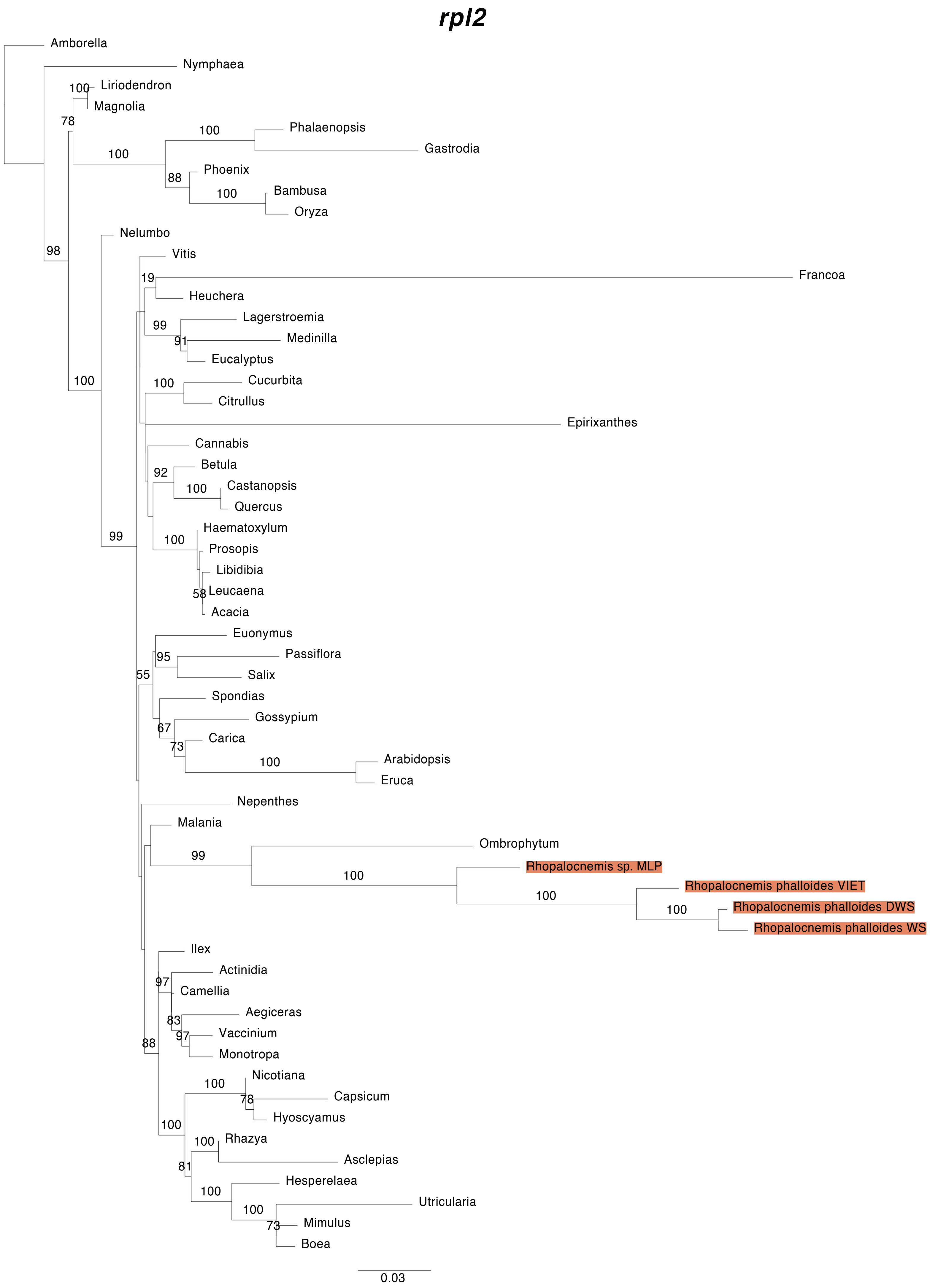


**Figure S6.** Maximum Likelihood (ML) phylogenetic analyses of protein-coding gene sequences in the mtDNAs of *Rhopalocnemis*. ML bootstrap support values ≥50% are shown. Scale bars correspond to substitutions per site.

### *rpl5*

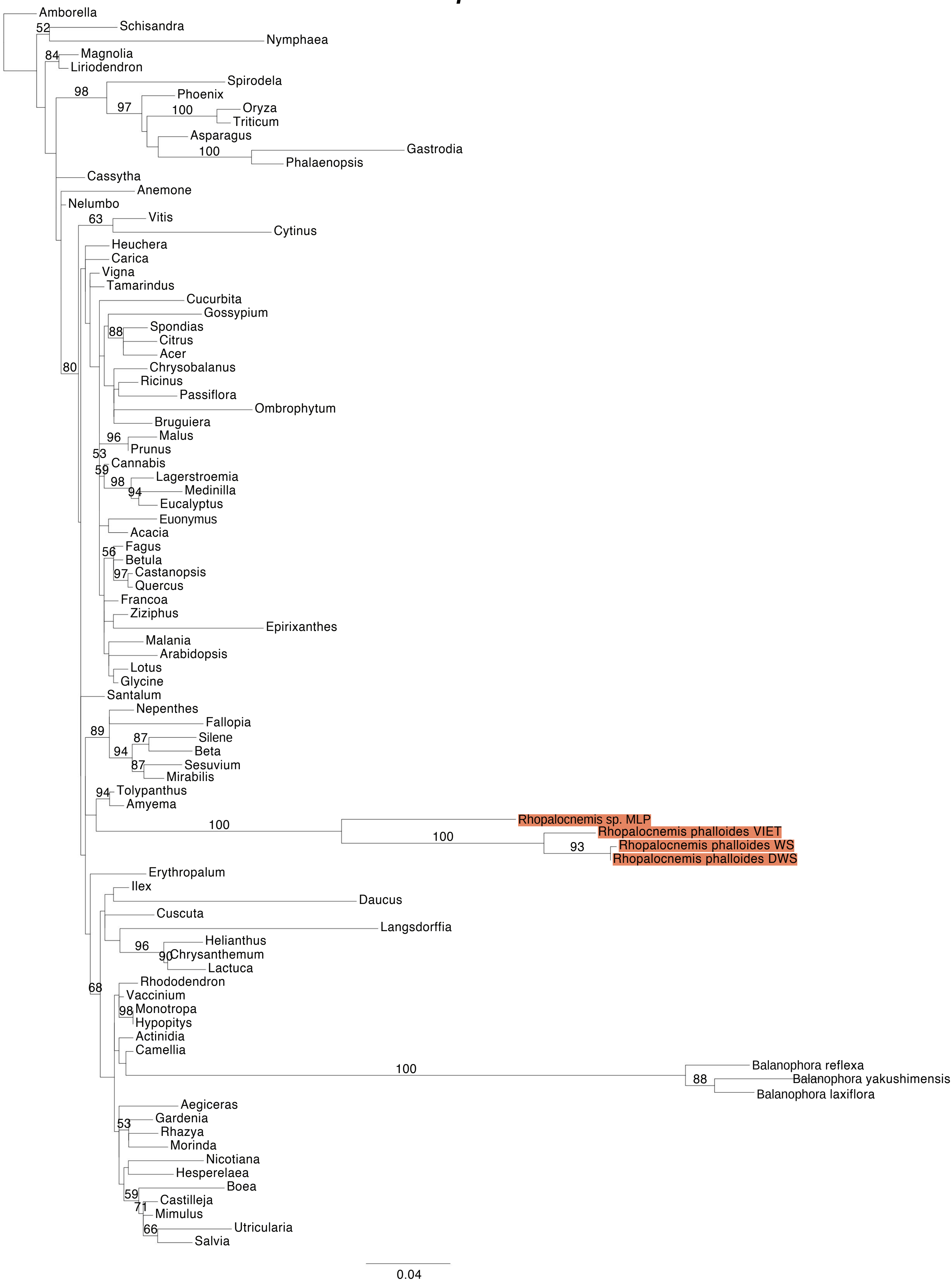


**Figure S6.** Maximum Likelihood (ML) phylogenetic analyses of protein-coding gene sequences in the mtDNAs of *Rhopalocnemis*. ML bootstrap support values ≥50% are shown. Scale bars correspond to substitutions per site.

### *rpl10*

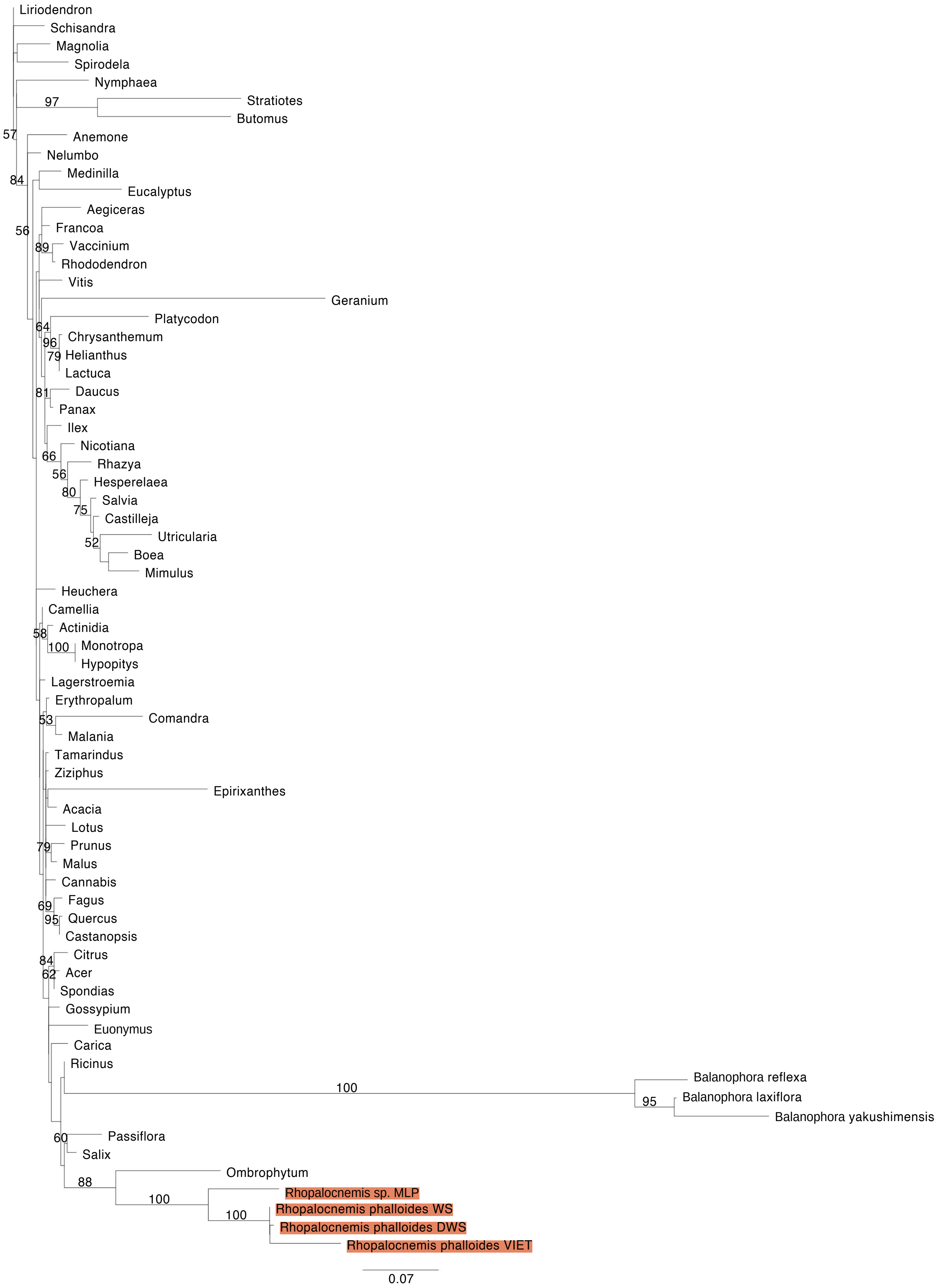


**Figure S6.** Maximum Likelihood (ML) phylogenetic analyses of protein-coding gene sequences in the mtDNAs of *Rhopalocnemis*. ML bootstrap support values ≥50% are shown. Scale bars correspond to substitutions per site.

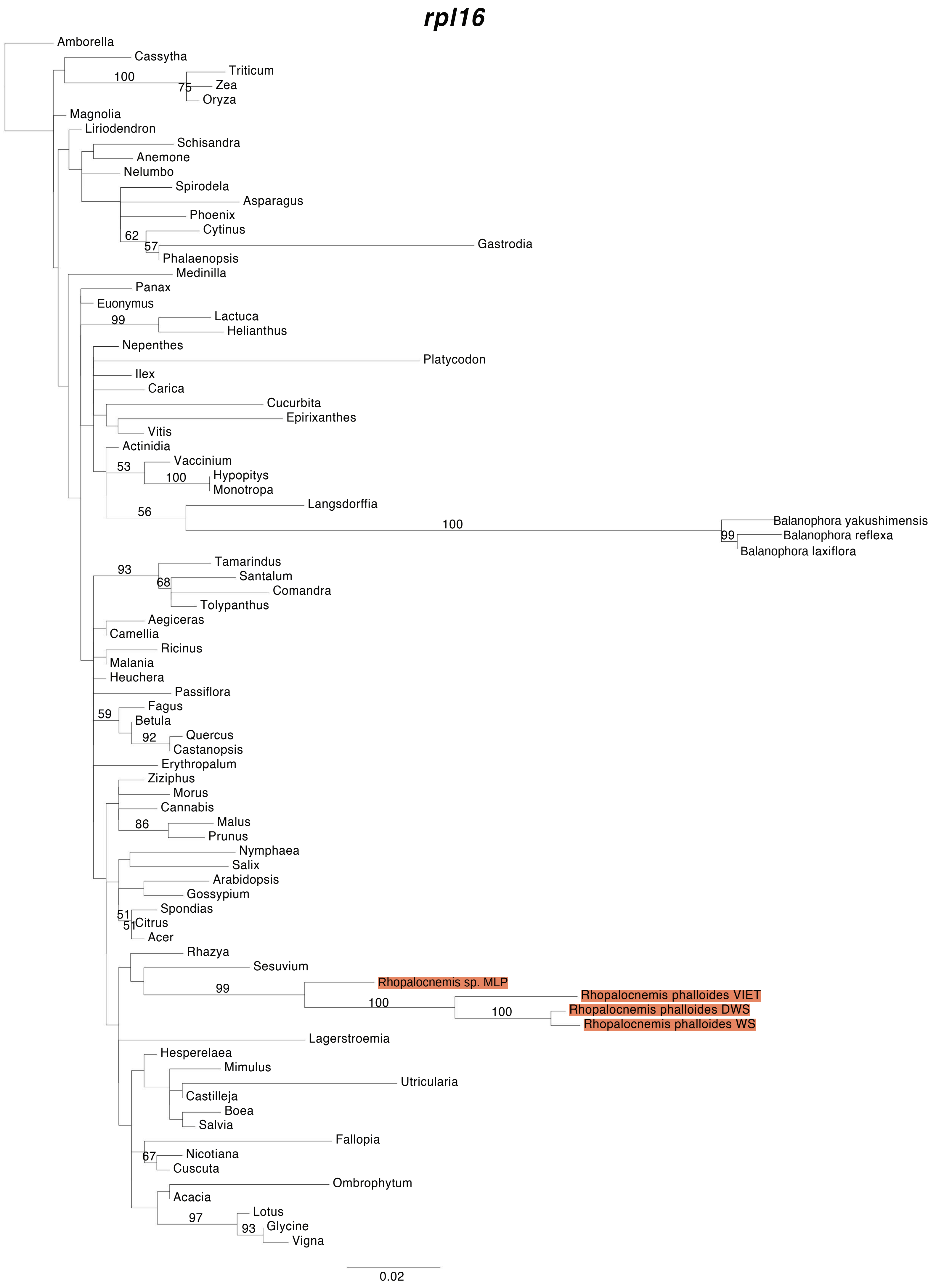


**Figure S6.** Maximum Likelihood (ML) phylogenetic analyses of protein-coding gene sequences in the mtDNAs of *Rhopalocnemis*. ML bootstrap support values ≥50% are shown. Scale bars correspond to substitutions per site.

### *rps1*

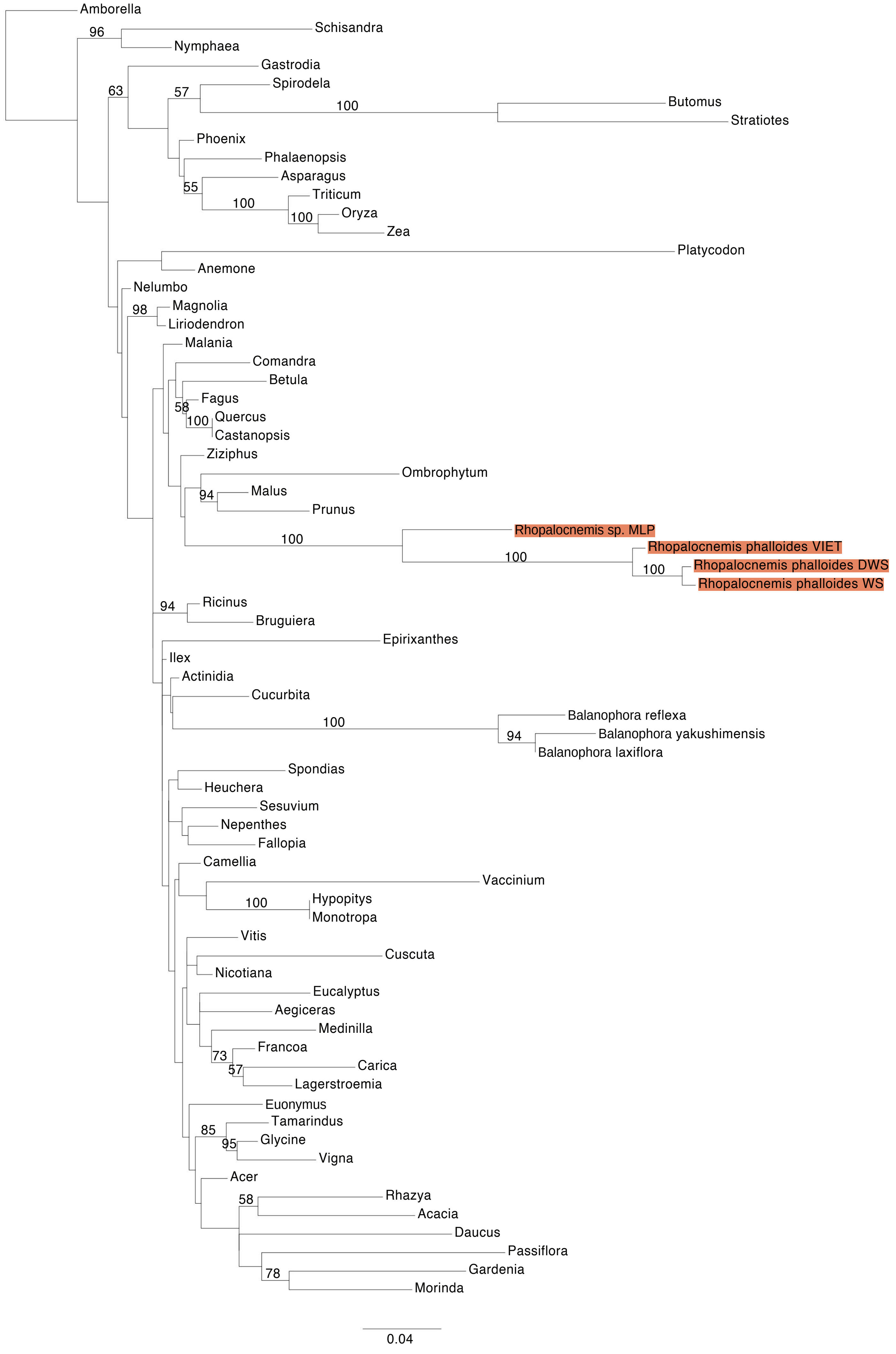


**Figure S6.** Maximum Likelihood (ML) phylogenetic analyses of protein-coding gene sequences in the mtDNAs of *Rhopalocnemis*. ML bootstrap support values ≥50% are shown. Scale bars correspond to substitutions per site.

### *rps3*

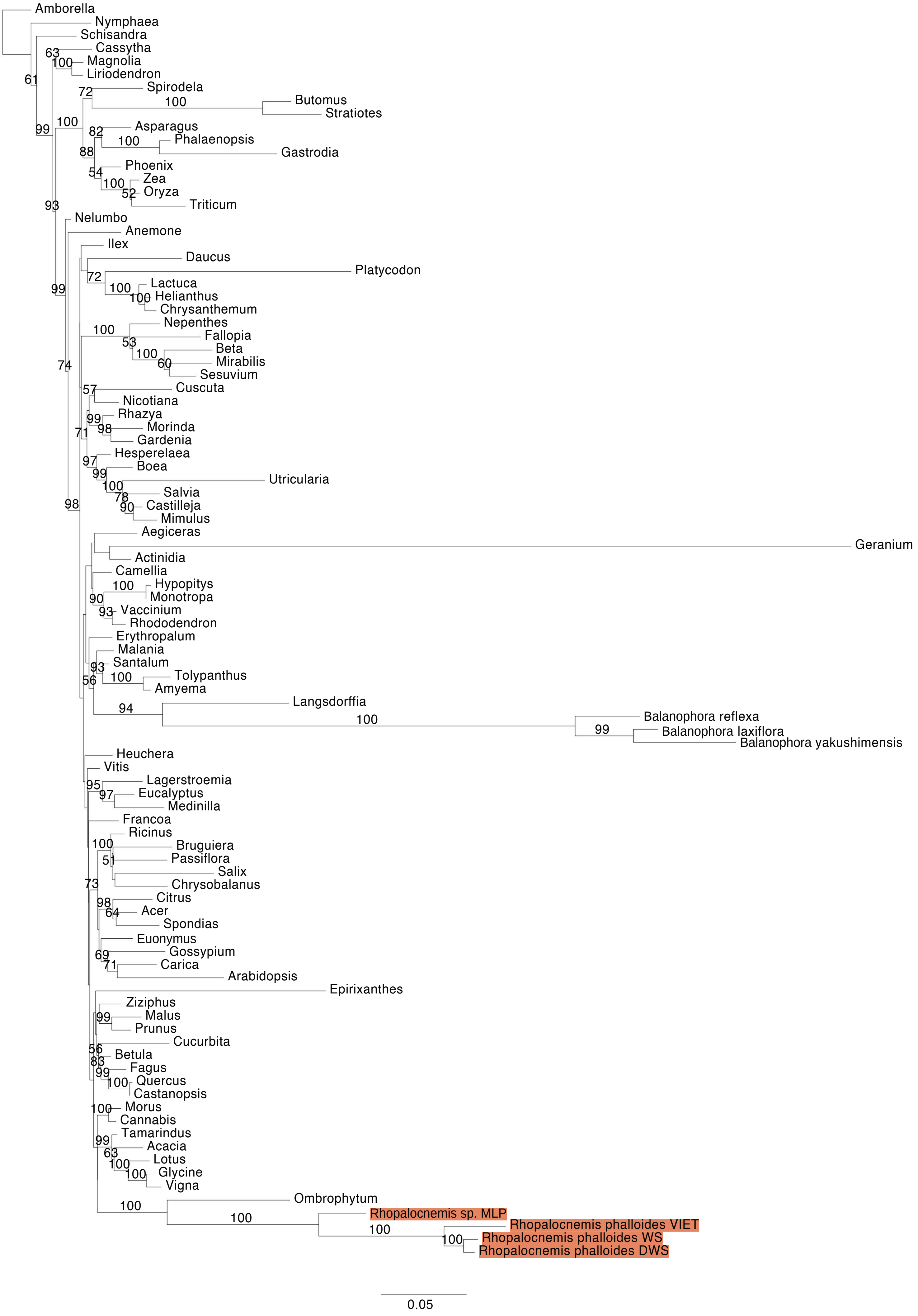


**Figure S6.** Maximum Likelihood (ML) phylogenetic analyses of protein-coding gene sequences in the mtDNAs of *Rhopalocnemis*. ML bootstrap support values ≥50% are shown. Scale bars correspond to substitutions per site.

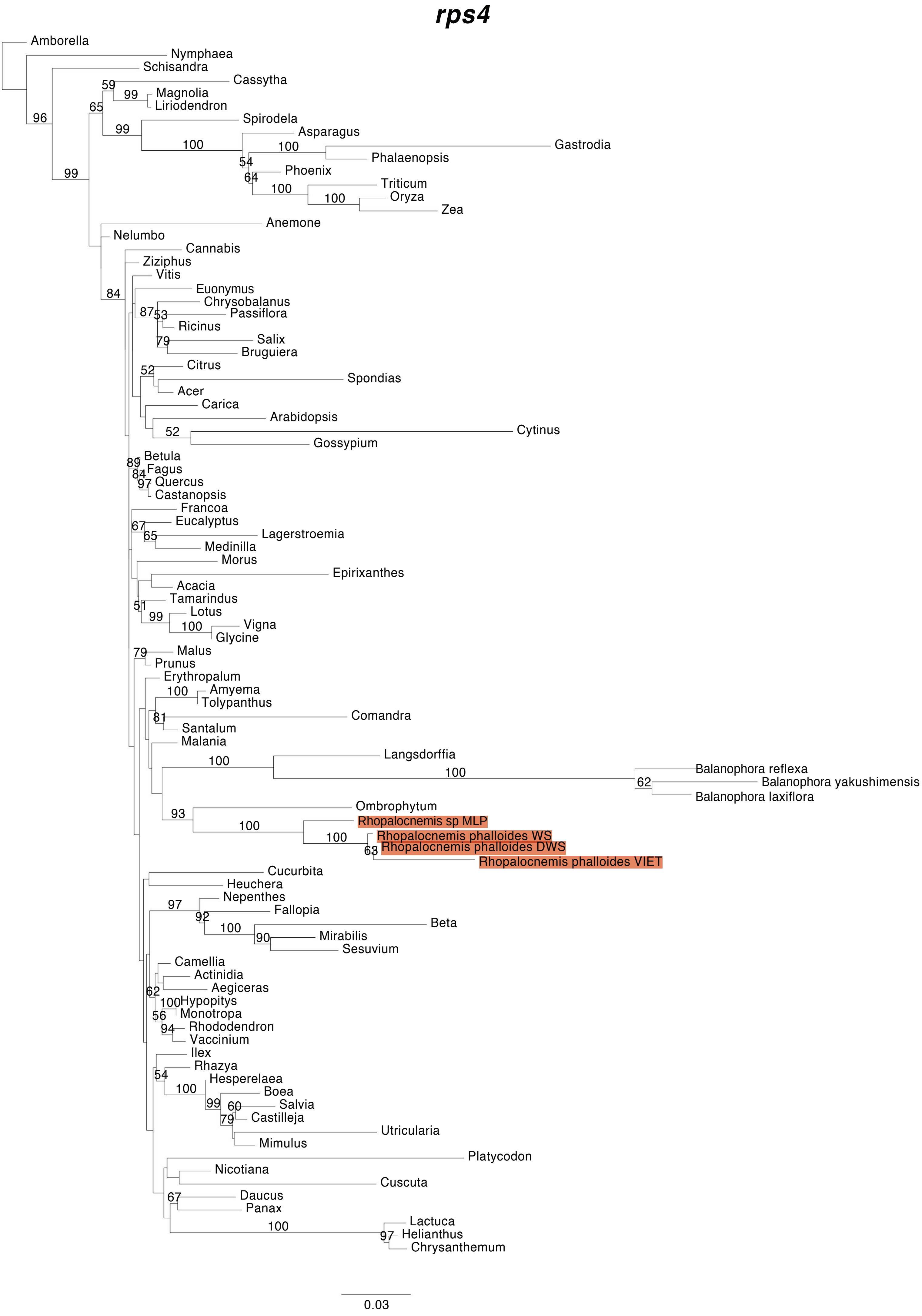


**Figure S6.** Maximum Likelihood (ML) phylogenetic analyses of protein-coding gene sequences in the mtDNAs of *Rhopalocnemis*. ML bootstrap support values ≥50% are shown. Scale bars correspond to substitutions per site.

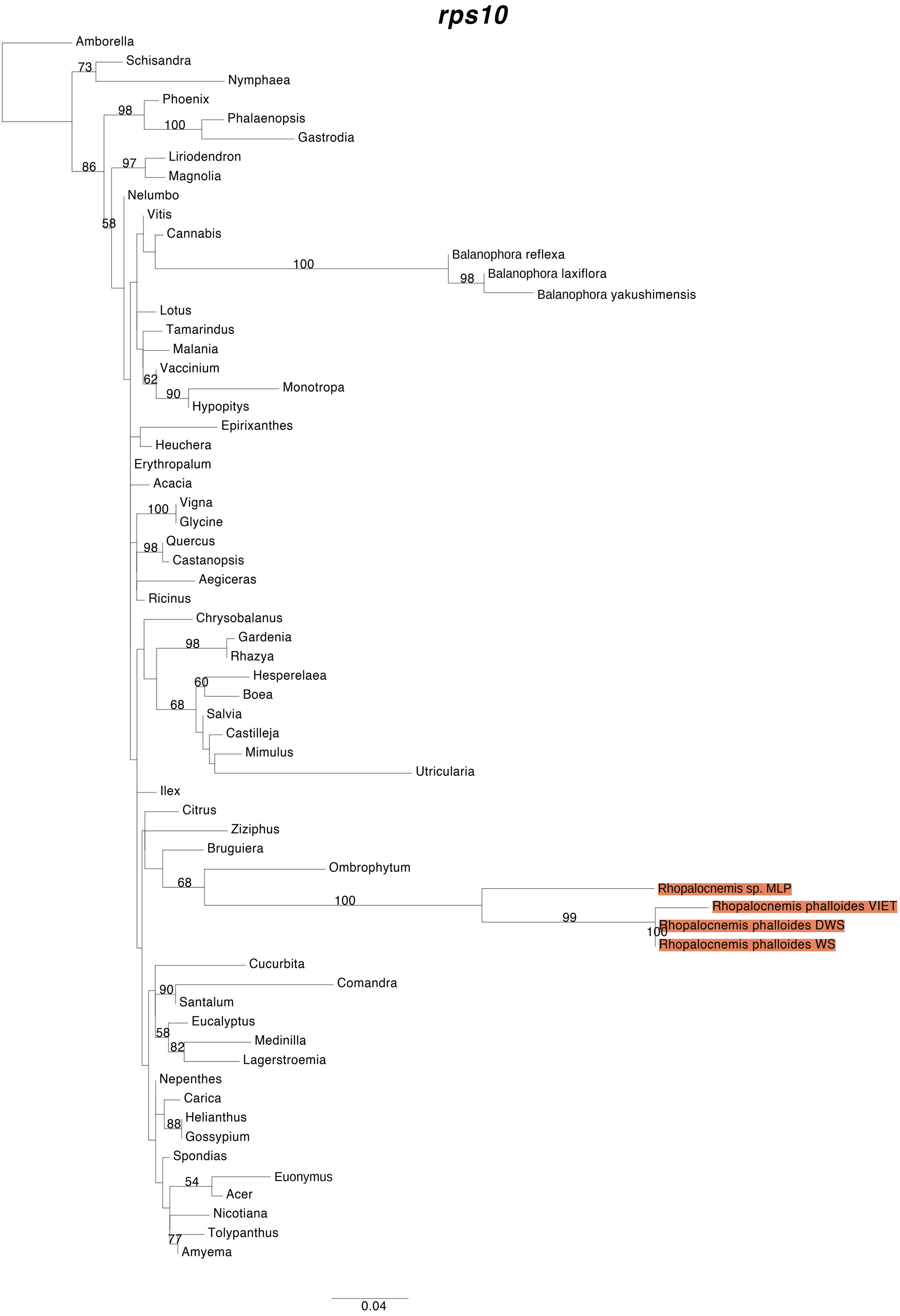


**Figure S6.** Maximum Likelihood (ML) phylogenetic analyses of protein-coding gene sequences in the mtDNAs of *Rhopalocnemis*. ML bootstrap support values ≥50% are shown. Scale bars correspond to substitutions per site.

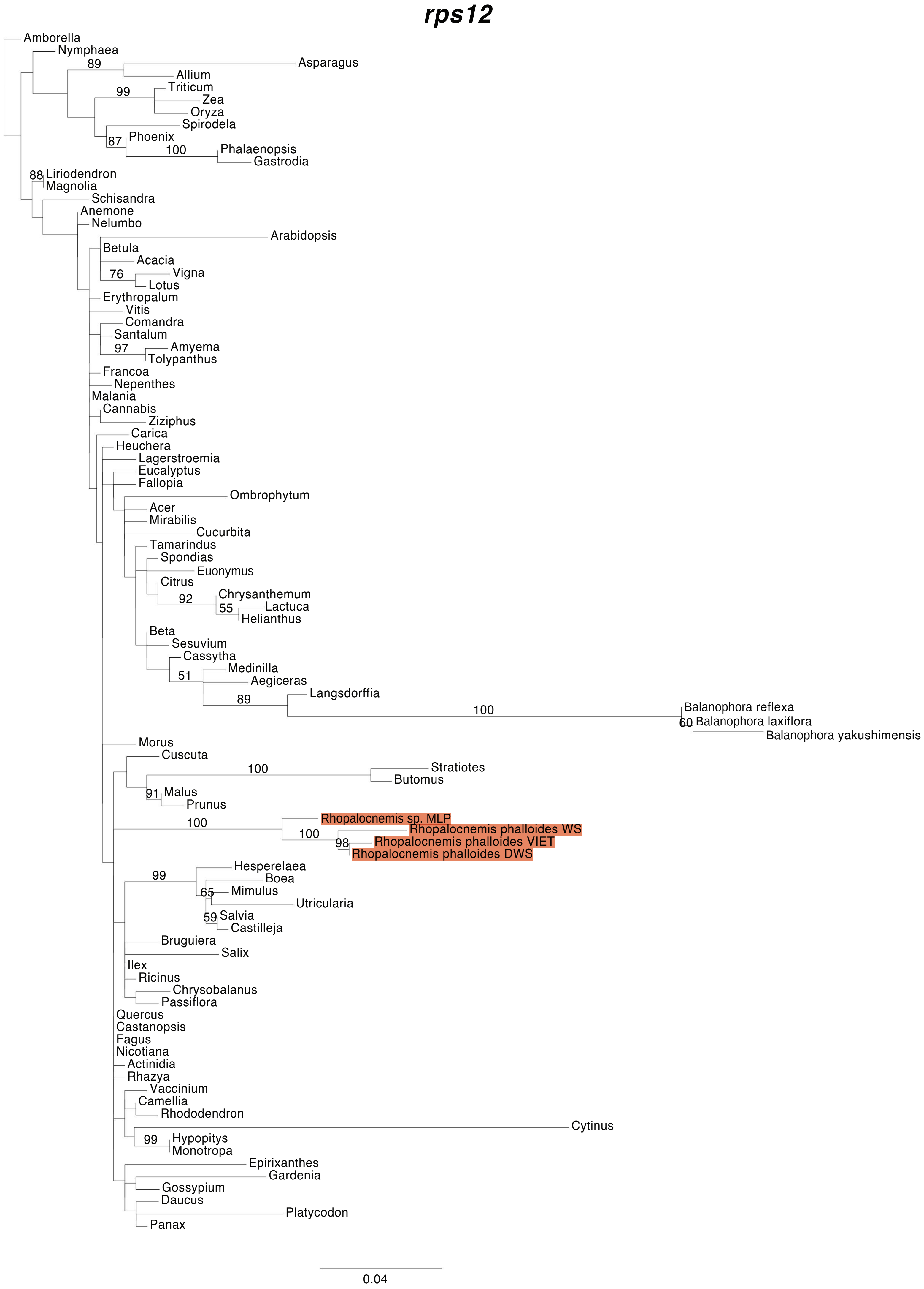


**Figure S6.** Maximum Likelihood (ML) phylogenetic analyses of protein-coding gene sequences in the mtDNAs of *Rhopalocnemis*. ML bootstrap support values ≥50% are shown. Scale bars correspond to substitutions per site.

### *rps19*

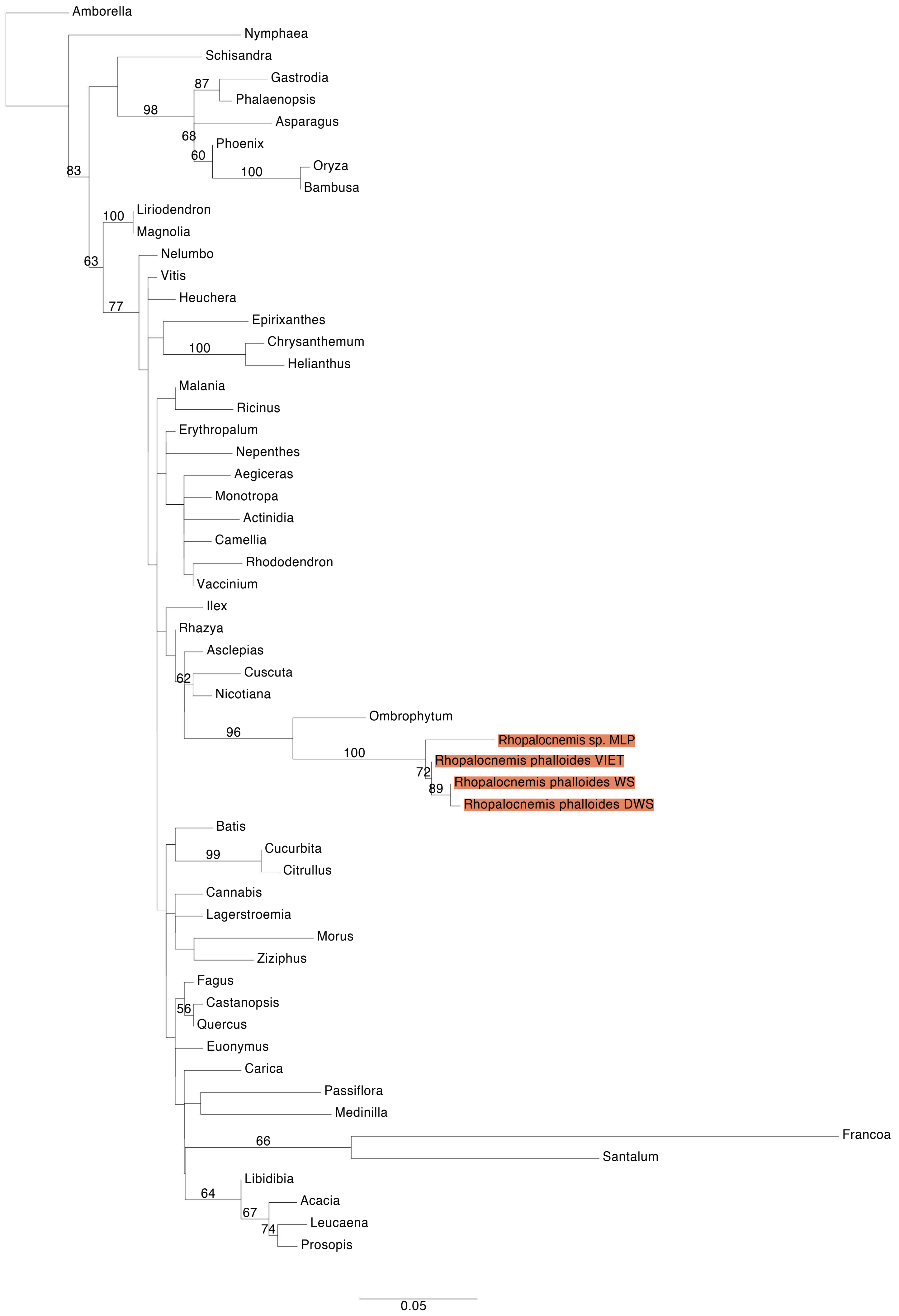


**Figure S6.** Maximum Likelihood (ML) phylogenetic analyses of protein-coding gene sequences in the mtDNAs of *Rhopalocnemis*. ML bootstrap support values ≥50% are shown. Scale bars correspond to substitutions per site.

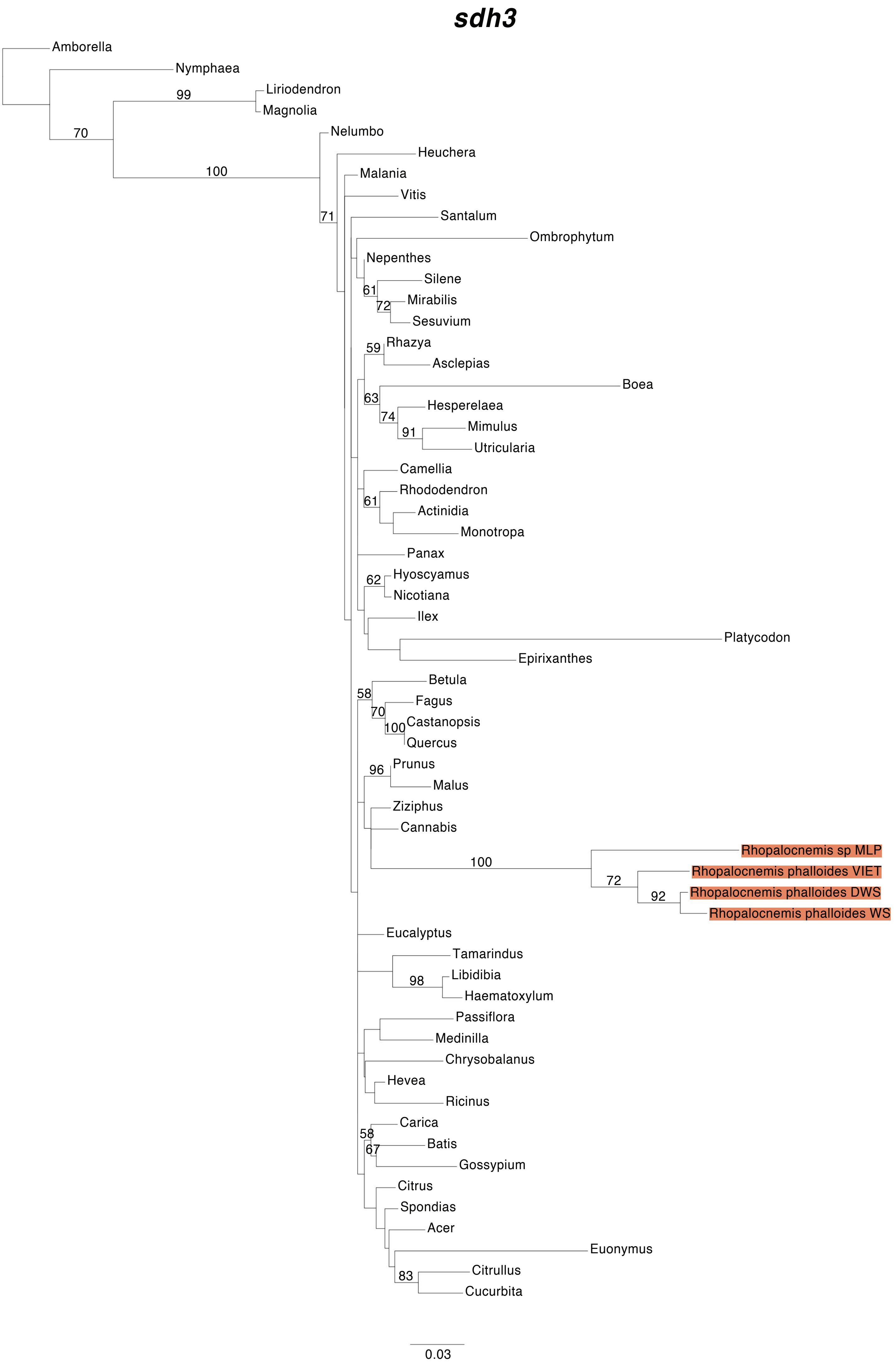


**Figure S6.** Maximum Likelihood (ML) phylogenetic analyses of protein-coding gene sequences in the mtDNAs of *Rhopalocnemis*. ML bootstrap support values ≥50% are shown. Scale bars correspond to substitutions per site.

#### *sdh 4*

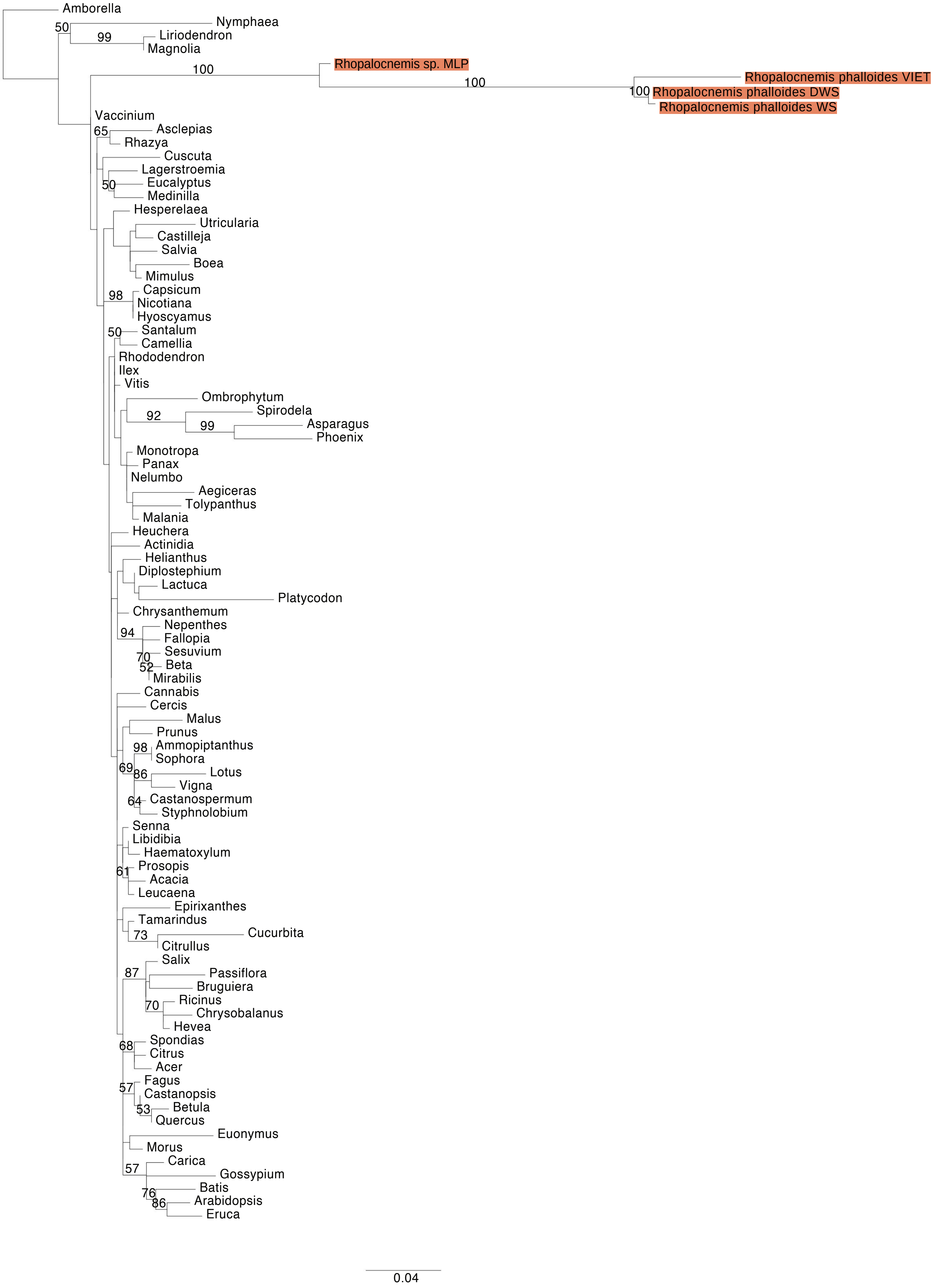


**Figure S6.** Maximum Likelihood (ML) phylogenetic analyses of protein-coding gene sequences in the mtDNAs of *Rhopalocnemis*. ML bootstrap support values ≥50% are shown. Scale bars correspond to substitutions per site.

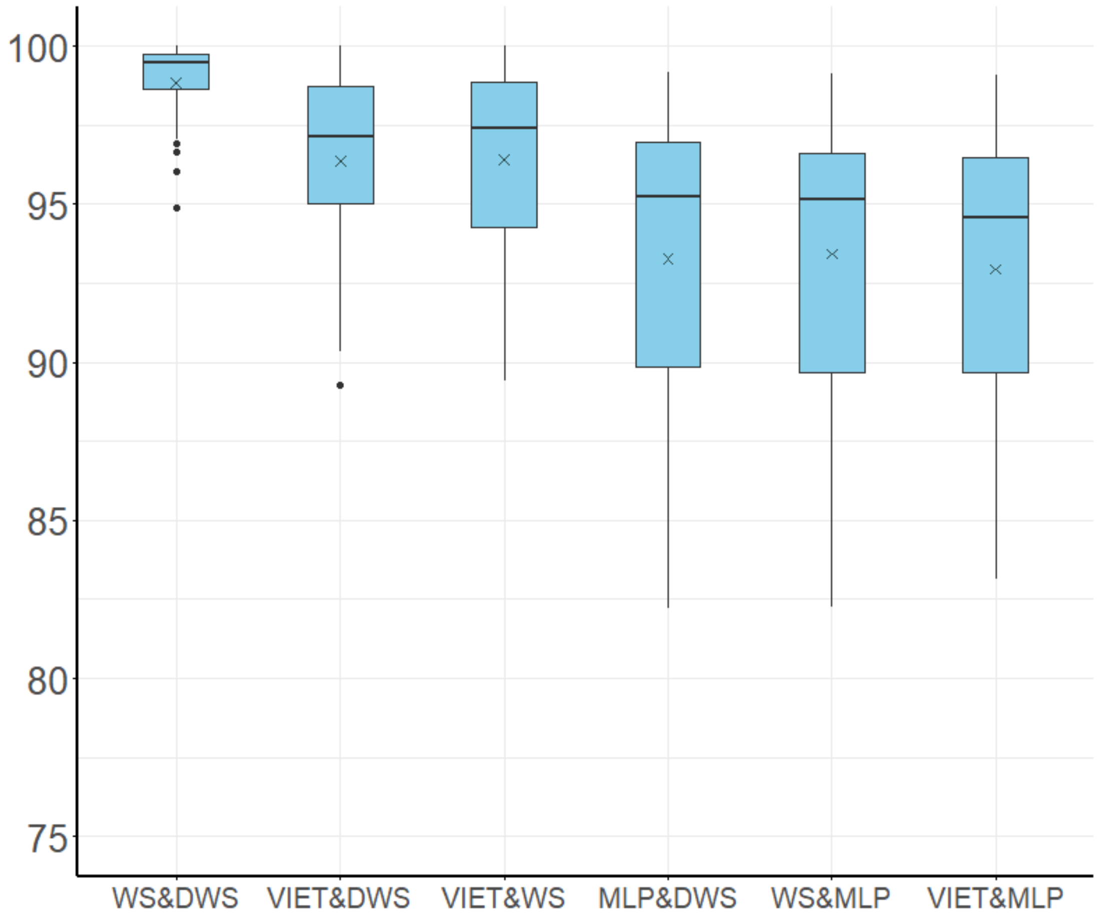


**Figure S7**. Pairwise sequence identity of 36 mitochondrial protein-coding genes among four individuals of *Rhopalocnemis* (WS, DWS, VIET and MLP). The vertical ordinate shows sequence identity (%). The boxes correspond to the first and third quartiles of sequence identity of 36 genes. The horizontal line and cross symbol inside the boxes correspond to median and mean of sequence identity of 36 genes. The whiskers extend to the smallest and largest values within 1.5-fold the interquartile range, respectively. Data that fall outside of the range of the whiskers are marked with dots.

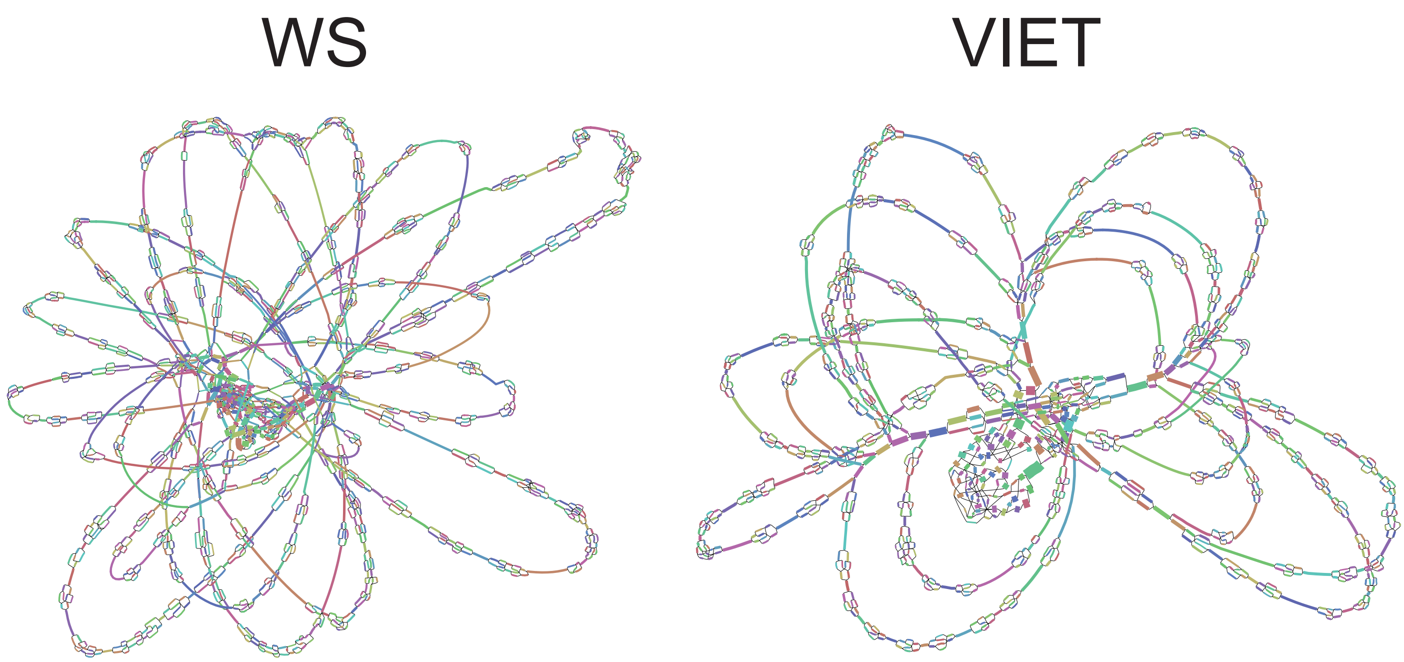


**Figure S8**. The mitogenome assembly graph of two individuals of *Rhopalocnemis phalloides* (WS and VIET) visualized in Bandage based on the raw GFA file produced by GetOrganelle. The contigs colors were displayed randomly.

OBS1/OSB2-4/OSBX

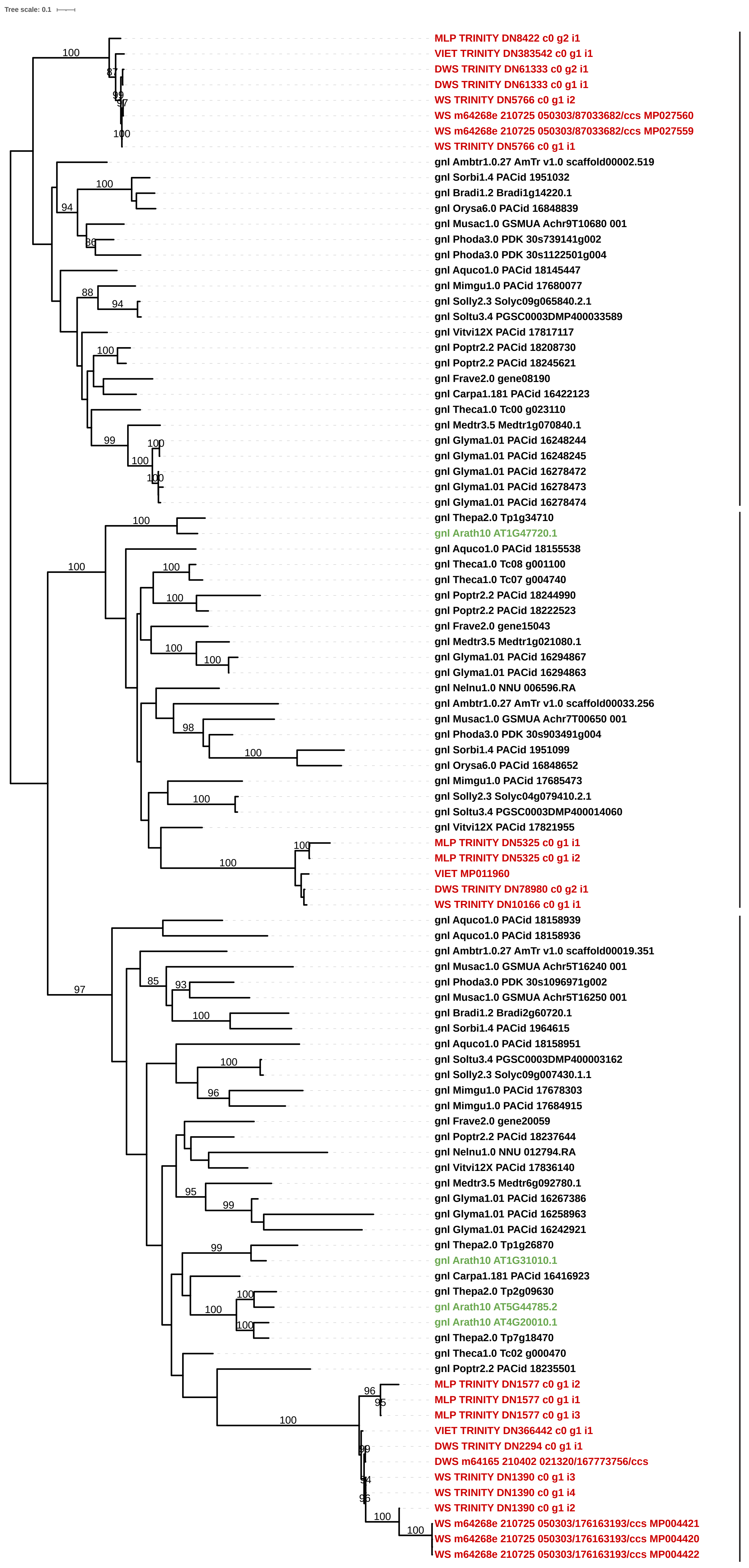


RECA2/RECA3

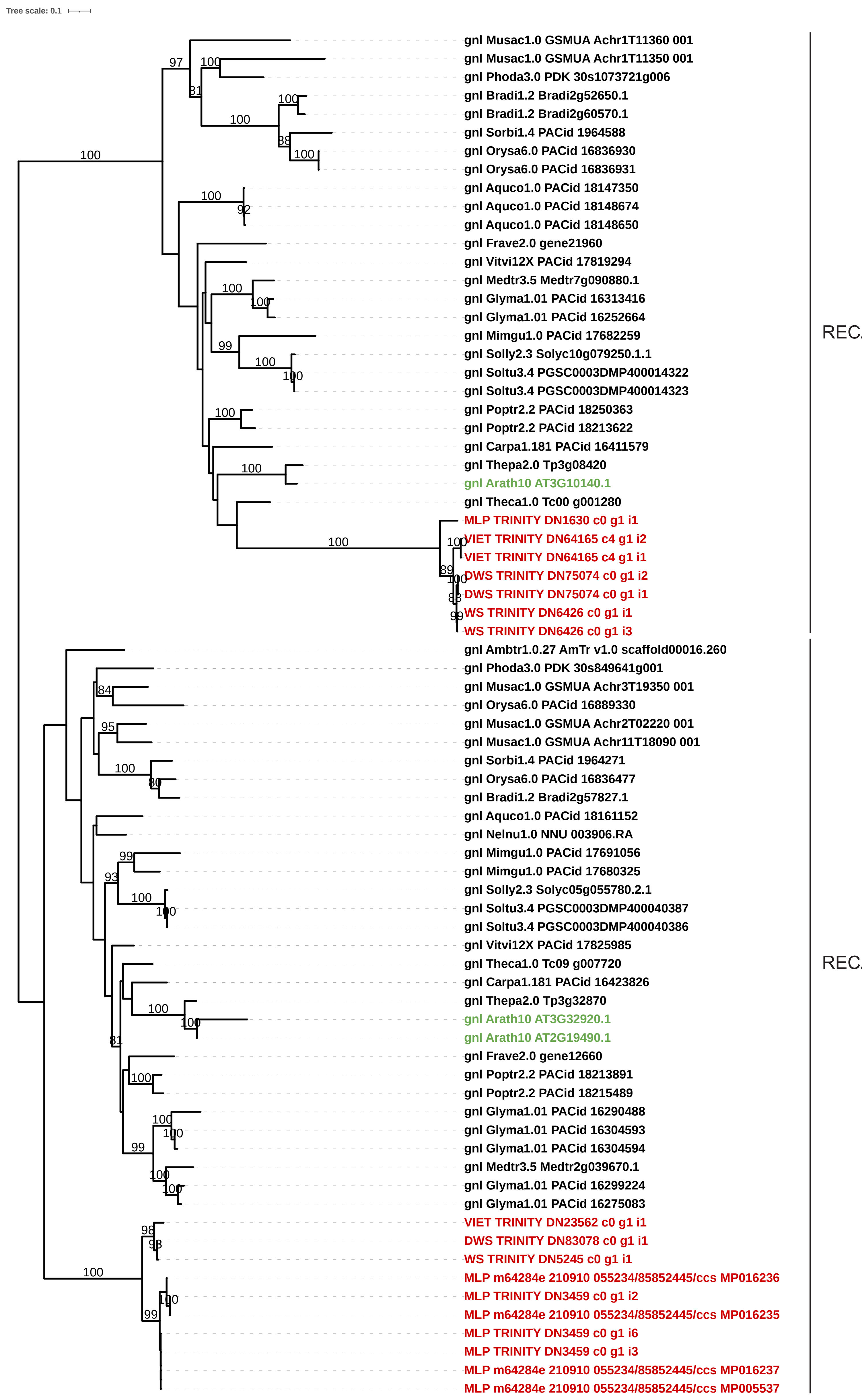


**Figure S9**. Maximum likelihood trees of two gene families that have experienced recent duplication events in the lineage leading to *Arabidopsis*. Genes from *Rhopalocnemis* and *Arabidopsis* are shown in red and green,respectively. *OSBX* is a homolog of other *OSB genes* that is present across angiosperms but absent in *Arabidopsis* (Ceriotti *et al*. 2022).
